# Destabilization of intratumor Tregs by CTLA-4 engagement confers anti-CTLA-4-driven immunotherapy

**DOI:** 10.64898/2026.07.28.741328

**Authors:** Na Liu, Junji Xu, Wai Lim Ku, Weiwei Chen, Yaqiang Cao, Rida Kazmi, Wenwen Jin, Thierry Gauthier, Shunqun Luo, Song Shen, Liliana C. Patiño, Yun-Ji Lim, Valentina Ottaviani, Emily Naylor, Keji Zhao, WanJun Chen

## Abstract

Immune checkpoint inhibitors (ICI) that are antibodies against CTLA-4 have achieved therapeutic effects on multiple types of cancers ^1–4^, but the mechanisms underlying the therapy remain incompletely understood. In contrast to the initial theory that the antibody blockades CTLA-4 on T cells is of central importance, it has recently been demonstrated that selective reduction of CD4^+^Foxp3^+^ regulatory T cells (Tregs) in the tumor microenvironment by the antibody plays a key role in the anti-tumor effects^5–8^, although this phenomenon remains under debate in human patients^9–12^. We show here that anti-CTLA-4 antibody engages CTLA-4 in Tregs specifically in the tumor tissues to reduce their stability and survival and weaken their suppressive function through upregulating TGF-β signaling. This leads to the antibody-mediated - upregulation of CD4^+^ and CD8^+^ effector T cell anti-tumor immunity and consequently cancer immunotherapy. Specifically, anti-CTLA-4 antibody directly stimulates CTLA-4 in intratumor Tregs to enhance their TGF-β receptor I (TβRI)-mediated TGF-β signaling^13^. This results in reduction of IL-2 receptor CD25 expression and downregulation of lactate metabolism in intratumor Tregs to reduce their stability and survival ^14–16^, and also compromises their suppressive function by inhibiting Foxp3 expression. This activity requires high levels of CTLA-4 expression in the intratumor Tregs and the presence of antibody Fc receptors in the tumor tissues. Strikingly, deletion of TβRI specifically in Tregs completely prevents the reduction of intratumor Tregs and abrogates the anti-CTLA-4-mediated cancer immunotherapy. In contrast to intratumor Tregs, anti-CTLA-4 antibody treatment blocks CTLA-4 in the intratumor CD4^+^ Foxp3^−^ and CD8^+^ T effector cells and in the peripheral Tregs to decrease their TβRI expression and increase their expansion due to their relatively low levels of CTLA-4 in the same tumor bearing mice. Significantly, the engagement of CTLA-4 by anti-human CTLA-4 antibody (ipilimumab) also upregulates *TGFBR1* but decreases *Il2RA* expression and lactate metabolism in human Tregs *in vitro* and in humanized mice *in vivo*, leading to suppression of tumor progression. We have provided additional mechanism underlying anti-CTLA-4-mediated anti-tumor effects through destabilizing intratumor Tregs by CTLA-4 engagement-mediated TGF-β signaling. This could lay a theoretical foundation for designing optimal immunotherapy based on anti-CTLA-4 antibody in cancer patients.

## Introduction

Cytotoxic T lymphocyte-associated protein 4 (CTLA-4) is a key negative regulatory molecule in T cell immune responses. It is constitutively expressed in CD4^+^Foxp3^+^ regulatory T cells (Tregs), but also induced for expression in activated CD4^+^ and CD8^+^ T cells ^17–21^. As the first FDA-approved immune checkpoint inhibitor antibody, anti-CTLA-4 therapy has achieved promising clinical effects on some cancers ^22–24^. However, the underlying cellular and molecular mechanisms by which anti-CTLA-4 antibody executes its therapeutic function remain incompletely understood. It was initially suggested that anti-CTLA-4 antibody achieved its anti-tumor effects by blocking CTLA-4 on effector T cells and consequently releasing the brake for T cell activation and anti-tumor activities ^25,26^. However, subsequent studies have revealed that its anti-tumor effects are primarily ascribed to the selective reduction of the intratumor Tregs ^27–29^. Although it is still debated whether the anti-CTLA-4 antibody-mediated tumor Treg decrease also explains its anti-tumor function in human cancer patients ^10,30,31^, this discrepancy might be due to the complexity of human Tregs and/or the time points and location of the Tregs examined following antibody therapy. For instance, human Tregs can be classified into several different subsets based on their phenotype and functionality ^32,33^, so FOXP3 expression alone might not be sufficient to identify the true effector Tregs that are affected by the anti-CTLA-4 antibody in the tumor tissues. Nevertheless, although it is known that anti-CTLA-4 antibody mediated reduction of tumor Tregs serves as a major mechanism to confer the antibody therapeutic effects in preclinical models, the detailed mechanisms by which the antibody decreases the tumor Tregs remain largely unclear. The abnormally high levels of CTLA-4 expression in intratumor Tregs may be the molecular basis for the unique anti-CTLA-4-mediated Treg reduction in the tumor tissues. It has been suggested that the CTLA-4 antibody kills intratumor Tregs through antibody-dependent cellular cytotoxicity (ADCC) and/or antibody-dependent cellular phagocytosis (ADCP) ^5, 7,8,29,34^. However, whether this antibody-mediated killing is the sole mechanism for the tumor Treg reduction remains unknown. In addition, it is unclear whether the anti-CTLA-4 treatment also affects the suppressive activity of Tregs in the tumor tissue. Furthermore, it remains to be discovered whether the effects on intratumor T effector cells and peripheral Tregs and mechanism for those effects are the same as those of intratumor Tregs.

In our study, we discovered unexpectedly that the therapeutic anti-CTLA-4 antibody selectively engages the CTLA-4 on intratumor Tregs to inhibit their stability/survival and compromise their function through upregulating TGF-β receptor I (TβRI)-mediated TGF-β signaling. The downregulation of IL-2 receptor α CD25 and the decrease in lactate metabolism in the tumor Tregs play key roles in their instability and reduced survival; the decreased Foxp3 expression accounts greatly for the compromised suppressive activity of the intratumor Tregs. We elucidated the underlying molecular mechanism that CTLA-4 stimulation activates the Rho-GTPase-P38 MAPK signaling pathway to up-regulate TGF-β signaling in Tregs. Remarkably, we proved that the deletion of *Tgfbr1* specifically in Tregs completely abolished the instability and the reduction of tumor Tregs induced by CTLA-4 antibody treatment, resulting in a complete failure to control tumor progression. Importantly, the engagement of CTLA-4 by anti-human CTLA-4 antibody (ipilimumab) also upregulates *TGFBR1* but decreases *Il2RA* expression and lactate metabolism in human Tregs *in vitro* and in humanized mice *in vivo*, leading to suppression of tumor progression. We have thus revealed a previously unrecognized mechanism underlying anti-CTLA-4-mediated anti-tumor effects.

## Results

### Therapeutic CTLA-4 antibody increases TGF-β signaling in Tregs in tumor tissues

To understand the mechanisms of therapeutic anti-CTLA-4 antibody relating to tumor rejection, we treated tumor-bearing mice with anti-CTLA4 antibodies (9H10, Syrian hamster IgG; 9D9, mouse IgG2b), both of which have been reported to significantly delay tumor growth in preclinical mouse models ^5,35,36^. Foxp3*^EGFP^* mice (mice with transgenic expression of GFP in Foxp3 protein) were inoculated with MCA-205 sarcoma tumor cells and then treated with a single dose of one of the indicated anti-CTLA-4 antibodies or their respective isotype control antibodies, when the tumors were palpable at days 6-7 (Figure S1A). As expected ^5,29^, even a single dose of anti-CTLA-4 treatment suppressed tumor growth and almost eliminated Tregs in the tumor tissues about one week later (day 13) (Figures S1B-S1D). We observed that another anti-CTLA-4 antibody 9D9 (mouse IgG2b) also led to significant suppression of tumor growth and reduction of intratumor Tregs (Figure S1E). We used 9H10 antibody for the subsequent studies unless otherwise indicated. As there were almost no detectable Tregs in the tumor tissues after one week of 9H10 antibody treatment, which precluded analyzing the Tregs for mechanistic studies, we sacrificed tumor mice at day three after anti-CTLA-4 treatment for a thorough analysis. Strikingly, even three days after a single dose of antibody treatment, there was already a significant reduction of tumor growth in the 9H10-treated mice when compared to the isotype control antibody-treated ones (Figure 1A). We thus chose day 3 after the treatment to analyze the immune cells in the tumor tissues to investigate the mechanisms of anti-CTLA-4 treatment. Analysis of immune cells (CD45^+^) in the tumor tissues revealed that the Tregs already exhibited significant reduction of both the frequency and absolute number in the tumor microenvironment at this time (Figure 1B) after anti-CTLA4 treatment compared with control antibody-treated groups. On the other hand, the IFN-γ^+^TNF-α^+^ cells in both CD4^+^ Foxp3^−^ non-Treg responder and CD8^+^ T effector cells were significantly upregulated in the mice that received anti-CTLA-4 antibody treatment (Figure 1C). Similar findings were obtained in other tumor models of mice with colon cancer MC38 treated with anti-CTLA-4 (Figures S1F-S1I). The data collectively indicate that anti-CTLA-4 antibody treatment suppresses tumor growth, and this is accompanied by a reduction of intratumor Tregs, suggesting a role of intratumor Tregs in the therapeutic effects.

**Figure. 1.**
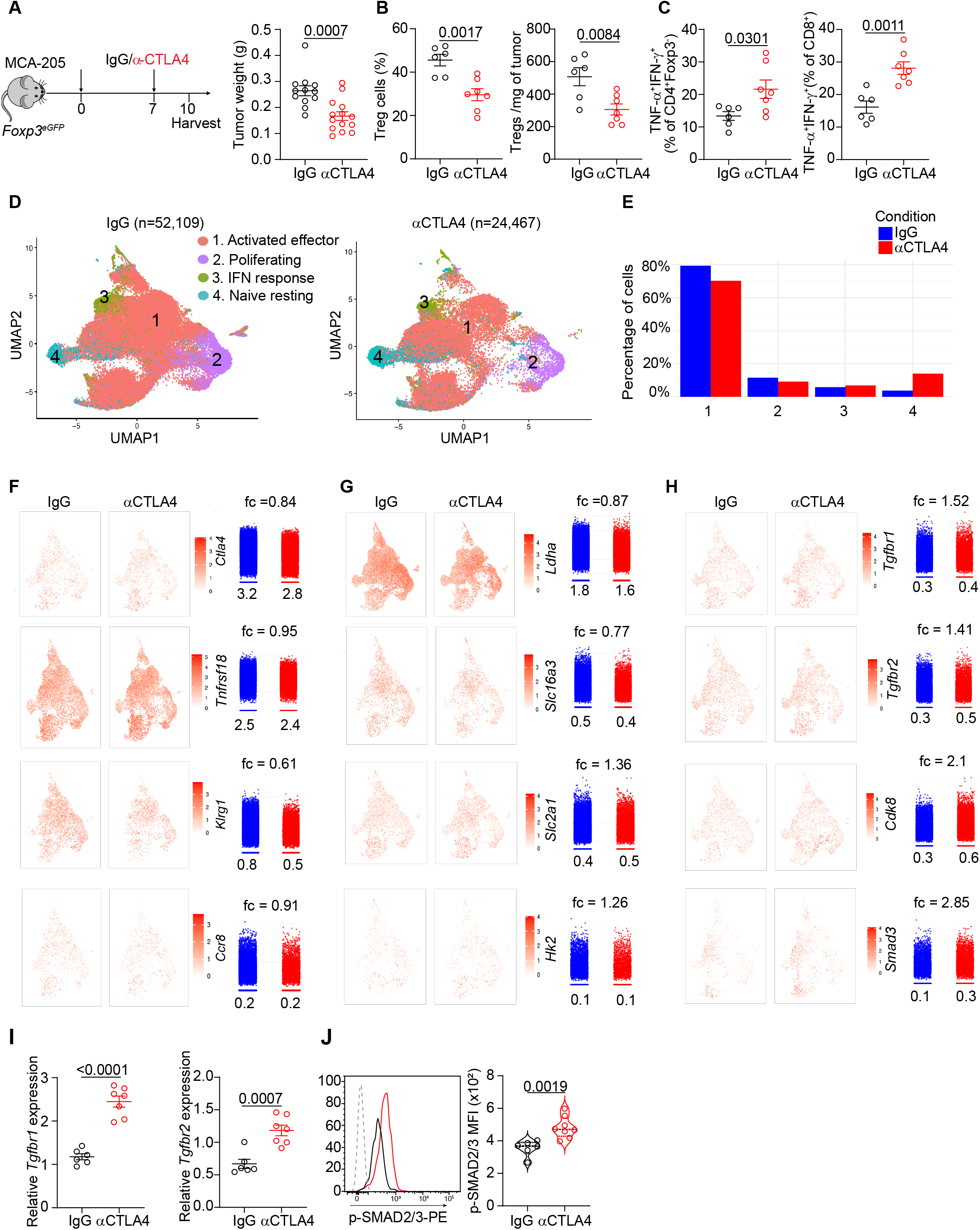
Upregulation of TGF-β signaling in intratumor Tregs induced by anti-CTLA-4 treatment. (A) Left, Diagram showing the experimental procedure, and right, tumor weight as indicated. (B) Left, Percentage of Tregs (CD4^+^Foxp3^+^) among total CD4^+^ T cells from tumors and right, Tregs number per mg of tumor in mice bearing Sarcoma MCA-205 cells. (C) Percentage of TNF-α^+^ IFN-γ^+^ producing CD4^+^Fxop3^-^ cells or CD8^+^ T cells in tumors between IgG vs. αCTLA-4 treatments. (D) UMAP plot of scRNA-seq data encompassing GFP^+^ Tregs from IgG or anti-CTLA-4 treatment. (E) Quantification of cluster designation for Tregs between IgG vs. αCTLA-4 treated mice. (F-H) The expression of canonical Treg subset genes, lactate and glycolysis metabolism and TGF-β signaling-associated genes were visualized in red scale using UMAP projection. (I) Expression of *Tgfbr1* and *Tgfbr2* mRNA in intratumor Tregs by RT-PCR from IgG group or αCTLA-4 group. (J) Representative histogram of p-SMAD3 in Tregs from tumors in IgG vs. αCTLA-4 treatments. Data in (A-C, I-J) are representative of three independent experiments with similar results. Significance was determined by unpaired two-tailed *t*-test (A-C, I-J). Data shown as mean ± s.e.m.

To provide indisputable evidence that intratumor Tregs are required for anti-CTLA-4-mediated tumor suppression, we next depleted Tregs with DT (diphtheria toxin) treatment of Foxp3*^DTR^* mice in the MCA-205 tumor models followed by a single dose of anti-CTLA-4 treatment for 3 days (Figure S1J). As expected, anti-CTLA-4 antibody treatment resulted in significant reduction of tumor weight with a dramatic decrease in Tregs in the tumor tissues (Figures S1K and S1L). Depletion of Tregs alone significantly decreased the tumor weight as expected (Figure S1K). However, anti-CTLA-4 antibody treatment failed to reduce further tumor growth compared to antibody control treatment in these Treg-depleted mice (Figure S1K), confirming a key role of Tregs as the main operator for anti-CTLA-4 antibody antitumor effects. It should be noticed that Tregs are depleted systemically, thus the potential effects of Tregs depletion in periphery organs cannot be completely excluded.

We next investigated the underlying mechanisms by which anti-CTLA-4 induced a rapid reduction of Tregs in the tumor microenvironment with an unbiased and global view using scRNA-seq analysis. We harvested the tumors 3 days after a single dose of antibody treatment and sorted the Tregs with flow cytometry to perform scRNA-seq analysis on the Tregs from the groups treated with anti-CTLA-4 (24,467 Tregs) and control IgG (52,109 Tregs). This large number of individual tumors Tregs transcriptomes enabled us to cluster them with relatively high resolution. Globally, we compared the differentially genes (DEGs) in whole tumor-infiltrating Tregs between IgG and anti-CTLA-4 treatment. We found that the expression of genes related to Tregs effector status (*Tnfrs18, Ccr8, Ctla4*, *Nr4a2, Klrg1*, *Il10, Lag3,* and *Nkg7*) was significantly decreased in intratumor Tregs after anti-CTLA-4 treatment (Figures S2A and S2B). As expected, Tregs were reduced in the tumors with anti-CTLA-4 therapy, and 4 distinct subclusters of intratumor Tregs were identified (Figure 1D). The majority of tumor Tregs manifested an activated phenotype (cluster 1), identified based on enriched expression of *Il2ra*, *Lag3*, *Cxcr6*, *Tnfrsf4*, *Tnfrsf18*, *Tnfrsf9*, *Icos*, *Klrg1,* and *Ccr8* ^37–39^, which are broadly associated with Treg activation (Figures 1D, 1E and S2C). Murine intratumor Tregs also contained proliferating phenotype (cluster 2) including *Stmn1*, *Mki67*, *Tubb5* and *Top2a,* which were slightly decreased with therapeutic anti-CTLA-4 antibody treatment (Figures 1D, 1E and S2C). The frequency of other distinct cluster sharing type I IFN-responsive signatures (cluster 3) enriched for IFN-inducible genes such as *Ifit3*, *Oasl2*, *Ifitm2*, *Xaf1* and *Irf7*, was retained following anti-CTLA-4 treatment (Figures 1D, 1E and S2C). In contrast, there were one cluster sharing a naïve/resting signature (cluster 4) ^40,41^ (*Klf2*, *Sell*, *Tcf7*, *Ccr7* and *S1pr1*) expressed in the tumor Tregs, and their frequencies were upregulated with anti-CTLA-4 antibody treatment (Figures 1D, 1E and S2C). Strikingly, cell proportion analysis identified that enriched representation of activated clusters was significantly reduced in the tumors after anti-CTLA-4 administration (Figures 1D and 1E). In addition, activated cluster 1 was dominantly enriched in the tumor Tregs and expressed high levels of *Ctla4*, *Tnfrsf18*, *Klrg1* and *Ccr8,* which were associated with tumor Treg function and were downregulated in anti-CTLA-4 treated tumors (Figure 1F). As it is reported that tumor Tregs are dependent on lactate uptake instead of glucose consumption to maintain their stability and suppressive function ^35,42,43^, we found that in this activated cluster those genes encoding lactate dehydrogenase (*Ldha)* and monocarboxylate transporter Mct4 (*Slc16a3*) were dramatically decreased in the tumor Tregs treated with anti-CTLA-4 antibody (Figure 1G). Corresponding to this, the genes involving glucose glycolysis such as: *Glut1* (*Slc2a1*) and *Hk2* were up-regulated in tumor Tregs upon anti-CTLA-4 treatment (Figure 1G).

Our scRNA-seq analysis revealed that naïve state gene signatures were enhanced in intratumor Tregs from anti-CTLA-4-treated mice, and our previous findings indicated that TGF-β signaling plays a vital role determining T cell quiescence and activation ^13^. We next asked whether TGF-β signaling in tumor Tregs was affected by anti-CTLA-4 antibody treatment. Surprisingly, we found an increase in TGF-β signaling, characterized by enhanced gene expression of *Tgfbr1*, *Tgfbr2*, *Cdk8,* and *Smad3*, in intratumor Tregs from the anti-CTLA-4-treated group (Figure S2B). Besides, a comprehensive TGF-β signature score which was based on a curated list of 84 core pathway genes was also increased in each Treg subsets (Figure S2B). This result was confirmed by quantitative polymerase chain reaction (qPCR) analysis demonstrating that intratumor Tregs indeed expressed significantly higher levels of *Tgfbr1* and *Tgfbr2* in tumor mice treated with the same anti-CTLA-4 (9H10) (Figure 1I). Consistently, a significantly enhanced phosphorylation of Smad-2/3 (pSmad2/3), a key mediator for TGF-β signaling ^44^, was detected in the tumor Tregs after anti-CTLA-4 treatment (Figure 1J). Even more surprisingly, these TGF-β signaling associated genes were also dramatically upregulated in the activated cluster 1 in response to anti-CTLA-4 treatment (Figure 1H). This upregulation of TGF-β signaling was also reproduced in the intratumor Tregs by another anti-CTLA-4 antibody (9D9) treatment compared with Tregs with its isotype IgG2b control antibody treatment (Figure S2D). Strikingly, even at day one after 9H10 treatment in the tumor mice, the *Tgfbr1* expression in the tumor Tregs, which was accompanying reduction of frequency and absolute number of Tregs in the tumors, was already significantly upregulated (Figures S2E and S2F). To further confirm that the upregulation of *Tgfbr1* expression in the intratumor Tregs is attributed to anti-CTLA-4 antibody rather than due to the contribution from consequent enrichment of less activated and/or quiescent Tregs following anti-CTLA-4 treatment, we harvested tumor-bearing mice after anti-CTLA-4 injection for 15 hours. At this time point, tumor size, as well as the frequency and number of Tregs in the tumors in anti-CTLA-4 treated mice, exhibited no any reduction compared to IgG-treated controls (Figures S2G and S2I). However, the *Tgfbr1* expression already significantly up-regulated in tumor-infiltrating Tregs after anti-CTLA-4 treatment (Figure S2J). Strikingly, we also obtained the similar results after anti-CTLA-4 treatment for only 4 hours (Figures S2K-S2M ). The data collectively reveal that the intratumor Tregs play an essential role for mediating the therapeutic effects of anti-CTLA-4 antibody on tumor development, and the antibody treatment results in a reduction of intratumor Tregs that is associated with an increase in TGF-β signaling.

### CTLA-4 engagement upregulates TGF-β signaling in Tregs

We next sought to understand the underlying molecular mechanisms by which anti-CTLA-4 antibody treatment enhances TGF-β signaling in Tregs in the tumor tissues. As anti-CTLA-4 treatment was originally thought of as immune checkpoint blockade (ICB), we initially hypothesized that CTLA-4 blockade would increase TGF-β signaling in Tregs. As TGF-β receptor I (TβRI) is the essential receptor and authoritative marker for TGF-β signaling in T cells ^13^, we used *Tgfbr1* expression to measure TGF-β signaling for the subsequent experiments unless otherwise indicated. To test this, murine CD4^+^CD25^+^Foxp3^+^ Tregs were isolated from spleens and LNs of *Foxp3^EGFP^* mice and stimulated with suboptimal doses of anti-CD3 (0.01μg/ml) and CD28 (1 µg/ml) antibodies (since a high dose of TCR stimulation completely inhibited the *Tgfbr1* expression ^13^), in the presence of soluble CTLA-4 Fab fragments (4F10-Fab’) ^18^ to block CTLA-4. We used IL-2 receptor CD25 as a marker for Treg cell activation and showed that blockade of CTLA-4 significantly increased the expression of CD25 (Figure 2A). Surprisingly, however, blockade of CTLA-4 with CTLA-4-(Fab’) significantly decreased TGF-β receptor I (*Tgfbr1*) expression in Tregs (Figure 2B). Notably, in these Treg cell cultures without antigen-presenting cells (APCs), we observed that Tregs express low but detectable levels of CTLA-4 ligands B7-1 and B7-2, suggesting it might provide the costimulatory signaling in an autocrine and/or paracrine manner in the context of CTLA-4 blockade (Figure S2N). The data suggests that the upregulation of TGF-β signaling in intratumor Tregs induced by anti-CTLA-4 administration is independent of CTLA-4 blockade.

**Figure. 2.**
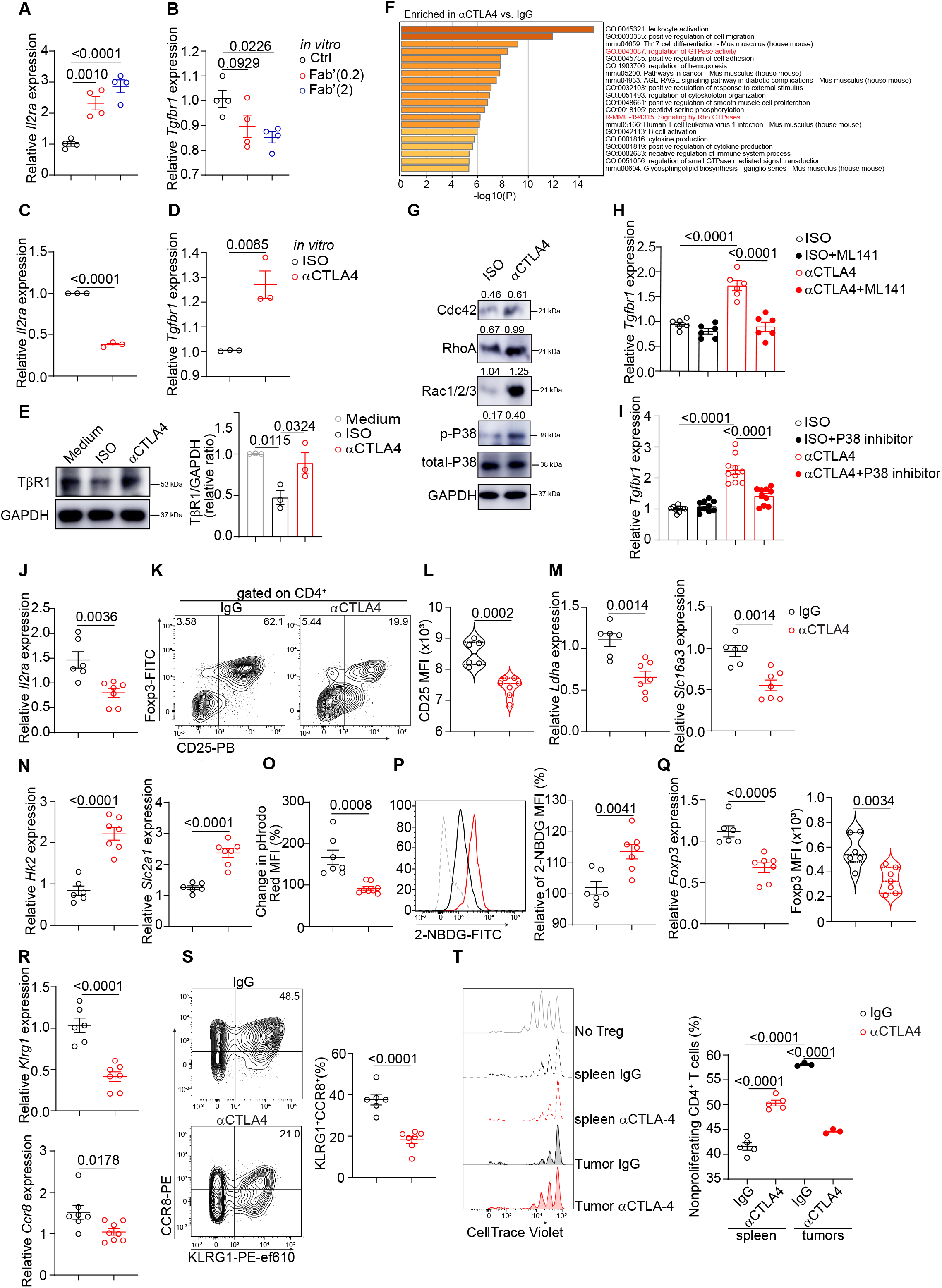
CTLA-4 engagement induces an instability and functional deficiency in intratumor Tregs through upregulation of TGF-β signaling. (A-B) Expression of *Il2ra* and *Tgfbr1* mRNA in Tregs cultured with different doses of CTLA-4 blocking antibody (4F10-Fab’) in the presence of a low dose of αCD3 (0.01µg/ml) plus αCD28 (1µg/ml) overnight. (C-D) RT-PCR analysis of *Il2ra* and *Tgfbr1* mRNA in Tregs cultured with 10 (µg/ml) of isotype or intact anti-CTLA-4 antibody cross-linked with corresponding secondary antibody, and in the presence with αCD3/CD28 coactivator beads for overnight. (E) Protein level of TβR1 in Tregs cultured under CTLA-4 engagement overnight. (F) Bar chart of clustered enrichment ontology categories was performed by Metascape on all (differentially expressed genes) DEGs in Tregs from scRNA-seq dataset between IgG vs. αCTLA-4 treatment. (G) Abundance of the indicated proteins in Tregs presented in IgG treatment or CTLA-4 engagement. (H-I) RT-PCR analysis of *Tgfbr1* expression in Tregs cultured with cross-linked IgG or anti-CTLA-4, in the presence of Rho-GTPase inhibitor (ML141, 5 µM) or P38 inhibitor (SB203580, 2.5 µM) for overnight. (J) RT-PCR analysis of *Il2ra* in intratumor Tregs from IgG vs. αCTLA4 treatment. (K) Flow cytometry of Foxp3 and CD25 gated from tumor-infiltrating CD4^+^ T cells between IgG vs. αCTLA4 groups. (L) Analysis of CD25 expression in intratumor Tregs between IgG vs. αCTLA4 treatment. (M-N) Expression of genes related to lactate metabolism and glucose metabolism in isolated intratumor Tregs after IgG vs. αCTLA4 treatment. (O) Lactate uptake reflected by relative change of pHrodo Red fluorescence in isolated intratumor Tregs. (P) Glucose uptake reflected by 2-NBDG fluorescence in isolated intratumor Tregs. (Q) Analysis of Foxp3 expression from intratumor Tregs between IgG vs. αCTLA4 injections at both mRNA and protein levels. (R-S) Analysis of KLRG1 and CCR8 expression in tumor-infiltrating Tregs between IgG vs. αCTLA4 treatment at both mRNA and protein levels. (T) Left, representative histogram showing the proliferation of CD4^+^CD25^-^ T cells (Tcon, CD45.1^+^) after 72h of co-culture with Tregs (CD45.2^+^) sorted from IgG or αCTLA-4 treated tumor-bearing mice. Right, ability of Tregs to suppress the proliferation of Tcon cells (at a 1:4 ration of Treg: Tcon cells) labeled with CellTrace Violet (CTV). Data in (A-D) were pooled from two or three independent experiments with similar results. Data in (E, G, H-T) are representative of three independent experiments. Significance was determined by one-way analysis of variance (ANOVA) with Tukey’s multiple comparison (A-B, E, H-I, T) or unpaired two-tailed *t*-test (C-D, J-S). Data shown as mean ± s.e.m.

Considering that CTLA-4 is acting as a co-inhibitory molecule that is constitutively expressed in a high level in intratumor Tregs ^29^, and these Tregs display a compromised stability and reduced number after anti-CTLA-4 treatment ^35^, we next postulated that the anti-CTLA-4 antibody might stimulate CTLA-4 on the tumor Tregs to upregulate TGF-β signaling. For this, we cultured Tregs with anti-CD3 and CD28 antibodies in the presence of intact anti-CTLA-4 antibody (9H10) followed by secondary goat anti-hamster cross-linking antibody to mimic CTLA-4 engagement ^45,46^. Remarkably, we found that the engagement of CTLA-4 on Tregs downregulated CD25 expression (Figure 2C) and significantly upregulated their *Tgfbr1* expression (Figure 2D) compared to isotype control antibody-treated Tregs. Immunoblotting analysis confirmed the upregulation of TβRI protein in Tregs after CTLA-4 engagement (Figure 2E) ^13^. This CTLA-4 engagement did not increase cell death of Tregs in vitro (data not shown). Thus, the engagement rather than the blockade of CTLA-4 upregulates TGF-β signaling in Tregs.

To understand the underlying molecular pathways involved in the upregulation of TGF-β signaling in Tregs upon CTLA-4 engagement, we first performed gene set enrichment analysis from activated cluster in the scRNA-seq data set and found that genes in the categories “regulation of GTPase activity” and “signaling by Rho GTPase” were markedly increased in the tumor Tregs after anti-CTLA-4 administration (Figure 2F). As Rho GTPase signaling was reported to play a crucial role in triggering multiple immune functions ^47^, we then validated this enhanced Rho-GTPase signaling in Tregs at both RNA and protein levels upon CTLA-4 engagement *in vitro* (Figures 2G and S2O). Accordingly, CTLA-4 engagement upregulated phosphorylated p-P38 MAPK, but not NF-κb, JNK or ERK signaling, all of which are downstream pathways of Rho-GTPase signaling (Figure 2G, and data not shown). Importantly, inhibition of Rho-GTPase with the specific inhibitor ML141 reduced *Tgfbr1* expression induced by CTLA-4 stimulation (Figure 2H). Consistently, inhibition of P38 MAPK with its specific inhibitor also abrogated the upregulation of *Tgfbr1* expression in Tregs driven by CTLA-4 engagement (Figure 2I). We also observed a similar reduction in *Tgfbr1* expression with other Rho-GTPase inhibitors (Casin and Ketorolac) or with P38 inhibitor (SB202190) in presence of CTLA-4 engagement (Figure S2P). Thus, the Rho-GTPase-p38 signaling pathway is involved in the upregulation of TGF-β signaling induced by CTLA-4 engagement.

### Engagement of CTLA-4 by antibody compromised the stability and function of intratumor Tregs

The current dominant theory believes that anti-CTLA-4 antibody treatment mediated reduction of Tregs in the tumor tissues is through direct killing by ADCC ^5,27,29^. While it is possible that ADCC-mediated Treg death might account for some degree of the Treg reduction in the tumor tissues, we reasoned that alternatively the CTLA-4 antibody through CTLA-4 signaling alternatively compromises the stability and survival leading to the reduction of Tregs, and/or inhibits the suppressive ability of Tregs, and these effects result, perhaps synergistically, in the enhancement of antitumor immunity. To address this, we analyzed the expression of genes related to Treg stability/survival and their suppressive function with different approaches. We first determined that the decrease in intratumor Tregs was not due to the inhibition of cell proliferative ability measured by Ki67 staining in anti-CTLA-4 antibody-treated tumor mice (Figure S3A). As IL-2 signaling through its receptor CD25 in Tregs is one of the most important pathways to maintain the stability and survival of Tregs ^14,16,48–50^, we next demonstrated with semi**-**quantitative RT-PCR analysis that anti-CTLA-4 treatment indeed significantly decreased the *Il2r* mRNA in the intratumor Tregs (Figure 2J). Flow cytometry analysis further validated the suppression of both CD25^+^ Treg frequency and CD25 protein levels (measured as mean fluorescence intensity, MFI) in the intratumor Tregs upon anti-CTLA-4 antibody therapy (Figures 2K and 2L). Strikingly, just one day after 9H10 treatment, the CD25 mRNA and protein was already significantly decreased in the tumor Tregs compared to control antibody treated mice (Figure S3B). This decrease in CD25 expression was attributed to CTLA-4 signaling, as engagement of CTLA-4 by the cross-linking antibody (Figure 2C), but not blockade of the molecule (Figure 2A), suppressed CD25 expression in culture. The data indicate that CD25-mediated IL-2 signaling is compromised in Tregs by CTLA-4 signaling in the tumor tissues treated with therapeutic CTLA-4 antibody.

To confirm that the downregulation of CD25-mediated IL-2 signaling plays a role in the instability and survival of intratumor Tregs upon anti-CTLA-4 therapy, we next administered exogenous IL-2 (6000 IU/0.1 ml) into the tumor mice before CTLA-4 antibody treatment (Figure S3C) in the hope of restoring the IL-2 signaling in the Tregs. We found that exogenous administration of IL-2 abolished the therapeutic effects of anti-CTLA-4 antibody on the tumors (Figure S3D). Analysis of the intratumor T cells revealed that exogenous IL-2 increased the frequency and absolute number of intratumor Tregs (Figure S3E). Anti-CTLA-4 antibody therapy did not significantly change the proliferation of intratumor Tregs measured by Ki-67 staining, and proliferation was not altered by the administration of exogenous IL-2 treatment (Figure S3F). IL-2 treatment also did not reverse the downregulation of Foxp3 expression induced by anti-CTLA-4 antibody (Figure S3G), suggesting that the restoration of Treg cells was primarily due to the protection of Treg survival. Indeed, the frequency of CD25^+^ Tregs and the expression levels of CD25 in Tregs were significantly upregulated in tumors treated with anti-CTLA-4 plus IL-2 (Figure S3H), suggesting increased IL-2 signaling in the Tregs. This was confirmed by CTLA-4 engagement with cross-linking antibody in TCR-stimulated Tregs *in vitro*, in which IL-2 significantly restored CD25 expression suppressed by CTLA-4 stimulation (Figure S3I). IL-2 treatment failed to increase the total number of CD4^+^Foxp3^−^ and CD8^+^ T cells in the tumor tissues (Figure S3J). Consequently, IL-2 treatment abolished anti-CTLA-4 antibody-mediated upregulation of IFN-γ^+^TNF-α^+^ in CD8^+^ T cells although not significantly in CD4^+^Foxp3^−^ T cells (Figure S3K). The data collectively indicate that IL-2 treatment restored the number of intratumor Tregs decreased by anti-CTLA-4 treatment, and the IL-2 treatment compromises antitumor effects by the antibody. This confirms that the downregulation of CD25 expression plays an important part in the anti-CTLA-4 mediated reduction of Tregs, and consequently in the therapeutic effects on tumor.

In addition to IL-2 signaling, lactate metabolism has been reported to be a key pathway for maintaining the stability of tumor infiltrating Tregs ^35,42,43^, whereas glycolysis does the opposite for intratumor Tregs ^35,42^. To investigate this possibility, we focused on changes in genes critical for lactate metabolism and glycolysis in tumor Tregs affected by anti-CTLA-4 treatment. Our scRNA-Seq analysis of tumor Tregs in mice treated with anti-CTLA-4 antibody revealed downregulation of lactate metabolism pathways, as evidenced by the reduction of *Ldha* and *Slc16a3* expression (Figure 1G), but upregulation of *Hk2* and *Slc2a1* gene expression, which are critical components in glycolysis (Figure 1G). We confirmed the downregulation of *Ldha* and *Slc16a3*, and the upregulation of *Hk2* and *Slc2a1* genes in tumor Tregs upon anti-CTLA-4 treatment by quantitative RT-PCR analysis (Figures 2M and 2N). As reported ^42^, the use of lactate by intratumor Tregs was significantly higher than Tregs in spleen and lymph nodes (Figure S4A). Importantly, the uptake of lactate by intratumor Tregs was significantly suppressed by anti-CTLA-4 treatment (Figure 2O). This downregulation of *Ldha* and *Slc16a3* in intratumor Tregs was already significant just one day after anti-CTLA-4 (9H10) therapy (Figure S4B). As previous reports indicated glucose avidity in Tregs leading to their instability and correlating with their poor suppressive capacity ^42^, we also evaluated glucose uptake by intratumor Tregs using the fluorescent glucose tracer 2-NBDG. We showed that intratumor Tregs significantly increased glucose uptake after treatment with anti-CTLA-4 compared those treated with the isotypic control antibody (Figure 2P). We confirmed that engagement of CTLA-4 by cross-linking antibody led to the suppression of *Ldha* and *Slc16a3* in culture (Figure S4C). The data collectively indicate that the reduced ability to respond to IL-2 and the reduced lactate metabolism in the tumor Tregs result in the instability and reduction of tumor Tregs.

In addition to its effects on stability and survival, anti-CTLA-4 treatment also compromised the suppressive ability of Tregs in the tumor tissues. Levels of Foxp3 expression in Tregs are the primary and key factor determining the function of Tregs ^51,52^. We found that anti-CTLA-4 therapy significantly decreased the mRNA and protein expression of Foxp3 in the tumor Tregs (Figure 2Q). We validated that CTLA-4 signaling reduced Foxp3 expression in Tregs by utilizing cross-linking anti-CTLA-4 antibody stimulation *in vitro* (Figure S4D). In addition, Chemokine receptor 8 (CCR8) and Killer cell lectin-like receptor subfamily G member 1 (KLRG1) are highly expressed in intratumor Tregs ^6,53^ and represent key markers associated with Treg-mediated suppression of anti-tumor immunity in tumor tissues ^53,54^. As expected, we found that anti-CTLA-4 antibody administration also resulted in decrease in the expression of *Ccr8* and *Klgr1* in our scRNA-seq analysis (Figure 1F), and this was confirmed by quantitative RT-PCR and flow cytometry (Figures 2R and 2S). Furthermore, we confirmed that CTLA-4 engagement inhibited the expression of *Klrg1* and *Ccr8* (Figure S4E). Strikingly, the decrease in *Foxp3*, *Ccr8* and *Klgr1* in intratumor Tregs was already apparent just one day after anti-CTLA-4 treatment in tumor mice (Figure S4F). The data suggests a compromised suppressive function of intratumor Tregs treated by therapeutic anti-CTLA-4 antibody.

To directly investigate whether the suppressive function of tumor Tregs was affected by CTLA-4 antibody administration, we isolated Tregs from tumor tissues and spleens in the tumor-bearing *Foxp3^EGFP^*mice 3 days after anti-CTLA-4 or control antibody treatment, and cocultured them with normal CD4^+^Foxp3^−^ responder T cells to perform the standard suppression assay ^55^. We demonstrated that the tumor Tregs exhibited more potent suppressive activity than did the splenic Tregs from the control antibody treated mice (Figure 2T). However, tumor Tregs isolated from CTLA-4 antibody-treated mice exhibited significantly reduced suppressive function compared to tumor Tregs from control antibody-treated mice (Figure 2T). This set of functional data matched well with the changes of the aforementioned functional molecules upon CTLA-4 antibody therapy, establishing that CTLA-4 antibody therapy compromised the suppressive function of tumor Tregs besides inducing loss of stability and reduction in numbers. The decrease in the stability/number and function together confers the anti-tumor immune response against tumor growth induced by anti-CTLA-4 antibody.

### TGF-β signaling is required for CTLA-4 engagement-mediated instability of Tregs *in vitro*

As the engagement rather than blockade of CTLA-4 induced upregulation of TGF-β signaling in the tumor Tregs, we next determined that TGF-β signaling was required for the instability and functional deficiency of Tregs caused by CTLA-4 therapy. We employed two different approaches to block TGF-β signaling in Tregs in culture. First, we blocked TβRⅠ (ALK5) activity with its specific inhibitor (SB-431542) in wild-type Tregs and showed that the TβRⅠ inhibitor reversed the downregulation of *Il2ra, Ldha* and *Slc16a3, Ccr8,* and *Klrg1* expressions in Tregs induced by CTLA-4 engagement in cultures (Figures S5A-S5C). As expected, the TβRI inhibitor also blocked its own upregulation induced by CTLA-4 engagement (Figure S5D), suggesting a positive autocrine effects of TGF-β signaling ^13^. Importantly, engagement of CTLA-4 by cross-linking antibody significantly downregulated Foxp3 expression in Tregs (Figures S3I and S5E), and inhibition of TβRI activity with its inhibitor restored the expression of Foxp3 (Figure S5E).

*Second,* we stimulated CTLA-4 in Tregs with *Tgfbr1* deficiency (*Tgfbr1f/f-Foxp3^eGFP-Cre-ERT^*^2^ mice treated with tamoxifen for 5 days) (Figure S5F) with cross-linking anti-CTLA-4 antibody and showed that indeed the CTLA-4 signaling completely failed to downregulate expression of the aforementioned genes, namely CD25, *Ldha* and *Slc16a3*, *Ccr8,* and *Klrg1,* in these knockout Tregs (Figures S5G-S5I). Importantly, cross-linking CTLA-4 in the ko Tregs failed to reduce the downregulation of Foxp3 expression to the same extent as the wild-type Tregs (Figure S5J). These data suggest that the upregulated TGF-β signaling is directly responsible for the instability and functional deficiency induced by CTLA-4 engagement by the therapeutic antibody in intratumor Tregs.

### Anti-CTLA-4 fails to upregulate TGF-β signaling, reduce intratumor Tregs and suppress tumor growth in experimental melanoma

Unlike the tumor models of MCA-205 and MC-38, it has been known that anti-CTLA-4 therapy alone fails to suppress tumor growth in the experimental melanoma models ^56^, but the underlying mechanisms remain largely unknown. We thus hypothesized that the anti-CTLA-4 antibody might be unable to upregulate TGF-β signaling in the intratumor Tregs, thus fail to reduce their survival and function. To this end, melanoma tumor cells (B16-F10) were subcutaneously injected into *Foxp3^EGFP^*mice, followed by the same anti-CTLA4 antibody (9H10) treatment for 3 days (Figure S6A). Consistent with the published data ^56^, anti-CTLA-4 immunotherapy had no beneficial and therapeutic effects in the B16-F10 melanoma model (Figure S6A). Analysis of the T cells in the tumor tissues revealed no any reduction of the frequency and cell number of intratumor Tregs (Figures S6B and S6C). Consistent with this, neither the expression of CD25, nor the molecules associated with lactate metabolism were reduced in response to anti-CTLA-4 injection (Figures S6D-S6F). The levels of Foxp3 and other functional markers related to Treg suppressive activity, including KLRG1 and CCR8 was also unaffected after anti-CTLA-4 treatment (Figures S6G and S6H). Notably, tumor-infiltrating Tregs exhibited no upregulation of TGF-β signaling, as indicated by *Tgfbr1* and *Tgfbr2* expression (Figures S6I). These altogether indicate a failure of CTLA-4 engagement-mediated increase in TGF-β signaling and consequent reduction of intratumoral Tregs survival and stability in the B16-F10 melanoma. Consequently, the IFN-γ^+^TNF-α^+^ CD4^+^Foxp3^-^ and CD8^+^ T cells in the tumor were not enhanced by the anti-CTLA-4 treatment (Figures S6J). To explore whether the short treatment duration accounted for the ineffective response, we extended the anti-CTLA-4 treatment for 6 days using a single dose of anti-CTLA-4 antibody. However, same results were obtained (Figures S6K-S6T). These findings indicate that in B16-F10 melanoma model, anti-CTLA-4 was unable to actively engage Tregs to up-regulated TGF-β signaling to induce Tregs instability. As a result, the inhibitory brake imposed by Tregs to reinvigorate effector was not released, thereby failing to promote anti-tumor immunity. Although the underlying mechanisms responsible for the failure of anti-CTLA-4 engagement to CTLA-4 in intratumor Tregs in this melanoma model remain elusive, the data provides further evidence to support that successful tumor control upon anti-CTLA-4 treatment manifests with a dysfunctional Treg signature, defined as upregulation of TGF-β signaling which was induced by CTLA-4 engagement in intratumor Tregs.

### Deletion of TGF-β signaling in Tregs abolishes anti-CTLA-4 antibody mediated anti-tumor effects

To validate the significance of upregulated TGF-β signaling induced by CTLA-4 engagement in intratumor Tregs upon anti-CTLA-4 antibody therapy in tumor development *in vivo*, we utilized two independent mouse models to block TGF-β signaling specifically in Tregs. In the first approach, we sorted *Tgfbr1^-/-^* and wild-type Tregs from *Tgfbr1^f/f/^ER-cre* mice (CD45.2) that had been treated with tamoxifen or oil for 5 days, respectively, and co-transferred them with naïve CD4^+^ and CD8^+^ T cells from CD45.1 mice into *Rag1^-/-^*mice, followed by tumor cell (MCA-205) injection and anti-CTLA-4 or control antibody treatment as outlined in (Figure 3A). We determined that *Tgfbr1* is deleted in the Tregs and the purity of the sorted Tregs was more than 90% (Figures S7A and S7B ). In this adoptive transfer model, we observed that the recipient mice transferred with *Tgfbr1^-/-^* Tregs completely lost the anti-tumor benefit caused by anti-CTLA-4 administration, whereas those mice that received wild-type Tregs exhibited marked suppression of tumor weight as expected (Figure 3B). Strikingly, analysis of intratumor Tregs (gating strategy in Figures S7C) revealed that deletion of *Tgfbr1* completely prevented the reduction of intratumor Tregs and the downregulation of Foxp3 expression induced by anti-CTLA-4 antibody therapy (Figures 3C-3E). Further analysis showed that anti-CTLA-4 antibody treatment-mediated suppression of CD25 expression, as well as the frequency of CD25^+^ Tregs in the tumor tissues were abolished in the *Tgfbr1^-/-^* tumor Tregs (Figure 3F). In addition, the downregulations of *Ldha* and *Slc16a3* observed in wild-type intratumor Tregs upon anti-CTLA-4 treatment were also restored in the ko Tregs (Figure 3G). Moreover, the suppression of *Foxp3*, *Ccr8*, and *Klrg1* expression in the wild-type intratumor Tregs induced by CTLA-4 antibody treatment was also completely reversed in the *Tgfbr1^-/-^* Tregs in the tumor tissues (Figures 3H, S7D and S7E), suggesting a reservation of the suppressive activity in the intratumor ko Tregs in response to CTLA-4 antibody treatment. Accordingly, anti-CTLA-4 antibody therapy completely lost its beneficial enhancement of IFN-γ^+^TNF-α^+^ in CD4^+^ Foxp3^−^ and CD8^+^ T effector cells in the tumor tissue in the mice co-transferred with *Tgfbr1^-/-^* Tregs (Figure 3I). Notably, we observed that the CTLA-4 expression in intratumor *Tgfbr1*^-/-^Tregs was not reduced compared to WT Tregs (Figures 37F and S8B). We also observed that the expression of CTLA-4 in *Tgfbr1^-/-^* intratumor Tregs was reduced to the same extent as the wild-type intratumor Tregs upon anti-CTLA-4 antibody treatment (Figure S7F). This suggests that the upregulation of TGF-β signaling is responsible for reshaping intratumor Tregs into the weaker phenotype.

**Figure. 3.**
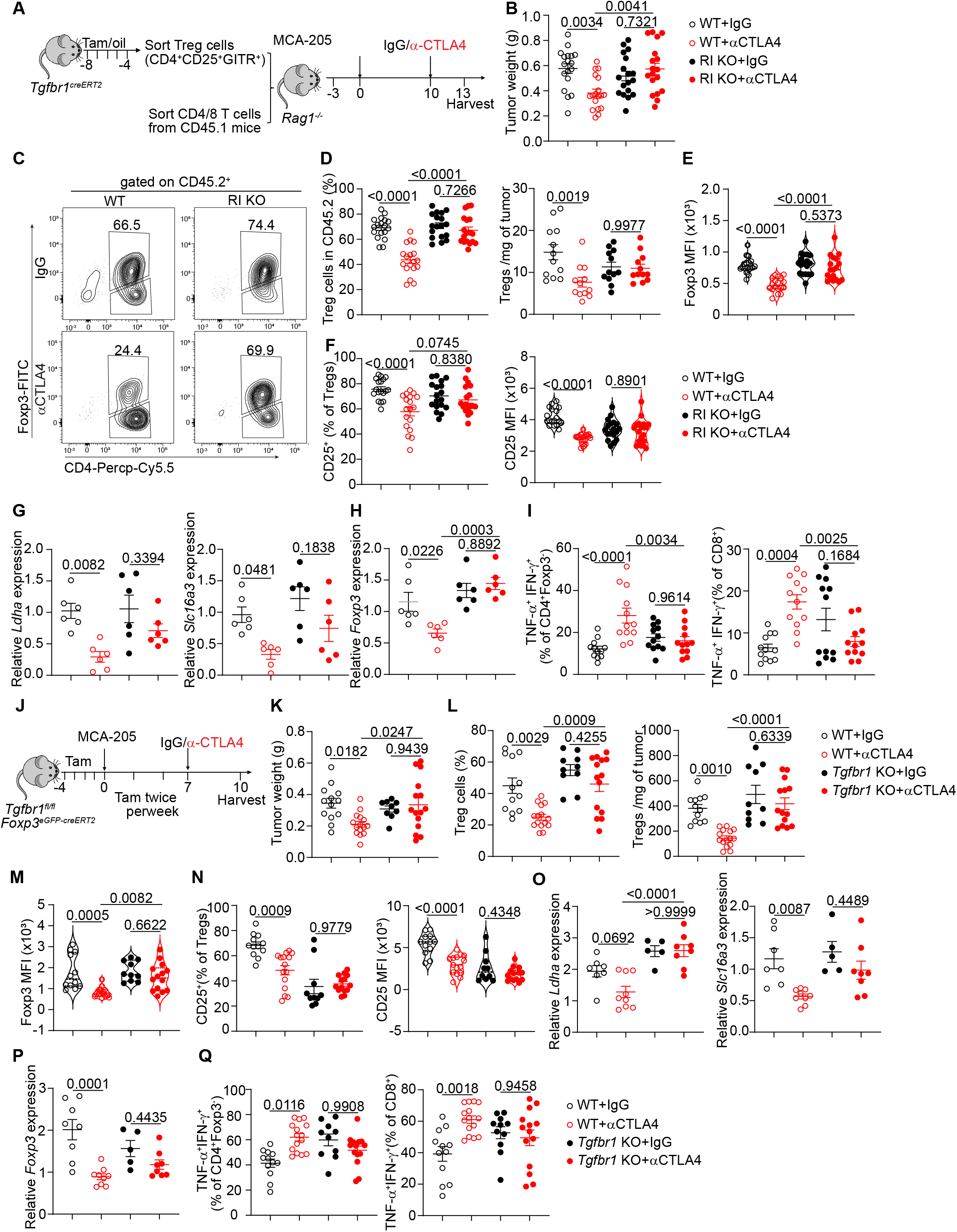
Deletion of *Tgfbr1* in Tregs abolishes αCTLA-4 mediated anti-tumor immunity. (A-B) Left, experimental protocol (A), and tumor weight (B) from *Rag1^-/-^* mice that were adoptively transferred with either WT or *Tgfbr1* ko Tregs (CD45.2^+^) plus naïve CD4^+^ and CD8^+^ T cells from CD45.1^+^ strain. (C) Flow cytogram plots of CD45.2 ^+^ T cells from tumor-bearing *Rag1^-/-^* mice that received WT or *Tgfbr1* ko Tregs, followed by the treatment with IgG or αCTLA-4 antibodies. (D) Left, Percentage of Tregs (CD4^+^Foxp3^+^) among CD45.2^+^ T cells from tumors, and right, Tregs number per mg of tumor in *Rag1^-/-^*mice bearing sarcoma tumor MCA-205 cells. (E) Quantification of Foxp3 MFI in intratumor Tregs from indicated groups. (F) Left, percentage of CD25^+^ cells among Tregs from tumors, and right, quantification of CD25 MFI in intratumor Tregs from indicated groups. (G-H) Expression of *Ldha*, *Slc16a3* (G) and *Foxp3* (H) mRNA in intratumor Tregs from indicated groups. (I) Quantification of TNF-α^+^ IFN-γ^+^ producing CD4^+^Foxp3^-^ cells or CD8^+^ T cells in tumors from indicated groups treated with IgG or αCTLA4 antibody. (J) *Tgfbr1^f/f^Foxp3^cre-ERT^*^2^ mice were given oil or tamoxifen for five consecutive 5 days before tumor injection, and then twice per week, followed by IgG or αCTLA-4 treatment for 3 days. (K) Tumor weight in mice treated as indicated. (L) Left, percentage of Tregs (CD4^+^Foxp3^+^) among CD4^+^ T cells from tumors and right, Tregs number per mg of tumor from indicated groups. (M-N) Graph depicting Foxp3 (M) and CD25 expression (N) in intratumor Tregs from indicated groups. (O-P) Expression of *Ldha*, *Slc16a3* (O) and *Foxp3* (P) mRNA in intratumor Tregs from indicated groups. (Q) Quantification of TNF-α^+^ IFN-γ^+^ producing CD4^+^Foxp3^-^ or CD8+ T cells in tumors from WT and *Tgfbr1* ko mice treated with IgG or αCTLA4 antibody. Data in (A-F, I-N, Q) are pooled from two independent experiments. Data in (G-H, O-P) are representative of two independent experiments with similar results. Significance was determined by one-way analysis of variance (ANOVA) with Tukey’s multiple comparison. Data are shown as mean ± s.e.m.

In the second approach, we used the tumor model in *Tgfbr1^f/f^ Foxp3^cre-ERT^*^2^ mice, where inducible deletion of *Tgfbr1* specifically in Tregs is achieved with tamoxifen treatment (Figure 3J). While inoculating *Tgfbr1f/fFoxp3^cre-ERT^*^2^ mice with MCA-205 tumor cells did not show any difference in tumor growth compared with WT control mice, deletion of *Tgfbr1* in Tregs resulted in resistance to neoadjuvant anti-CTLA-4 therapy (Figure 3K). The characterization of intratumor Tregs showed that Treg frequency, number, and Foxp3 and CD25 expression in *Tgfbr1^f/f^Foxp3^cre-ERT^*^2^ mice were unaffected, except for a reduction in CTLA-4 expression similar to that of the wild-type intratumor Tregs in response to anti-CTLA-4 administration (Figures 3L-3N, S8A and S8B). Notably, the expression of FcγRs is not reduced in tumor antigen presenting cells (APCs) in *Tgfbr1*^-/-^ mice (Figures S8C-S8F). The data suggests that the loss of anti-CTLA-4 antibody mediated tumor suppression was not primarily due to the different expression of CTLA-4 in the intratumor Tregs and the FcγRs in APCs ^57^. Analysis of *mRNA* expression in intratumor Tregs from *Tgfbr1^f/f^Foxp3^cre-ERT^*^2^ mice revealed that the reductions of *Ldha* and *Slc16a3*, and *Foxp3*, *Ccr8* and *Klrg1* expression observed in the wild-type Tregs caused by CTLA-4 treatment were completely abolished (Figures 3O, 3P and S8G). This set of data resembles intratumor Tregs from the mice adoptively transferred with *Tgfbr1*^-/-^ Tregs (Figures 3A-3H). Consequently, the upregulation of IFN-γ^+^TNF-α^+^-producing CD4^+^Foxp3^-^ and CD8^+^ T effector cells in the wild-type tumors after anti-CTLA-4 antibody treatment was completely abolished in *Tgfbr1^-/-^* tumors (Figures 3Q and S8H), explaining the ineffectiveness of the antibody therapy in the *Tgfbr1^-/-^* tumor (Figure 3K).

The similar levels of reduction of CTLA-4 in intratumor Tregs (Figures S7F and S8B), yet completely different tumor growth (Figures 3B and 3K) between the *Tgfbr1^-/-^* and wild-type mice, suggest that the CD80 and CD86 deprivation by CTLA-4 on intratumor Tregs might not serve as a major mechanism for anti-CTLA-4 mediated suppression of tumor growth. Indeed, measurement of CD80 and CD86 in APCs in the tumor tissues showed no differences between the *Tgfbr1^-/-^* and wild-type mice or even higher in the *Tgfbr1^-/-^* mice upon anti-CTLA-4 antibody therapy (Figures S8I and S8J). Thus, the data collectively provide compelling evidence that the instability and the compromised function of intratumor Tregs are mediated by the upregulated TGF-β signaling driven by CTLA-4 engagement upon anti-CTLA-4 antibody therapy.

### CTLA-4 is required for TGF-β signaling in tumor Tregs upon anti-CTLA-4 treatment

Tregs in the tumor tissues express high levels of CTLA-4 ^29^, which signals to upregulate TGF-β signaling upon anti-CTLA-4 antibody engagement *in vivo* and *in vitro* (Figures 1I and 2D). To directly determine the direct role of CTLA-4 for the upregulation of TGF-β signaling in tumor Tregs *in vivo*, which would consequently affect their stability and function to confer anti-tumor effects of CTLA-4 antibody, we implanted MCA-205 tumor cells in transgenic mice with specific *Ctla4* deletion in Tregs induced by tamoxifen treatment (*Ctla4f/f-Foxp3^eGFP-Cre-ERT^*^2^) ^57^ (Figure 4A and S9A). In this condition, deletion of *Ctla4* in Tregs completely abolished the tumor repression following anti-CTLA-4 treatment (Figure 4B), confirming an essential role of CTLA-4 in Tregs. Notably, deletion of *Ctla4* dramatically increased the frequency of Tregs in the tumor tissues, further validating a restrict function of CTLA-4 in Treg expansion (Figure 4C) ^57,58^, similar to its role in normal T cells ^18,45,46^. Remarkably, while anti-CTLA-4 antibody treatment significantly decreased the number of intratumor Tregs in wild-type mice as expected, the antibody completely failed to change the intratumor Tregs in the *Ctla4* knockout mice (Figure 4C) Importantly, the upregulation of *Tgfbr1* in the tumor Tregs induced by anti-CTLA-4 treatment was completely abrogated in the *Ctla4* ko mice, further confirming the direct effect of the antibody on tumor Tregs (Figure 4D). Consequently, the inhibitions of *Il2ra* (Figures 4E and S9B), and *Ldha and Slc16a3* (Figure 4F) in CTLA-4 antibody-treated intratumor Tregs were also completely abolished in the *Ctla4* ko mice. In addition to the increase in Treg frequency (Figure 4C), the levels of Foxp3 were also upregulated in the *Ctla4* ko tumor Tregs spontaneously (Figure 4G). However, anti-CTLA-4 antibody treatment failed to alter the levels of Foxp3 expression in the ko Tregs, whereas the littermate wild-type tumor Tregs showed significant reduction of Foxp3 upon anti-CTLA-4 treatment as expected (Figure 4G). Similarly, the downregulation of *Ccr8* and *Klrg1* gene expression in wild-type tumor Tregs induced by anti-CTLA-4 antibody treatment was also completely abolished in the *Ctla4* ko tumor Tregs (Figures 4H and S9C-S9D).

**Figure. 4.**
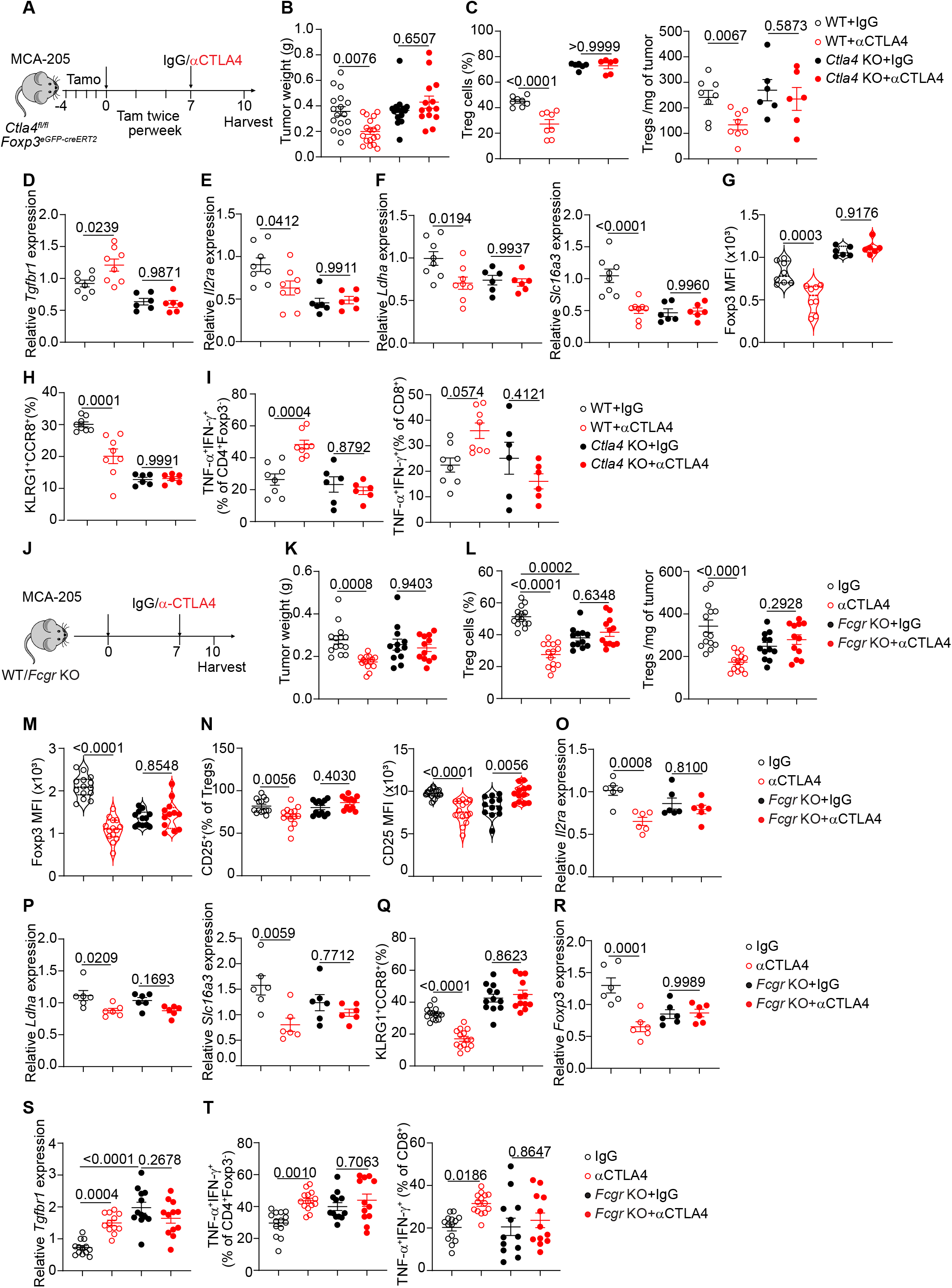
CTLA-4 presence in Tregs and FcγR expression in tumor microenvironment are required for αCTLA-4 mediated upregulation of TGF-β signaling in intratumor Tregs. (A) Schematic for *Ctla4* deletion in Tregs. (B) Tumor weight in mice treated as indicated. (C) Left, Percentage of Tregs (CD4^+^Foxp3^+^) among total CD4^+^ T cells from tumors, and right, Tregs number per mg of tumor in indicated groups. (D-F) Expression of *Tgfbr1*, *Il2ra*, lactate metabolism genes (*Ldha* and *Slc16a3*) in intratumor Tregs between IgG and αCTLA-4 treatments from WT and *Ctla4* ko mice. (G) Analysis of Foxp3 MFI in intratumor Tregs from indicated groups. (H) Analysis of KLRG1 and CCR8 expression from intratumor Tregs in indicated groups. (I) Percentage of TNF-α^+^ IFN-γ^+^ producing CD4^+^Foxp3^-^ cells or CD8^+^ T cells from indicated groups. (J) Schematic for experiment procedure in WT and *Fcgr* ko mice treated with IgG or αCTLA4 antibody. (K) Tumor weight in mice treated as indicated. (L) Percentage of Tregs (CD4^+^Foxp3^+^) among total CD4^+^ T cells from tumors and Tregs number per mg of tumor in indicated groups. M, Foxp3 MFI in intratumor Tregs in indicated groups. (N-O) Analysis of CD25 expression in intratumor Tregs at both protein and mRNA levels in indicated groups. (P) RT-PCR analysis of the expression of *Ldha* and *Slc16a3* in intratumor Tregs. (Q) Analysis of the percentage of KLRG1^+^CCR8^+^ in intratumor Tregs treated with IgG or αCTLA4 from WT and *Fcgr* ko mice. (R-S) RT-PCR analysis of *Foxp3*(R) and *Tgfbr1* (S) in intratumor Tregs from indicated groups. (T) Quantification of TNF-α^+^ IFN-γ^+^ producing CD4^+^Foxp3^-^ cells or CD8^+^ T cells in tumors from WT and *Fcgr* ko mice treated with IgG or αCTLA4 antibody. Data in (A-I, O, P, R) are representative of two independent experiments with similar results. Data in (J-N, Q, S-T) are pooled from two or three independent experiments. Significance was determined by one-way analysis of variance (ANOVA) with Tukey’s multiple comparison. Data shown as mean ± s.e.m.

To further confirm this, we cultured Tregs isolated from *Ctla4* ko and wild-type mice with TCR stimulation in the presence and absence of cross-linking anti-CTLA-4 antibody. We found that *Ctla4* ko Tregs also revoked all the instability signatures, characterized by the completely loss of the upregulation of *Tgfbr1* and the suppressed CD25 as well as lactate metabolism markers induced by CTLA-4 engagement in cultures (Figures S9E-S9G), resemble to the changes in tumor Tregs in mice treated with anti-CTLA-4 antibody (Figures 4D-4F). Consequently, the IFN-γ^+^TNF-α^+^ cells in both CD4^+^Foxp3^−^ and CD8^+^ effector T cells failed to increase in the *Ctla4* ko tumors after anti-CTLA-4 antibody treatment (Figures 4I and S9H-S9I). The data indicate that the presence of CTLA-4 in the intratumor Tregs is not only critical to restrict their expansion and Foxp3 expression, but also essential for mediating the instability and functional deficiency of tumor Tregs and the consequent antitumor effects by anti-CTLA-4 antibody treatment through TGF-β signaling.

### FcγR is required for anti-CTLA-4-mediated upregulation of TGF-β signaling in intratumor Tregs

Antibody IgG Fc receptor gamma chain (FcγR) has been suggested to be crucial for mediating the anti-tumor activity induced by anti-CTLA-4 antibody treatment through reduction of tumor Tregs ^5,29^. As we have demonstrated that CTLA-4 antibody treatment induced instability and functional deficiency of intratumor Tregs through upregulation of TGF-β signaling, we next investigated the function of FcγR in anti-CTLA-4 antibody mediated effects by utilizing γ-chain-knockout mice, which are deficient in the γ chain subunit of the FcγRⅠ, Ⅲ, Ⅴ subtypes (*Fcgr* ko) (Figure 4J). Consistent with the previous finding ^5^, *Fcgr* ko mice did not exhibit a positive response to anti-CTLA-4 injection-induced tumor repression (Figure 4K). Although the *Fcgr* ko tumors showed spontaneously lower frequency of Tregs compared to wild-type tumor, the anti-CTLA-4 antibody therapy completely failed to decrease intratumor Tregs in the *Fcgr* ko mice (Figure 4L). We also observed that Foxp3 expression (MFI) was retained in the *Fcgr* ko mice after anti-CTLA-4 injection, as was the expression of CD25 (Figures 4M, 4N and S10A). Analysis of the gene profile in the intratumor Tregs revealed that indeed the decreases in *Il2ra*, *Ldha and Slc16a3*, as well as functional markers *Foxp3*, *Ccr8,* and *Klrg1* induced by anti-CTLA-4 treatment in the wild-type mice were completely abolished in the *Fcgr* ko Tregs in the tumor tissues (Figures 4O-4R and S10B). Importantly, deletion of *Fcgr* in mice abrogated the upregulation of *Tgfbr1* in intratumor Tregs induced by anti-CTLA-4 treatment (Figure 4S). Consequently, there was no upregulation of TNFα^+^IFNγ^+^ producing CD4^+^ Foxp3^−^ and CD8^+^ effector T cells in the tumor tissues of *Fcgr* ko mice after anti-CTLA-4 antibody treatment (Figures 4T, S10C and S10D), explaining the inability of anti-CTLA-4 antibody to suppress tumor growth in the ko mice (Figure 4J). Intriguingly, *Fcgr* ko mice exhibited a spontaneously lower frequency of intratumor Tregs compared to the wild-type mice (Figure 4L), which should be at least in part due to the decrease in their proliferation determined by Ki67^+^ staining (Figure S10E).

We next investigated which subtype(s) of FcγRs is involved in anti-CTLA-4-mediated upregulation of *Tgfbr1*. As it has been demonstrated that FcγR IV plays a key role in the reduction of intratumor Tregs following therapeutic anti-CTLA-4 antibody administration^5,29,59^, we studied whether FcγR IV was required for the upregulation of TGFβ signaling in Tregs after anti-CTLA-4 antibody treatment by administering anti-CD16.2 antibodies to block FcγRIV expression in the tumor-bearing mice. As expected, the expression of FcγR IV, but not FcγRI or FcγRⅢ, is significantly reduced in macrophages/monocytes after anti-CD16.2 treatment (Figure 5H). We show that blocking FcγRIV significantly reversed the anti-tumor therapeutic effects of anti-CTLA-4 antibody as reported before ^60^ (Figure 5A). The reductions of Tregs, and the decreases in CD25 and Foxp3 expression in intratumor Tregs driven by anti-CTLA-4 antibody therapy were abrogated when FcγR IV was blocked by its specific antibody CD16.2 (Figures 5B-5D). Consequently, the increased TNFα^+^IFNγ^+^ CD4^+^ Foxp3^−^ and CD8^+^ effector T cells in the tumor tissues were also abolished with anti-CD16.2 administration (Figures 5E and 5F). Importantly, the anti-CTLA-4 antibody-mediated upregulation of *Tgfbr1* in intratumor Tregs was completely abolished by the blockade of FcγR IV (Figure 5G). Strikingly, the up-regulation of *Tgfbr1* after 4 hours treatment of anti-CTLA-4 treatment was also abrogated with anti-FcγR IV blockade antibody (Figure 5I). The data collectively provide compelling evidence that the upregulation of TGF-β signaling in intratumor Tregs induced by CTLA-4 engagement is dependent on FcγR expression, most likely through FcγR IV.

**Figure. 5.**
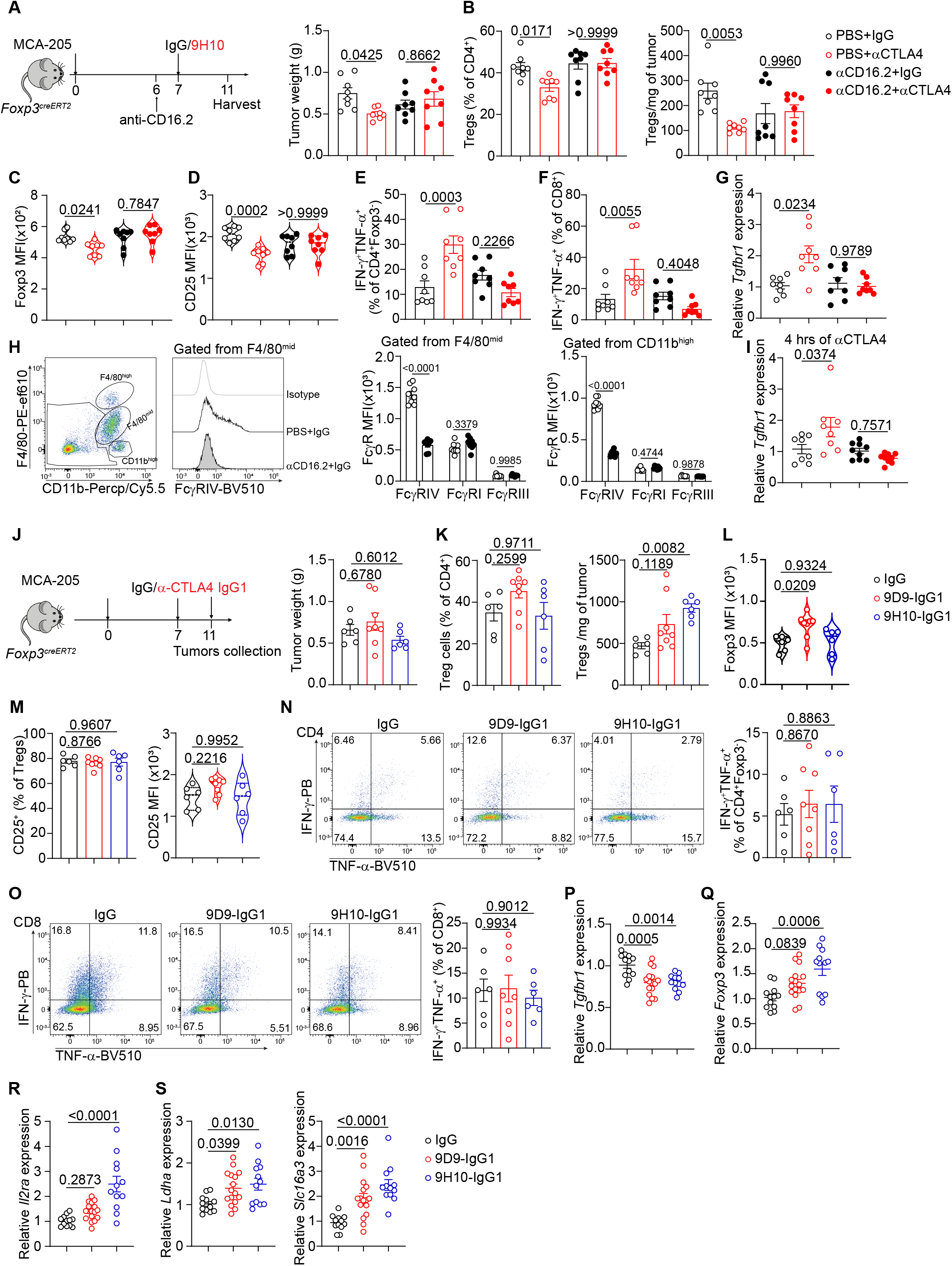
Anti-CTLA-4 induced upregulation of TGFβ signaling in tumor Tregs is dependent on the presence of FcγR IV. (A) Left, Diagram showing the experimental procedure, and right, tumor weight as indicated. (B) Left, Percentage of Tregs (CD4^+^Foxp3^+^) among total CD4^+^ T cells from tumors and right, Tregs cell number per mg of tumor in mice bearing sarcoma MCA-205 cells with αCTLA-4 therapy. (C) Quantification of Foxp3 MFI in intratumor Tregs from IgG vs. αCTLA-4 treatments. (D) Quantification of CD25 MFI in intratumor Tregs from IgG vs. αCTLA-4 treatment. (E-F) Percentage of TNF-α^+^IFN-γ^+^ producing CD4^+^Foxp3^−^ (E) or CD8^+^ T cells (F) in tumors. (G) Expression of *Tgfbr1* mRNA in intratumor Tregs by RT-PCR from indicated groups. (H) Left, Gating strategy used for identification of tumor-infiltrating innate immune cells, and right, analysis of FcγRs MFI expression in F4/80^high^ cells, F4/80^mid^ cells and CD11b^high^ cells. (I) Expression of *Tgfbr1* mRNA in intratumor Tregs by RT-PCR from indicated groups with αCTLA-4 treatments for 4 hours. (J) Left, Diagram showing the experimental procedure, and right, tumor weight as indicated. (K) Left, Percentage of Tregs (CD4^+^Foxp3^+^) among total CD4^+^ T cells from tumors and right, Tregs cell number per mg of tumor in mice bearing sarcoma MCA-205 cells with αCTLA-4 therapy. (L) Quantification of Foxp3 MFI in intratumor Tregs from IgG vs. αCTLA-4 treatments. (M) Left, percentage of CD25^+^ cells among Tregs from tumors, and right, quantification of CD25 MFI in intratumor Tregs from IgG vs. αCTLA-4 treatment. (N-O) Percentage of TNF-α^+^IFN-γ^+^ producing CD4^+^Foxp3^−^ (N) or CD8^+^ (O) T cells in tumors from indicated groups. (G-J) Expression of *Tgfbr1* (G), *Foxp3* (H), *Il2ra* (I) and *Ldha, Slc16a3* (J) mRNA in intratumor Tregs by RT-PCR from indicated groups. Data is representative of two independent experiments with similar results (J-S) or are pooled from two independent experiments with similar results (A-I). Significance was determined by one-way analysis of variance (ANOVA) with Tukey’s multiple comparison (A-G and I-S) or two-way analysis of variance (ANOVA) with Sidak’s multiple comparison (H). Data shown as mean ± s.e.m.

The finding that blockade of FcγR IV abrogates the upregulation of TGFβ signaling and anti-tumor effects induced by anti-CTLA-4 antibody treatment suggests that other FcγRs such as FcγRⅡ/Ⅲ might be dispensable for anti-CTLA-4 -mediated upregulation of *Tgfbr1*. We next treated tumor mice with anti-CTLA-4 in mouse IgG1 isoform (anti-CTLA-4-mIgG1), which has high affinity to bind with FcγRⅡ/Ⅲ. We treated the tumor mice with two mIgG1 anti-CTLA-4 antibodies (9D9-IgG1, and 9H10-IgG1) respectively (Figure 5J), and observed that both anti-CTLA-4-mIgG1 antibodies failed to suppress tumor growth (Figure 5J), which is accompanied with no reduction of Treg number and no decrease in CD25 proteins in Tregs (Figures 5K-5M). Consequently, both anti-CTLA-4-mIgG1 antibodies failed to enhance TNFα^+^IFNγ^+^ producing CD4^+^ Foxp3^−^ and CD8^+^ effector T cells compared to isotype IgG treated mice (Figures 5N and 5O). Importantly, both anti-CTLA-4-mIgG1 antibodies were unable to upregulate *Tgfbr1*, instead, they might decrease *Tgfbr1* expression in intratumor Tregs compared to isotype IgG treated mice (Figure 5P). Consequently, these two anti-CTLA-4 antibodies failed to decrease *Il2ra*, *Foxp3*, and genes related to lactate metabolism, instead increased the expression of the afteromentioned genes (Figures 5Q-5S). The data suggest that these CTLA-4-mIgG1 antibodies, despite their high-binding to FcγR II/III, may act like a blocking antibodies to promote Tregs stability and survival through reduction of *Tgfbr1*.

### Anti-CTLA-4 decreases TGF-β signaling in intratumor T effector cells by blocking CTLA-4

In addition to the constitutive expression of CTLA-4 in Tregs, activated CD4^+^ and CD8^+^ T cells in the tumor tissues can also express CTLA-4 ^18,61,62^, although the levels of expression in these effector T cells are much lower than in the Tregs ^19,58^ (Figures S11A and S11B). Having determined that anti-CTLA-4 antibody destabilizes the survival and function of intratumor Tregs through engaging the molecule, we next studied the effects of the antibody treatment on CD4^+^ Foxp3^−^ and CD8^+^ effector T cells in the same tumor tissues. For this, we first analyzed their phenotypical and functional changes upon anti-CTLA-4 treatment in MCA-205 and MC-38 models and found that the anti-CTLA-4 treatment increased the number of both CD4^+^ Foxp3^−^ and CD8^+^ T cells (Figures S11C-S11F), and these changes were likely due to their enhanced proliferation as determined by Ki67 staining (Figure S11G). Functionally, the anti-CTLA-4 antibody significantly enhanced the production of TNF-α^+^IFN-γ^+^ producing CD4^+^ Foxp3^-^ and CD8^+^ T effector cells (Figures 1C and S1I). In contrast to Tregs, however, the intratumor CD4^+^ Foxp3^−^ and CD8^+^ T cells significantly reduced their *Tgfbr1* expression, but upregulated *Il2* expression and *Il2ra* mRNA expression upon the same anti-CTLA-4 treatment (Figures S11H and S11I), suggesting a blockade of CTLA-4 in these effector T cells in the same tumor tissues.

As a major therapeutic effect of anti-CTLA-4 antibody treatment on tumors is through TGF-β signaling-mediated instability and decreased function of intratumor Tregs, the observed increase in T cell proliferation and/or higher cytokine production in CD4^+^ Foxp3^−^ and CD8^+^ T effector cells in the same tumor tissues after CTLA-4 treatment can be attributed to at least two mechanisms, namely an indirect effect by the release of the Treg suppression and/or a direct action of the antibody in the effector T cells. To distinguish these two possibilities, we first employed *Foxp3*^DTR^ transgenic mice, in which diphtheria toxin (DT) administration causes rapid and specific elimination of Tregs in the tumor tissues (Figures S1J-S1L and S12A). Analysis of tumors showed that depletion of Tregs alone dramatically increased the infiltrated CD4^+^Foxp3^−^ T cells, and the proliferation of both CD4^+^Foxp3^−^ and CD8^+^ T cells, but anti-CTLA-4 therapy further enhanced the proliferation of these T effector cells (Figures S12B and S12C), suggesting the antibody could directly target T effector cells. However, despite their increased proliferation, anti-CTLA-4 treatment failed to upregulate the frequency of IFN-γ^+^TNF-α^+^ cells within CD4^+^Foxp3^−^ and CD8^+^ T cells in the absence of Tregs in the tumors (Figure S12D), suggesting that the antibody therapeutic effects are primarily targeting Tregs (Figures S1J-S1L). We then isolated these intratumor T effector cells and demonstrated that indeed anti-CTLA-4 antibody treatment also significantly downregulated *Tgfbr1* and upregulated *Il2ra* and *Il2* expression in the absence of Tregs (Figures S12E and S12F). The data collectively suggests that anti-CTLA-4 antibody may directly increase T effector phenotype and function through blockade of CTLA-4.

However, this direct effect is subordinate to the indirect influence of Treg reduction induced by the therapeutic antibody and is not sufficient to suppress tumor growth.

The effects of anti-CTLA-4 on CD4^+^ Foxp3^−^ and CD8^+^ T effector cells in the tumor tissues were further determined in the inducible Treg-specific *Ctla4-*deficient mice. When the mice with Tregs lacking the *Ctla4* gene were treated with anti-CTLA-4 antibody, the treatment also significantly downregulated *Tgfbr1* in these intratumor CD4^+^ Foxp3^−^ and CD8^+^ T effector cells in the Treg-specific *Ctla4* ko mice (Figure S12G), further suggesting the blockade of their CTLA-4 signaling. However, the heightened intratumor Tregs due to the lack of CTLA-4 superseded any CTLA-4 blockade effects on the non-Treg T cells, and consequently failed to show any antitumor activity by the antibody (Figures 4B and 4C).

Further supporting evidence was drawn from the *Fcgr* ko mice. In the *Fcgr* ko mice, CTLA-4 antibody treatment blocked CTLA-4 in all T cells in the tumor tissues, because of the lack of FcγR cross-linking. Indeed, both CD4^+^ Foxp3^−^ and CD8^+^ T effector cells in the tumor tissues showed downregulated *Tgfbr1* expression with anti-CTLA-4 treatment compared to control IgG-treated mice (Figure S12H). The blockade of CTLA-4 in intratumor Tregs led to complete failure of Treg reduction and no decrease in their function (Figures 4L and 4M). Under this pressure, blockade of CTLA-4 in CD4^+^Foxp3^−^ and CD8^+^ T cells in the tumor tissues did not increase their cytokine IFN-γ^+^ TNF-α^+^ production (Figures 4T, S10C and S10D), further confirming the limitation of blockade of CTLA-4 in the effector cells by the antibody in the anti-tumor effects.

Importantly, CTLA-4 antibody treatment of mice deficient in *Tgfbr1* specifically in Tregs also resulted in a downregulation of *Tgfbr1* expression in CD4^+^ Foxp3^−^ and CD8^+^ T effector cells in the tumor tissues, further confirming a direct blockade of CTLA-4 on these intratumor effector T cells (Figure S12I). However, this led to no increase in TNF-α^+^IFN-γ^+^ cells in either CD8^+^ or CD4^+^Foxp3^−^ T cells, because of the lost function of the anti-CTLA-4 antibody in *Tgfbr1* ko tumor Tregs (Figures 3Q and S8H). Thus, this blockade of CTLA-4 and activation in T effector cells was not sufficient to mount effective anti-tumor therapeutic effects (Figures 3B and 3K).

We next validated in a series of *in vitro* experiments that the downregulation of *Tgfbr1* expression and signaling in the T effector cells was indeed due to the blockade, but not signaling, of CTLA-4 on these cells. The blockade of CTLA-4 with CTLA-4 Fab’ antibody resulted in the downregulation of *Tgfbr1* expression with an upregulation of *Il2* and CD25 expression in naïve CD4^+^ T cells and CD8^+^ T cells stimulated with low doses of anti-CD3 and anti-CD28 in culture (Figures S13A and S13B). Similar results were obtained in naïve CD4^+^ T cells stimulated with a low dose of anti-CD3 antibody in the presence of irradiated splenic antigen presenting cells (APCs) (Figure S13C). In contrast, engagement of CTLA-4 in naive CD4^+^ and CD8^+^ T cells prevented the downregulation of *Tgfbr1* expression induced by TCR activation ^13^, which was accompanied with downregulation of IL-2 production and CD25 expression, as well as decreased IFN-γ production (Figures S13D-S13G). Importantly, stimulation of CTLA-4 in *Tgfbr1* ko CD4^+^ and CD8^+^ T cells with crosslinking anti-CTLA-4 antibody together with TCR stimulation failed to significantly suppress IFN-γ production (Figures S13H and S13I).

Taken together, our results lead us to conclude that anti-CTLA-4 antibody therapy stimulates CTLA-4 in Tregs but blocks it in CD4^+^ Foxp3^−^ and CD8^+^ T effector cells in the same tumor tissues. However, the blockade of CTLA-4 in CD4^+^ and CD8^+^ effector T cells alone is insufficient to mediate the antibody-driven tumor suppression.

### CTLA-4 antibody treatment increases peripheral Tregs by reducing TGF-β signaling

In contrast to the marked reduction of intratumor Tregs, the frequency of Tregs in the spleen and LNs were significantly increased in the same tumor models upon therapeutic CTLA-4 antibody 9H10 treatment (Figure 6A). The same results were obtained by therapy with a different anti-CTLA-4, 9D9 (Figure 6B). The increase in the peripheral Tregs is attributed to the enhanced Treg proliferation, as determined by Ki67 staining (Figure 6C), whereas the Tregs in the tumor tissues showed no changes in proliferation (Figure S3A). Consistently, the expression of CD25 protein and the frequency of CD25^+^ Tregs in the spleens and LNs were also markedly increased (Figure 6D). In contrast to tumor tissues, the expression of *Ldha* and *Slc16a3* were increased in peripheral Tregs in mice treated with anti-CTLA-4 antibody (Figure 6E). Importantly, Foxp3 expression was also increased in the peripheral Tregs upon CTLA-4 antibody therapy (Figure 6F), suggesting an increased suppressive function of these Tregs. Indeed, purified Tregs from spleen treated with anti-CTLA-4 antibody exhibited much stronger suppressive activity than Tregs from spleen treated with control antibody from the same tumor mice in a standard Treg suppression assay *in vitro* (Figure 2T). whereas the tumor Tregs showed the opposite (Figure 2T). The opposite phenotypical and functional changes upon anti-CTLA-4 treatment between intratumor and peripheral Tregs suggest that the antibody blocks the CTLA-4 in peripheral Tregs, and consequently increases their stability, proliferation, and function by downregulating the TGF-β signaling. Indeed, measurement of *Tgfbr1* gene expression in the peripheral Tregs showed a significant decrease upon anti-CTLA-4 treatment compared to control antibody-treated Tregs (Figure 6G). It was confirmed *in vitro* that Tregs showed a significant decrease in *Tgfbr1* expression, when their CTLA-4 was blocked with the specific CTLA-4 Fab’ blocking antibodies (Figure 2B). Consistently, blockade of CTLA-4 in peripheral Tregs enhanced *Ldha* and *Slc16a3* expression in culture (Figure 6H).

**Figure. 6.**
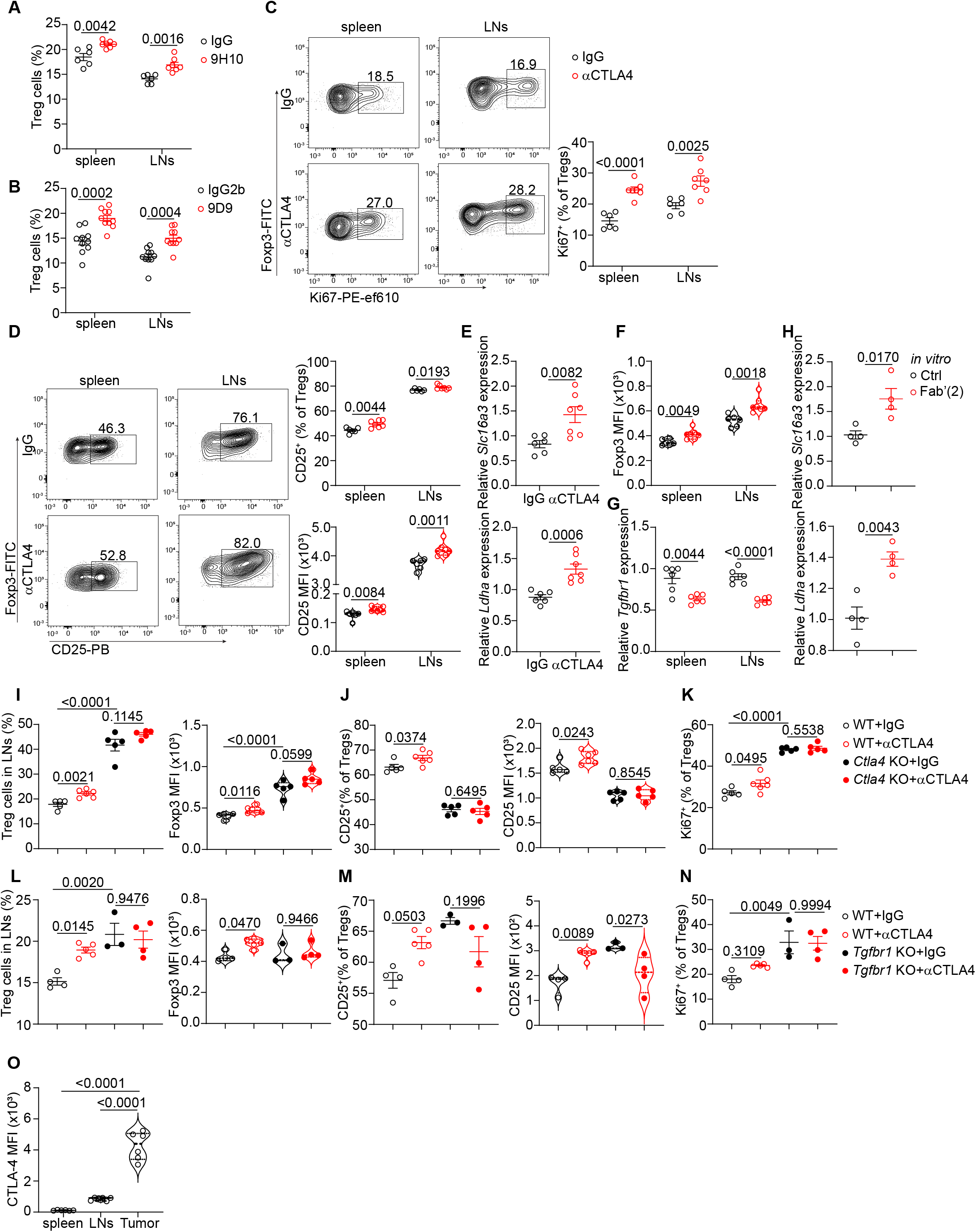
αCTLA-4 immunotherapy increases peripheral Tregs in tumor-bearing mice. (A-B) Percentage of Tregs in spleen and draining lymph nodes (LNs) from *Foxp3^EGFP^* mice bearing MCA-205 tumors treated with 9H10 (A) or 9D9 (B) antibody. (C) Left, Representative cytometry flow plot of Ki67 staining in Treg cells from spleen and LNs, and right, percentage of Ki67 in Tregs from spleen and LNs. (D) Left, Representative flow cytometry of CD25 staining in Tregs from spleen and LNs, and right, percentage of CD25 expression in Tregs from spleen and LNs. (E) Expression of *Slc16a3* and *Ldha* in Tregs from LNs between IgG vs. αCTLA-4 therapy. (F) Quantification of Foxp3 MFI in spleen and LNs from *Foxp3^EGFP^* mice bearing MCA-205 tumors treated with IgG vs. αCTLA-4 antibody. (G) RT-PCR analysis of *Tgfbr1* in Tregs from spleen and LNs. (H) Expression of *Slc16a3* and *Ldha* in isolated Tregs *ex vivo* cultured under CTLA-4 blockade condition overnight. (I) Left, percentage of Tregs, and right, quantification of Foxp3 MFI in LNs from *Ctla4^f/f^Foxp3^cre-ERT^*^2^ mice bearing MCA-205 tumors. (J) Percentage, and quantification of CD25 expression in Tregs in LNs from indicated groups. (K) Percentage of proliferation of Tregs in LNs from indicated groups. l, Left, Percentage of Tregs and right, quantification of Foxp3 MFI in LNs from *Tgfbr1^f/f^Foxp3^cre-ERT^*^2^ mice bearing MCA-205 tumors. (M) Percentage, and quantification of CD25 expression in Tregs in LNs from indicated groups. (N) Percentage of Ki67 of Tregs in LNs from indicated groups. (O) Quantification of CTLA-4 expression in Tregs in tumors and periphery lymph tissues. Data are representative of from two independent experiments with similar results. Significance was determined by unpaired two-tailed *t*-test (A-H) or one-way analysis of variance (ANOVA) with Tukey’s multiple comparison (I-O). Data shown as mean ± s.e.m.

Notably, peripheral Tregs from *Ctla4* ko mice lost the changes in all of the aforementioned markers in Tregs observed in the wild type mice upon anti-CTLA-4 antibody treatment. (Figures 6I-6K), further indicating that the blockade of CTLA-4 is responsible for the increased activation, proliferation, and function of the peripheral Tregs. Most importantly, the peripheral Tregs with *Tgfbr1* specific deficiency in tumor mice failed to exhibit the changes observed in the wild-type Tregs induced by anti-CTLA-4 antibody therapy (Figures 6L-6N), indicating a key role of TGF-β signaling in restraining peripheral Tregs ^63^. Although the exact mechanisms underlying the opposite changes between intratumor and peripheral Tregs remain to be elucidated, the high and low amounts of CTLA-4 between the intratumor and peripheral Tregs, respectively, are likely to account for the different phenotype and function upon anti-CTLA-4 antibody treatment (Figure 6O).

### Engagement of CTLA-4 in human Tregs by ipilimumab suppresses Tregs by upregulating *TGFBR1 in vitro*

Having established that CTLA-4 engagement suppresses tumor Tregs by upregulation of TGF-β signaling, we next leveraged the CTLA-4 signaling effects on human Tregs. For this, we performed both engagement and blockade experiments with the anti-human CTLA-4 antibody ipilimumab in purified human Tregs in culture. Human Tregs (CD4^+^CD25^hi^ CD127^−^) ^64^ were sorted and cultured with cross-linked ipilimumab with goat anti-human IgG to mimic engagement in the presence of anti-human αCD3/CD28 beads. Engagement of CTLA-4 in human Tregs dramatically upregulated TGF-β signaling (*TGFBR1* and *TGFBR2* expression), impaired lactate metabolism (*LDHA* and *SLC16A3* expression) and IL-2RA expression, and increased glucose metabolism (*HK2* expression) (Figures 7A-7C). In contrast, blockade of CTLA-4 with soluble ipilimumab in the presence of a low dose of αCD3 (0.01µg/ml) together with soluble anti-human CD28 antibodies led to a completely opposite result in human Tregs. (Figures 7D-7F). The data indicate that, as in murine Tregs, CTLA-4 signaling also increases *TGFBR1* expression in human Tregs.

**Figure. 7.**
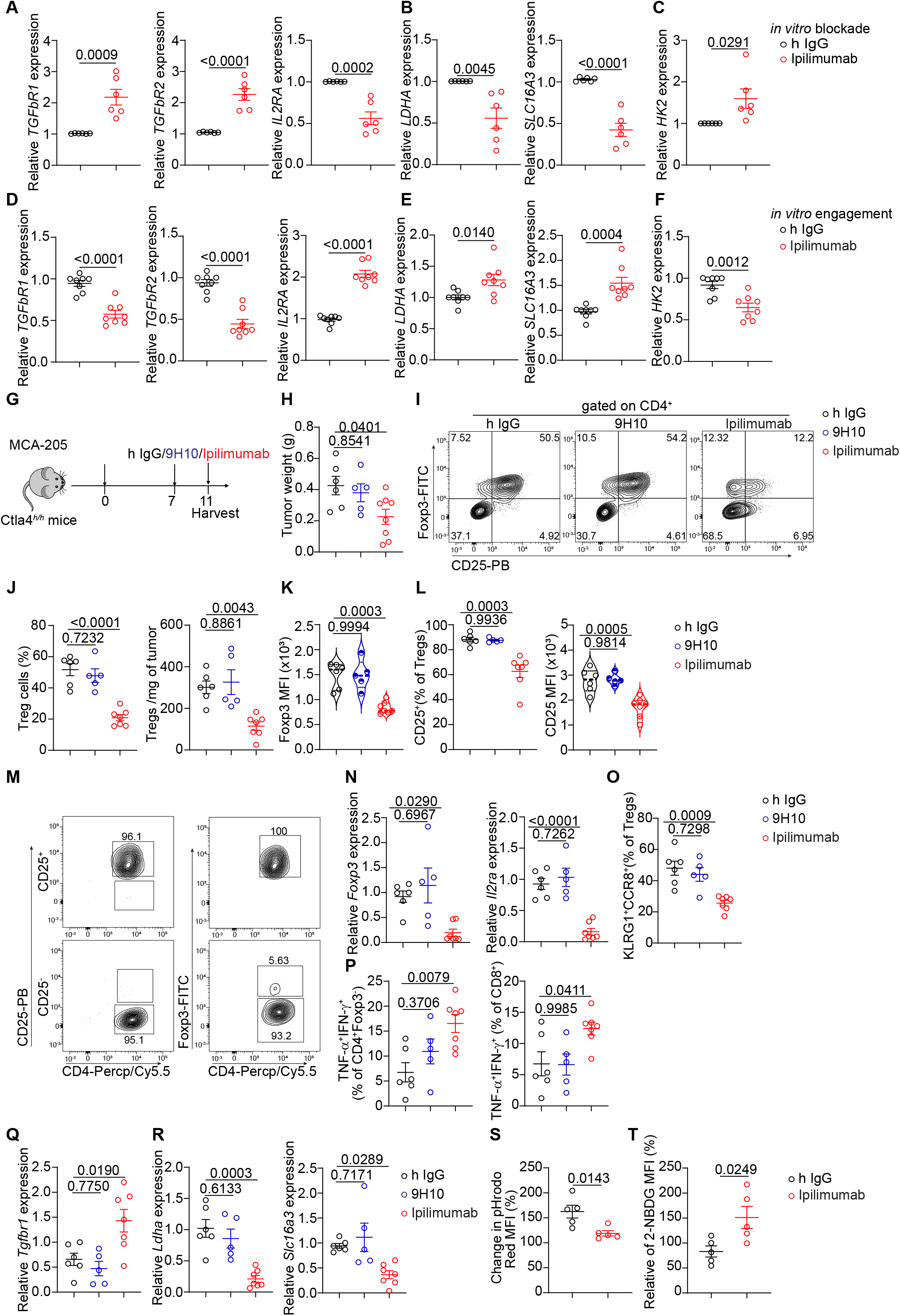
Ipilimumab treatment upregulates TGF-β signaling in intratumor Tregs to suppress tumor in human *CTLA-4* knock-in mice. (A-C) Expression of *TGFBR1*, *TGFBR2*, *IL2RA* (A), *LDHA*, *SLC16A3* (B) and *HK2* (C) in human Tregs cultured with human coactivator αCD3/CD28 beads in the presence of human IgG or ipilimumab plus mouse-anti human IgG overnight. (D-F) Expression of *TGFBR1*, *TGFBR2*, *IL2RA* (D), *LDHA*, *SLC16A3* (E) and *HK2* (F) in human Tregs cultured with low dose of αCD3 (0.01 ug/ml) together with CD28 (1 ug/ml) in presence of soluble human IgG or ipilimumab overnight. (G) Schematic to access the impact of ipilimumab or 9H10 on human *CTLA-4* knock-in mice. (H) Tumor weight from tumor-bearing human *CTLA-4* knock-in mice that received with human IgG, 9H10 or ipilimumab treatment. (I) Flow cytometry of Foxp3 and CD25 gated from tumor-infiltrating CD4^+^ T cells from indicated groups. (J) Left, percentage of Tregs (CD4^+^Foxp3^+^) among CD4^+^ T cells from tumors, and right, Treg number per mg of tumor in human *CTLA-4* knock-in mice bearing sarcoma tumor MCA-205 cells. (K) Quantification of Foxp3 MFI in intratumor Tregs from indicated groups. (L) Left, percentage of CD25^+^ among Tregs from tumors, and right, quantification of CD25 MFI in tumor-infiltrating Tregs from indicated groups. (M) Flow cytometric analysis of Foxp3 expression from sorted CD25^high^ and CD25^−^ population from tumors. (N) Expression of *Foxp3* and *Il2ra* mRNA in tumor-infiltrating Tregs from indicated groups. (O) Percentage of KLRG1^+^CCR8^+^ among Tregs from tumors (P) Percentage of TNF-α^+^IFN-γ^+^ producing CD4^+^Foxp3^−^ cells or CD8^+^ T cells in tumors. (Q) RT-PCR analysis of the expression of *Tgfbr1* mRNA in intratumor Tregs. (R) Expression of *Ldha* and *Slc16a3* mRNA in intratumor Tregs by RT-PCR. (S) Measurement of relative lactic acid uptake by intratumor Tregs. (T) Quantification of 2-NBDG uptake in intratumor Tregs from indicated groups. Data in (G-S) are representative of two independent experiments with similar results. Data in (A-F) are pooled from three independent experiments with similar results. Significance was determined by one-way analysis of variance (ANOVA) with Tukey’s multiple comparison (G-R) or unpaired two-tailed *t*-test (A-F, S-T). Data shown as mean ± s.e.m.

### Ipilimumab therapy suppresses tumors in human CTLA-4 knock-in mice by upregulation of TGF-β signaling in tumor Tregs

To investigate whether CTLA-4 engagement-mediated dysfunctional Treg cells were replicable in the clinical patients treated with human anti-CTLA-4 (ipilimumab, IgG1), we employed human CTLA-4 knock-in mice ^65^, in which Tregs express human CTLA-4 and thus respond specifically to anti-human CTLA-4 antibody. We treated tumor mice with the same sarcoma tumor cells (MCA-205) with single i.p. injection of ipilimumab for 3-4 days (Figure 7G). In addition to the isotype control antibody for ipilimumab, we also included a group of tumor mice treated with anti-mouse CTLA-4 (9H10) as a negative control (Figure 7G). Ipilimumab treatment significantly delayed tumor growth compared with human IgG1 control antibody, whereas 9H10 antibody treatment failed to exhibit any therapeutic effects on tumor growth (Figure 7H). Consistent with the decrease in tumor weights, ipilimumab, but not 9H10, dramatically decreased the frequency and total number of Tregs in the tumor tissues compared to control antibody injections (Figures 7I and 7J). Further cytometric analysis of Tregs in the tumor tissues revealed that ipilimumab treatment significantly inhibited the expression of both their Foxp3 and CD25, which was also confirmed by qPCR on sorted CD25^high^CD4^+^ T cells, where the percentage of Tregs was more than 98-99% (Figures 7K-7N). Additionally, the dysfunctional signatures of Tregs induced by ipilimumab administration were also evidenced by reduced expression of KLRG1 and CCR8 (Figure 7O). Consequently, the production of IFN-γ^+^ TNF-α^+^ in both CD4^+^Foxp3^−^ and CD8^+^ T cells were heightened in ipilimumab-treated mice compared with 9H10 or control IgG1 groups (Figure 7P). Importantly, an upregulation of TGF-β signaling in Tregs characterized by enhanced expression of *Tgfbr1* was detected (Figure 7Q), together with compromised lactate metabolism (Figures 7R and 7S) and increased glucose avidity (Figure 7T).

Taken together, the data strongly support that the upregulation of TGF-β signaling by CTLA-4 engagement in human tumor Tregs also correlates positively with a favorable response to anti-CTLA-4 treatment in human patients, in addition to preclinical tumor models.

## Discussion

As the first FDA approved immune checkpoint blockade antibody for immunotherapy to human cancers ^3,4^ , anti-CTLA-4 antibody has been shown to be beneficial to effectively treat multiple human cancers including melanoma, lung cancers, and kidney cancers ^2,23^. Despite the impressive success of the antibody as cancer therapy, the underlying mechanisms by which the anti-CTLA-4 antibody executes its therapeutic function in suppressing tumor growth remain largely unknown. Here, we present a previously unrecognized mechanism to explain the anti-CTLA-4 antibody-mediated anti-tumor effects, namely, that anti-CTLA-4 antibody engages CTLA-4 in intratumor Tregs to upregulate TGF-β signaling, which in turn suppresses CD25-mediated IL-2 signaling and lactate metabolism leading to reduced Treg stability and survival, and also compromises Treg suppressive function in the tumor (**Extended data Fig. 16**).

Several unprecedent conclusions can be drawn from the current studies. *First*, CTLA-4 therapeutic antibody stimulates, rather than blocks, CTLA-4 in intratumor Tregs to suppress their stability and function in tumor tissues. Supporting this conclusion are several pieces of experimental evidence. The stability and survival of Tregs, especially those in tumor tissues, require at least two important pathways, the CD25-mediated IL-2 signaling pathway^14,15^ and the lactate metabolism pathway ^35,42^. We uncovered that upon anti-CTLA-4 antibody treatment the intratumor Tregs significantly downregulate their CD25 expression, and also the expression of the *Ldha* and *Slc16a3* genes, which are key markers of lactate metabolism ^42,43,66^. Importantly, these reductions in gene expression occur very early, such as at day 3, day 1 or even 15 hours after CTLA-4 antibody administration. The reduction of both IL-2 signaling, and lactate metabolism together could cause destabilization and increase susceptibility to death, and thus could account for the reduction in the intratumor Treg numbers ^16,42^. The underlying mechanism responsible for the downregulation of CD25-mediated IL-2 signaling and lactate metabolism is ascribed to the engagement of CTLA-4 by cross-linking antibody rather than by blockade of the molecule by CTLA-4 Fab’ fragment in Tregs in culture, resembling the reduction of the genes found *in vivo*. In addition, *in vivo* deletion of FcγR in mice, which abolishes FcγR-mediated anti-CTLA-4 signaling, completely abrogates the downregulation of CD25 and lactate metabolic molecules in Tregs in the tumor tissue observed in wild-type mice after anti-CTLA-4 antibody therapy. Importantly, this FcγR-mediated effect is primarily through FcγRIV rather than other FcγRs. This conclusion of CTLA-4 engagement and signaling rather than blockade is further supported by experimental data showing that the absence of CTLA-4 specifically in Tregs completely abolishes the therapeutic effects of the anti-CTLA-4 antibody. Indeed, if the blockade of CTLA-4 were the mechanism for anti-CTLA-4 antibody therapy, mice with Treg-specific *Ctla4* knockout mice should exhibit reduced tumor growth, as Tregs lose all CTLA-4. However, these conditional ko mice fail to exhibit any interference with tumor growth. Unexpectedly, these ko mice showed a substantially increased number of Tregs in the tumor tissues spontaneously compared to the wild-type mice, further strengthening the function of CTLA-4 signaling as an inhibitor for Treg growth ^57,58^. Our data do not necessarily contradict with the findings obtained using *Foxp3^ER^ ^Cre^-Ctla4^f/f^* conditional mice ^58^, as those mice delete Foxp3 from the beginning of the mouse’s life when Foxp3 starts to be expressed, leading to uncontrolled systemic inflammation, which could suppress tumor development, whereas the inducible *Ctla4* deletion in Tregs in the transgenic mice used in our study are otherwise healthy, without any overt systemic inflammation during the period of tumor development. Finally, the fact that the same anti-CTLA-4 administration in the inducible Treg-depletion mice with same tumor models completely fails to induce anti-tumor activity further validates the key role of intratumor Tregs through CTLA-4 signaling in suppression of anti-tumor effector immunity in the tumor microenvironment. Based on all of this compelling evidence, we propose that the reduction in the number of intratumor Tregs in response to anti-CTLA-4 antibody treatment is also through the loss of stability and thus survival caused by CTLA-4 signaling in the Tregs, although an ADCC contribution to the decrease in intratumor Tregs cannot be completely ruled out ^5^.

In addition to the decrease in the number of Tregs in the tumor tissues, CTLA-4 signaling-mediated deficiency of intratumor Treg function may also contribute to the enhancement of the anti-tumor effector T cell immunity induced by the anti-CTLA-4 antibody therapy. Supporting this conclusion is the fact that Foxp3, the key molecule for Tregs function, is significantly suppressed in intratumor Tregs upon anti-CTLA-4 antibody treatment in mice. This decrease starts as early as 24 hours after the antibody injection and can last for as long as two weeks, the point at which the experiments ended. The suppressed Foxp3 expression in these treated intratumor Tregs impedes their suppressive activities toward other effector T cells. Indeed, when intratumor Tregs are isolated from the same mice treated with antibodies and the standard *in vitro* suppressive assay is performed ^55^, the intratumor Tregs treated with anti-CTLA-4 antibody showed significantly weaker suppressive function toward effector T cells compared to intratumor Tregs isolated from mice that received control antibody, consistent with the downregulation of Foxp3 in tumor Tregs in CTLA-4 treated mice. It is confirmed *in vitro* that CTLA-4 engagement by cross-linking anti-

CTLA-4 antibody decreases Foxp3 expression in Tregs. Besides Foxp3, several other molecules associated with the suppressive function of Tregs, including CCR8 and KLRG1, in the tumor tissues are also inhibited by anti-CTLA-4 therapy. The downregulation of these molecules together with Foxp3 confers the weakening of the Treg function in the tumor tissues, which together with the reduction in the number of Tregs unleashes the antitumor immune responses by CD4^+^ and CD8^+^ T effector cells to control tumor growth.

Second, CTLA-4 engagement-mediated instability and functional deficiency of intratumor Tregs occurs through upregulation of TGF-β signaling. This previously unrecognized conclusion is supported by a series of experimental evidence. First, anti-CTLA-4 antibody therapy induces significant upregulation of TGF-β signaling in intratumor Tregs. This was first shown in our global scRNA-seq analysis in in which *Tgfbr1* expression and other TGF-β signaling-associated molecules are increased in CTLA-4 antibody-treated Tregs. This was then confirmed by our semi-quantitative RT-PCR analysis, including the upregulation of *Tgfbr1* gene expression and increased phosphorylation of Smad3 (pSmad3), which are critical markers and mediators for TGF-β signaling ^13,44^. That this upregulation of TβRI-mediated TGF-β signaling is induced by CTLA-4 engagement rather than blockade is validated by *in vitro* culture experiments in which stimulation of CTLA-4 by cross-linking antibody but not blockade of it with CTLA-4 Fab’ blocking antibody enhances *Tgfbr1* gene expression. Furthering this confirmation, we found that mice lacking FcγR, which abolishes anti-CTLA-4 antibody cross-linking-induced CTLA-4 engagement, completely abrogated the increase in *Tgfbr1* upregulation in intratumor Tregs observed in wild-type mice following anti-CTLA-4 antibody treatment. Moreover, this results from a direct effect on CTLA-4 in intratumor Tregs rather than indirect influence by other factors, as intratumor Tregs with specific deletion of the *Ctla4* gene show complete failure of *Tgfbr1* gene upregulation in response to anti-CTLA-4 therapy.

Most strikingly, the causal effects of TGF-β signaling on the suppression of stability and function in intratumor Tregs are confirmed by the fact that specific deletion of TGF-β signaling in Tregs completely abolishes the suppression of the stability and function in intratumor Tregs induced by CTLA-4 engagement by anti-CTLA-4 antibody *in vitro* and *in vivo*. Consequently, the enhancement of IFN-γ^+^TNF-α^+^-producing CD4^+^ and CD8^+^ T effector cells driven by anti-CTLA-4 treatment is totally abolished, because the number and function of intratumor Tregs are increased in the mice with *Tgfbr1* deleted specifically in Tregs. This leads to the therapeutic effects of anti-CTLA-4 antibody on tumor growth being completely blocked, and actually the tumor development is even faster, and the tumors were bigger in the *Tgfbr1* ko mice than in the wild-type tumor bearing mice. Thus, TGF-β signaling in intratumor Tregs is important for anti-CTLA-4-mediated therapeutic effects on cancer, at least in preclinical animal models.

Anti-CTLA-4 therapeutic antibody administration enhances the anti-tumor effector T cells, including CD4^+^Foxp3^−^ conventional T cells, and CD8^+^ T cells ^26,36^. However, at least two possibilities account for this activation and enhancement of T effector cells, namely, the weakening of intratumor Treg-mediated suppression and/or a direct effect of the anti-CTLA-4 antibody on the T effector cells in the tumor tissues. Our studies here have indicated that anti-CTLA-4 antibody-induced Treg instability and functional deficiency play a major role in releasing the brake for Treg-mediated suppression of T effector cells.

The antibody can also directly enhance the activation of CD4^+^Foxp3^−^ conventional T cells and CD8^+^ T cells by blocking their CTLA-4 in the tumor tissues, but this blockade itself is insufficient for mounting effective anti-tumor function. There are several pieces of experimental evidence supporting this conclusion. First, in the absence of intratumor Tregs as shown in the mice with Treg-specific deletion, anti-CTLA-4 antibody can indeed increase the proliferation of CD4^+^Foxp3^−^ and CD8^+^ T cells. The directly enhanced proliferation and activation of T effector cells in response to anti-CTLA-4 antibody treatment is due to the blockade of CTLA-4, not engagement, on these effector cells by the antibody in the tumor tissues. This conclusion is based on the fact that these activated T effector cells show decreased TGF-β signaling when measured ex vivo, which can be mimicked by the CTLA-4 blockade experiment *in vitro,* in which both CD4^+^ Foxp3^−^ and CD8^+^ cells decrease *Tgfbr1*. This is further supported by the mice with Treg-specific deficiency of the *Ctla4* gene or the *Tgfbr1* gene, and the *Fcgr* deficient mice. In this set of conditional knockout mice, anti-CTLA-4 antibody also downregulates *Tgfbr1* expression in CD4^+^Foxp3^−^ and CD8^+^ T effector cells in the tumor tissues. This raises an intriguing yet exciting question, as to why the same anti-CTLA-4 could have opposite effects on Tregs and T effector cells in the same tumor tissues. Although the exact mechanism remains to be elucidated, it can be reasoned that the difference is attributable to the different levels of CTLA-4 expression between the two populations of T cells, namely, the extremally high levels in Tregs and the much lower levels in T effector cells ^27^. The limited amounts of CTLA-4 molecules on the T effector cells restrict their ability to receive the signal from the anti-CTLA-4 antibody, and thus they are unable to reach the minimal threshold to stimulate the signaling cascade through CTLA-4. Instead, the CTLA-4 signaling is blocked. This possibility needs to be explored in the future. Nevertheless, the distinct effects of CTLA-4 antibody on Tregs and T effector cells should provide excellent targets for future manipulation of these cells for more effective immunotherapy using the anti-CTLA-4 antibody.

In contrast to its effects on intratumor Tregs, anti-CTLA-4 antibody enhances peripheral Tregs by blocking CTLA-4. The number of peripheral Tregs from spleens and lymph nodes unexpectedly increased in the same tumor mice treated with anti-CTLA-4 therapeutic antibody ^8,67^, while the number of Tregs in the tumor tissues is dramatically lowered. Analysis of the peripheral Tregs revealed that the increase in Tregs can be ascribed to the enhanced T cell proliferation evidenced by the higher Ki67 staining, whereas the same intratumor Tregs show no changes in the amounts of Ki67-positive cells upon anti-CTLA-4 administration. The increased T cell proliferation may be due to heightened IL-2 signaling, as the CD25 expression is significantly enhanced in peripheral Tregs in CTLA-4 treated mice. Conversely, *Tgfbr1* expression in the same peripheral Tregs is significantly downregulated. These data together indicate the blockade of CTLA-4 on the peripheral Tregs by the anti-CTLA-4 antibody treatment. Similar to the questions raised in the T effector cells, the mechanism underlying the opposite effect by the same anti-CTLA-4 antibody between intratumor and peripheral Tregs is not known. This question is worth further investigation in future studies. However, the substantially lower levels of CTLA-4 in the peripheral Tregs compared to the intratumor Tregs may play a role. This could block rather than engage the CTLA-4 on these peripheral Tregs, consequently releasing the suppression of TGF-β signaling to enhance their activation and growth. The lack of reduction and even upregulation of the peripheral Tregs upon anti-CTLA-4 antibody treatment has significant clinical implications for designing immunotherapy by manipulating anti-CTLA-4 antibody-mediated side effects. For example, one significant side effect for anti-CTLA-4 antibody therapy is the induction of inflammation and autoimmunity in patients with cancer. In this regard, a single injection, i.e., avoiding repeated anti-CTLA-4 antibody injection, may be beneficial for the patients for both cancer suppression and prevention of immune-related adverse events (irAEs). Indeed, it has been reported that multiple rounds of repeated anti-CTLA-4 antibody injection did not show enhanced benefit for cancer suppression in patients ^22,23^, but could increase the risk of autoimmunity and inflammation.

In the context of cancer patients enrolled in clinical trials, a comprehensive understanding of the mechanism underlying immunotherapy inhibitors such as anti-CTLA4, remains elusive. Technical and ethical limitations hinder investigators’ ability to meticulously study these mechanisms in human samples immediately before and after treatment. Animal models offer unparalleled advantages for unravelling the intricate impact of different immunotherapy antibodies. Despite the lack of a definitive conclusion regarding the mechanism of action of anti-CTLA-4 in cancer patients, a multitude of preclinical studies illustrating the pivotal role of Tregs depletion have fueled the drive to develop a new class of anti-CTLA-4 antibodies with enhanced capacity to deplete Tregs in human patients, although this remains in debate ^2^. In fact, it has been recently shown that anti-CTLA-4 antibody ipilimumab therapy of melanoma patients substantially decreased Foxp3^+^ Tregs in the tumor tissues, especially in clinical responders ^68^. Resolving the discrepancy with regard to Treg changes upon anti-CTLA-4 antibody treatment between the mice and the patients requires consideration of several factors. Unlike in mice, the Foxp3^+^ Tregs in humans can be divided into several subpopulations ^33^. It is likely that CTLA-4 antibody targets and decreases certain populations of Tregs, such as CD45RA^−^ Foxp3 ^hi^ effector Tregs, but not other subsets, and thus fails to a cause dramatic difference in the overall Treg population ^69^. Consistent with this notion, it has been demonstrated that the effector CD45RA^−^Foxp3^hi^ effector Tregs are enriched in the tumor tissues, but not in the blood in the patients with cancer ^27^. Unlike in the animal models where the entire tumor tissues can be excised and different kinetics can be examined, human clinical samples are limited to a single or in the best cases to a couple of biopsies, which provide a partial and heterogeneous representation of the tissues of the whole tumor ^70^. Thus, it might be informative to analyze changes in Treg subsets in upon the CTLA-4 antibody-treated patients, rather than the entire population of Foxp3^+^ T cells. In addition, as we showed in our mouse model, anti-CTLA-4 antibody specifically decreases the stability and function in intratumor Tregs, but not the peripheral Tregs. Thus, the changes in peripheral Tregs in the patients may not reflect the actual changes in the tumor tissues. Finally, it is also possible that the anti-CTLA-4 antibody weakens the function of human intratumor Tregs without changing their number; thus, more careful analysis of the functional molecules on Tregs after CTLA-4 antibody therapy may provide some clues. Nevertheless, our results in a series of in vitro CTLA-4 engagement experiments of human Tregs in culture and in ipilimumab treated humanized tumor mice not only unveil additional underlying mechanisms explaining the reduction of tumor-infiltrating Tregs following anti-CTLA-4 treatment, potentially serving as a reference for the clinicians to predict the outcome after administration of anti-CTLA-4 treatment, but they also shed light on potential challenges in achieving of an ideal synergistic effect through combination with anti-TGF-β.

In summary, our data emphasize that the elevation of TGF-β signaling proved essential for CTLA-4 engagement in causing Tregs instability and subsequent tumors rejection. This heightened TGF-β signaling was contingent upon both CTLA-4 expression in intratumor Tregs and the presence of FcγR in the tumor tissues. We propose that the anti-CTLA-4 antibody-mediated upregulation of TGF-β signaling in intratumor Tregs represents an additional and important pathway besides ADCC/ADCP pathway. Thus, it would be important to quantify the relative contribution of each mechanism to anti-CTLA-4-mediated tumor rejection in future studies. This underscores the importance of tailoring treatment approaches to individual cancer patients. Those with substantial infiltration of CTLA-4-expressing Tregs within tumors may experience a favorable response outcome. Conversely, patients with lower CTLA-4 expression in

Tregs may benefit from avoiding anti-CTLA-4 treatment, as it could potentially interfere with TGF-β signaling and inadvertently facilitate tumor progression.

## STAR ★ METHODS

Detailed methods are provided in the online version of this paper and include the following:

- KEY RESOURCES TABLE
- EXPERIMENTAL MODEL AND STUDY PARTICIPANT DETAILS

`o` Animals
`o` In vitro cell culture
- METHOD DETAILS

`o` Tumor growth and experiment and therapy
`o` Flow sorting and flow cytometric analysis
`o` Glucose-uptake assay and pHrodo Red assay for lactic-acid uptake
`o` Suppression and proliferation assay
`o` RT-qPCR
`o` Immunoblotting
`o` ScRNA-sequencing
`o` Treg scRNA-seq computational analysis

## SUPPLEMENTAL INFORMATION

## ACKNOWLEDGMENTS

This research was supported by the Intramural Research Programs of NIDCR and NHBLI, National Institute of health (NIH), USA. We thank Dr. Arlene Sharp from Harvard Medical School for providing us with the *Ctla4^f/f^* mice. We would like to thank Combined Technical Research Core, Genomics and Computational Biology Core and Veterinary Resource Core at NIDCR for their service and technical assistance.

## AUTHOR CONTRIBUTIONS

N.L. designed and performed the experiments, analyzed data, and drafted the manuscript. J.X., W.W.C, R.K., W.J., T.G., S.L. S.S, L.C.P, Y-J L, V.O and E.N performed the experiments. W.L.K., Y.C analyzed the scRNA-seq data. K.Z provided crucial scientific input, supervised the study and edited the manuscript.

W.J.C conceived and supervised the research, designed the experiments, and wrote the manuscript.

## STAR ★ METHODS

### KEY RESOURCES TABLE

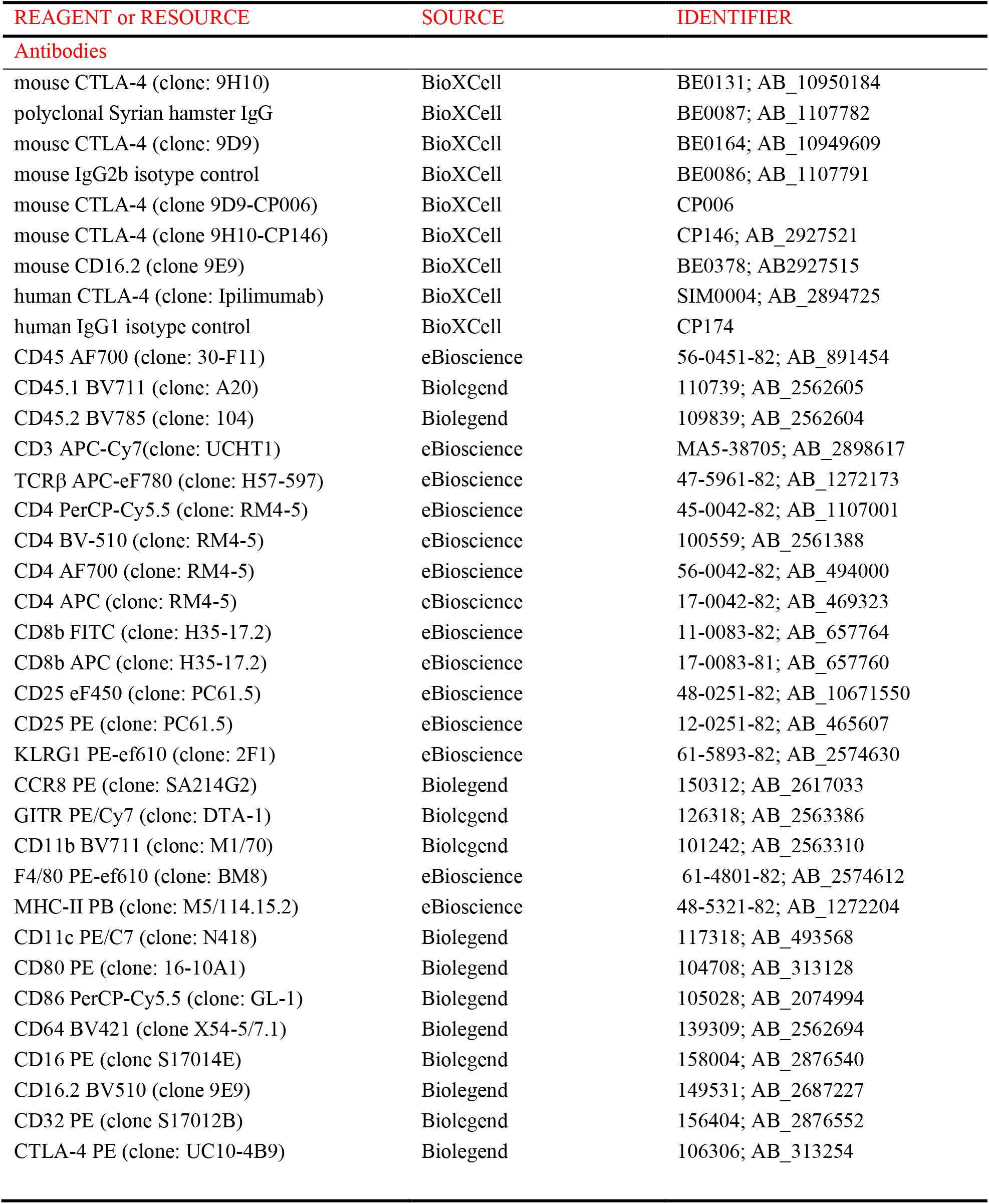
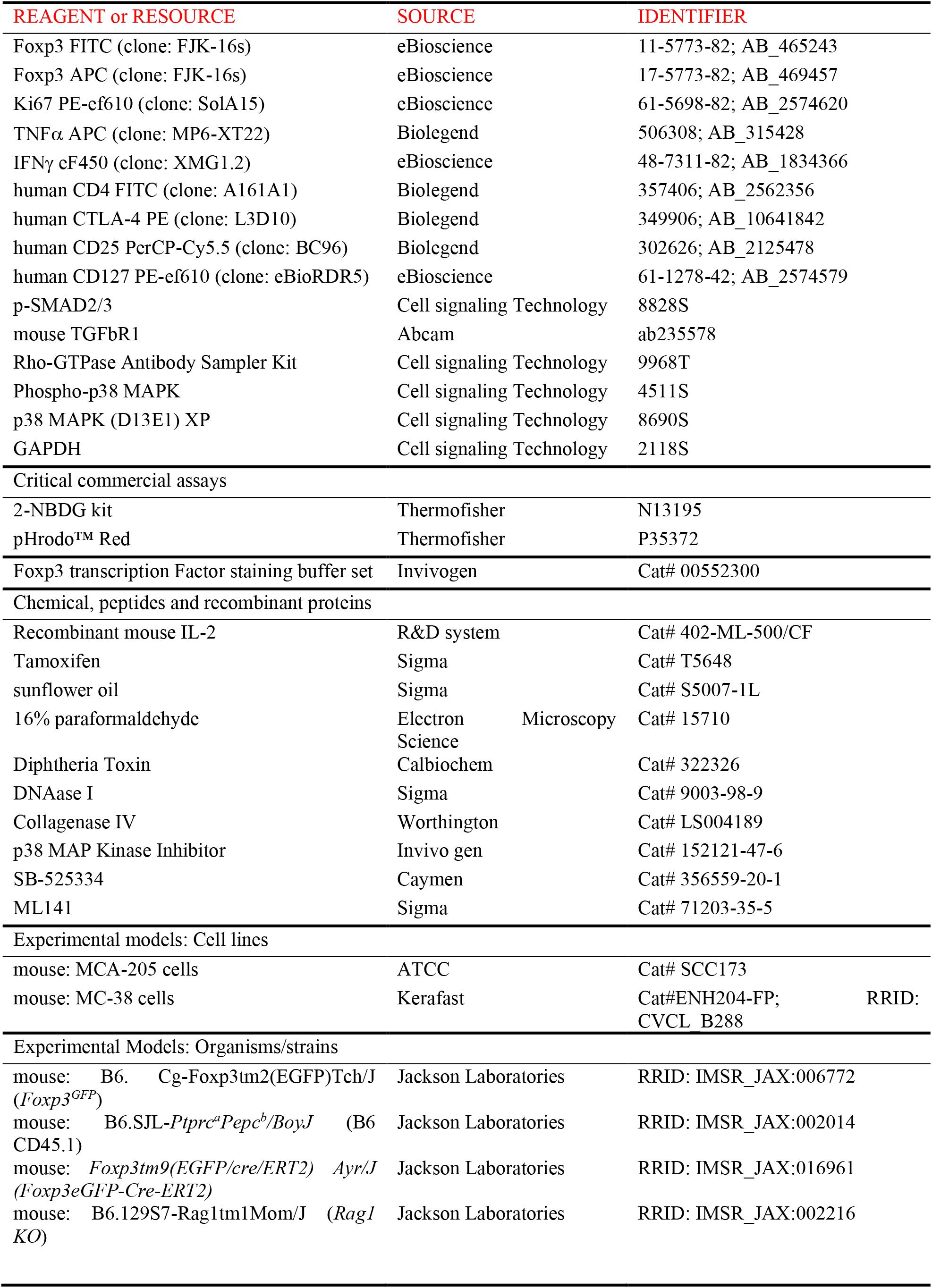
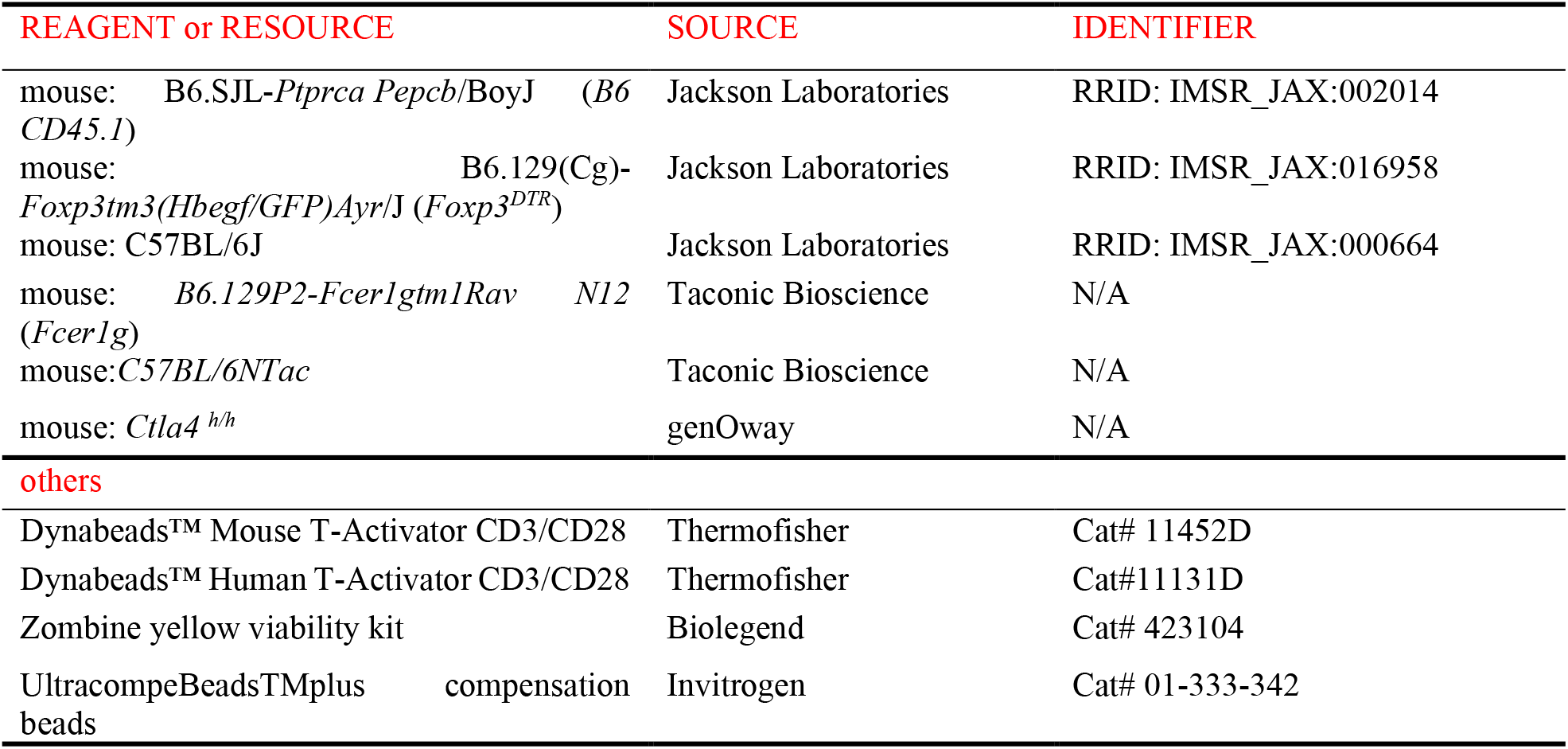

## RESOURCE AVAILABILITY

### Lead contact

Further information and requests should be directed to and will be fulfilled by the lead contact, Dr. Wanjun Chen.

### Materials availability

This study did not generate new unique reagents.

### Data and code availability

scRNA-seq files are deposited in Gene Expression Omnibus (GEO) database under accession number (GSE295085). This data will be publicly available upon the publication of the paper. Other datasets used and analyzed in the current study are available from the corresponding author upon reasonable request.

## EXPERIMENTAL MODELS AND STUDY PARTICIPANT DETAILS

### Animals

All mice used in this study were 8-12 weeks old. CD45.1, C57BL/6 -*Foxp3^GFP^*, *Foxp3^GFP.cre.ERT^*^2^, *Rag1^-/-^*, *Foxp3^DTR^* mice were purchased from Jackson Laboratory. C57BL/6 and α-chain-KO mice (*Fcgr* KO) were purchased from Taconic Bioscience. *Ctla4^h/h^* mice were purchased from genOway company. *Ctla4^f/f^* mice were a gift from Arlene Sharp’s Lab in Harvard University and were used to generate *Ctla4^f/f^ Foxp3^GFP.cre.ERT^*^2^ (termed *Ctla4^f/f^ Foxp3^cre-ERT^*^2^). *Tgfbr1^ER^ ^cre^* mice were generated in house by crossing *Ert2^cre+^* mice with *Tgfbr1^f/f^* mice. *Tgfbr1*- *Foxp3^cre-ERT^*^2^ were generated in house by crossing *Foxp3^GFP.cre.ERT^*^2^ mice with *Tgfbr1^f/f^* mice. Mice were housed in the animal facility of National Institute of Dental and Craniofacial Research (NIDCR) and kept under specific pathogen free conditions. All experimental procedures were approved by the Animal Care and Use Committee (ACUC) of NIDCR. Mice were monitored twice per week so that the maximal tumor size won’t exceed 2.0 cm in diameter.

### In vitro cell culture

MCA-205 cells were purchased from the American Type Culture Collection (ATCC), MC-38 cells were purchased from Kerafast, Both cell lines were cultured in DMEM supplemented with 10% heat-inactivated fetal bovine serum, 2 mM L-glutamine, 1% penicillin plus streptomycin. Cell lines were validated for lack of mycoplasma infection. For in vitro conditioning experiments, isolated Treg or Tcon cells were activated in complete DMEM (10% heat-inactivated fetal bovine serum, 2 mM L-glutamine, 1% penicillin plus streptomycin, non-essential amino acid, 1mM sodium pyruvates, 5mM HEPES and 0.1 mM β-mercaptoethanol). Cells were cultured in 100-mm tissue plates or 96-well round bottom plates in an incubator for 1 day or 3 days with humidifier air and 5% CO2 at 37 °C. For in vitro CTLA-4 engagement experiment, T cells were activated using CD3/28 Dynabeads (Thermofisher) as per the manufacture’s protocol. This activation was carried out in the presence of anti-CTLA-4 or hamster IgG control at a dose of 10 µg/ml, crosslinked with a secondary antibody at dose of 2.5 µg/ml, specifically goat anti-hamster from Jackson ImmunoResearch Laboratory. For the expression of *Klrg1* and *Ccr8* genes, Tregs were cultured under CTLA-4 engagement in presence of tumor-derived conditional medium overnight. For the in vitro CTLA-4 blockade experiment, T cells were cultured with a low dose of soluble anti-CD3 (0.01 µg/ml) and CD28 and treated with anti-CTLA-4 or hamster IgG control for 1 day. For the expression of CD80 and CD86, Tregs were cultured with a low dose of soluble anti-CD3 (0.01 µg/ml) and CD28 for 1 day, and analysis of CD80 and CD86 in Tregs (CD4^+^Foxp3^+^) by flow cytometry. For the human Tregs, healthy donors were obtained from the NIDCR blood bank. T cells were obtained by elutriation and then staining for sort. Tregs were cultured for 1 day in DMEM with 10% heat-inactivated fetal bovine serum, 2 mM L-glutamine, 1% penicillin plus streptomycin, non-essential amino acid, 1mM sodium pyruvates, 5mM HEPES and 0.1 mM β-mercaptoethanol). Human Tregs were activated using human αCD3/28 Dynabeads (Thermofisher) as per the manufacture’s protocol. This activation was carried out in the presence of anti-Ipilimumab or human IgG1 control at a dose of 10 µg/ml, crosslinked with a secondary antibody at dose of 2.5 µg/ml, specifically mouse anti-human IgG from Jackson ImmunoResearch Laboratory. For human Tregs blockade experiment, human Tregs were cultured with low dose of αCD3 (0.01 µg/ml) together with αCD28 (1 µg/ml), in presence of human IgG1 or Ipilimumab overnight.

### Tumor growth experiment and therapy

Eight- to 12 weeks old mice were inoculated subcutaneously on both sides with 5× 10^5^ MCA-205 cells or 2.5× 10^5^ B16-F10 cells in PBS. The experiments were stopped once the tumor reached 20 mm in any direction, or the tumors developed ulcerations. For the tumor-growth experiments in *Foxp3^GFP^*, C57BL/6 and α-chain-KO mice (*Fcgr* KO) and *Ctla4^h/h^* mice, mouse anti-CTLA-4 (9H10 clone, 9D9 clone), mouse anti-CTLA-4 (9H10-CP146 clone, 9D9-CP006 clone), anti-CD16.2 (9E9 clone) and human anti-CTLA4 (Ipilimumab) antibodies or isotype control (Syrian hamster IgG, mouse IgG2b and mouse IgG1), (Armenian hamster IgG) and human IgG1 isotype control (Bio X cell) treatment (200 µg) started on day 7 after tumor inoculation, and harvest mice after treatment for 3 days. For the inducible deletion of *Ctla4* or *Tgfbr1* in Tregs, *Ctla4^f/f^ Foxp3^cre-ERT^*^2^ mice and *Tgfbr1*- *Foxp^cre-ERT^*^2^ mice were injected intraperitoneally with 1mg of tamoxifen (T5648, Sigma) or sunflower oil from day -4 to day 0. On day 0, mice were inoculated subcutaneously with 5× 10^5^ MCA-205 cells in PBS at both sides. Tamoxifen treatment continued twice weekly until the endpoint of the experiment. On day 7, mice were administrated with 200 µg anti-CTLA-4 or Syrian hamster IgG isotype control once until collection. MAC-205 tumor-bearing *Foxp3^DTR^* mice were intraperitoneally injected with DT at dose 500 ng/each mouse, then followed the above protocol to treat with anti-CTLA-4 antibody. Tregs depletion level was accessed by flow cytometry by quantifying Foxp3^+^CD4^+^ T cells.

### Flow sorting and cytometric analysis

Single-cell suspension from mouse tumors (day 10 or 13 post tumor inoculation) were prepared for flow sorting and cytometric analysis as described previously ^71^. In brief, tumors were gently minced with scissors to small pieces less than 1mm in size. Afterwards, tumor tissues were placed in cold serum-free DMDM medium with Collagenase IV (0.2mg/ml; Gibco) and DNase (0.1 mg/ml; Sigma-Aldrich) and then incubated in a shaking incubator at 37 °C for 45 mins at 200 r.p.m. Specimens were passed through 70-μm strainers and centrifuged at 350g for 5 mins. Tumor-infiltrating lymphocytes were separated by gradient centrifugation with 40%/80% Percoll. Cell pellets were resuspended, and cell labeling was performed after blocking FcγR III/II with an antibody to CD16/32 (93, catalogue number 101320, BioLegend, dilution 1:100) Prior to flow sorting, tumor cell suspension and splenocytes were stained with Zombie yellow (RUO, catalogue number 423104, BioLegend, dilution 1:300 in PBS), anti-CD45 (30-F11, catalogue number 56-0451-82, eBioscience, dilution 1:200), TCRβ (H57-597, catalogue number 47-5961-82, eBioscience, dilution 1:200), CD4 (RM4-5, catalogue number 45-0042-82, BioLegend, dilution 1:200), CD8β (H35-17.2, catalogue number 225-0083-82, eBioscience, dilution 1:200), CD25 (PC61.5, catalogue number 48-0251-82, eBioscience, dilution 1:200), Gitr (DTA-1, catalogue number 126318, BioLegend, dilution 1:200). For the human Tregs sort, requested PBMCs were stained with: anti-human CD4 (A161A1, catalogue number 357406, BioLegend, dilution 1:200), anti-human CD25 (BC96, catalogue number 302626, BioLegend, dilution 1:200), anti-human CD127 (eBioRDR5, catalogue number 61-1278-42, eBioscience, dilution 1:200). FACS-sorting was performed on the BD FACSAria.

Staining for expression markers of cytometric analysis was performed with anti-mouse specific antibodies including anti- CD45.1 (A20, catalogue number 110739, BioLegend, dilution 1:200), CD45.2 (104, catalogue number 109839, BioLegend, dilution 1:200), CD3 (17A2, catalogue number 47-0032-82, eBioscience, dilution 1:200), CD4 (RM4-5, catalogue number 100559, BioLegend, dilution 1:200), CD4 (RM4-5, catalogue number 56-0042-82, eBioscience, dilution 1:200), CD4 (RM4-5, catalogue number 17-0042-82, eBioscience, dilution 1:200), CD8β (H35-17.2, catalogue number 11-0083-82, eBioscience, dilution 1:200), CD25 (PC61.5, catalogue number 45-0251-82, eBioscience, dilution 1:200), KLRG1 (2F1, catalogue number 61-5893-82, eBioscience, dilution 1:200), CCR8 (SA214G2, catalogue number 150311, BioLegend, dilution 1:200), NRP1 (3E12, catalogue number 145205, BioLegend, dilution 1:200), Gitr (DTA-1, catalogue number 126316, BioLegend, dilution 1:200), CD11b (M1/70, catalogue number 101242, BioLegend, dilution 1:200), F4/80 (BM8, catalogue number 61-4801-82, eBioscience, dilution 1:200), CD11c (N418, catalogue number 117318, BioLegend, dilution 1:200), MHC-II (M5/114.15.2, catalogue number 48-5321-82, eBioscience, dilution 1:200), CD80 (16-10A1, catalogue number 104708, BioLegend, dilution 1:200), CD86 (GL-1, catalogue number 105028, BioLegend, dilution 1:200), CD64 (FcγRⅠ) (X54-5/7.1, catalogue number 139309, BioLegend, dilution 1:200), CD16 (FcγRⅢ) (S17014E, catalogue number 158004, BioLegend, dilution 1:200), CD16.2 FcγⅣ (9E9, catalogue number 149531, BioLegend, dilution 1:200), CTLA-4 (UC10-489, catalogue number 106305, BioLegend, dilution 1:200), FoxpP3 (FJK-16s, catalogue number 11-5773-82, eBioscience, dilution 1:200), Ki67 (SolA15, catalogue number 61-5698-82, eBioscience, dilution 1:200),TNF-α (MP6-XT22, catalogue number 506308, BioLegend, dilution 1:200), IFN-γ (XMG1.2, catalogue number 505818, BioLegend, dilution 1:200), anti-human specific antibody: anti-CTLA-4 (L3D10, catalogue number 349906, BioLegend, dilution 1:200). Intracellular staining was performed using the Foxp3 fix/perm buffer set (eBioscience) according to the manufacture’s protocol, and staining was performed overnight at 4°C. Stained cells were analyzed on LSRFortessa (BD). Cell doublets and dead cells were excluded from analysis by assessing side-scatter and forward-scatter width to area, and by utilizing zombie yellow staining. Flow cytometric data and proliferation modelling were analyzed with FlowJo version 10 software (Tree Star).

### Glucose-uptake assay and pHrodo Red assay for lactic-acid uptake

Single-cell suspension from tumors were put in PBS containing 100 µM 2NBDG for 25 mins at 37°C. For the lactic-acid uptake, cell suspensions from tumor tissues were loaded with pHrodo Red AM (ThermoFisher Scientific) according to the maufacturer’s protocol with 20 mM HEPES in PBS. Cell surface staining for multicolor flow cytometry was performed after the normal protocol in 20 mM HEPES/PBS. At the cytometry, lactic acid was added into each sample at final concentration of 2.5 mM. Samples were running at 0 mins and 30 mins after supplement of lactic acid.

### Suppression and proliferation assay

In vitro suppression assay was performed as described. In brief, Tregs (CD4^+^Foxp3^+^) cell populations were isolated by flow-assisting sorting from tumors or spleens of C57BL/6 -*Foxp3^EGFP^* mice, Responder cells (CD4^+^Foxp3^-^) and antigen-presenting cells were isolated from the spleens of CD45.1 mice. Tregs were cocultured at 37 °C for 72 hrs with irradiated APCs and CellTrace Violet-labeled responder cells at ratios of 1:4 (Treg: responder) in complete DMEM medium with 0.5 µg/ml anti-CD3 antibodies. The proliferation of responding cell populations was modelled using FlowJo Version 10.

### RT-qPCR

Total RNA was extracted from cultured cells or sorted cells with a RNeasy Mini Kit (Qiagen), and complementary DNA was transcribed using a high-capacity cDNA reverse transcription kit (Applied Biosystems). Quantitative real-time PCR was performed according to the protocol of TaqMan gene expression master (Applied Biosystem) using a QuantStudio 3 Real-Time PCR Systems (Applied Biosystem) with the following primers: *Hprt* (Mm00446968_m1), *Il2* (Mm00434256_m1), *Il2ra* (Mm01340213_m1), *Tgfbr1* (Mm00436964_m1), *Tgfbr2* (Mm03024091_m1), *pro-Tgf*β*r1* (Rn00688966_m1), *Foxp3* (Mm00475162_m1 ), *Ctla4* (Mm00486849_m1), *Klrg1* (Mm00516879_m1), *Ccr8* (Mm99999115_s1), *Slc2a1* (Mm00441480_m1), *Hk2* (Mm00443385_m1), *Ldha* (Mm01612132_g1), *Slc16a3* (Mm00446102_m1), *HPRT* (Hs02800695_m1), *TGFBR1* (Hs00610320_m1), *TGFBR2* (Hs00234253_m1), *IL2RA* (*Hs00907777_m1*)*, LDHA* (Hs01378790_g1)*, SLC16A3* (Hs00358829_m1)*, HK2* (Hs00606086_m1). qPCR data were analyzed by ddCt method by normalizing the expression of each gene to HPRT and then to the control group.

### Immunoblotting

The protein samples were extracted in ice-cold RIPA lysis buffer. Total protein was quantified using the assay kit according to the manufacture’s instruction (BioRad). Immunoblotting was performed as described previously ^71^, 10% polyacrylamide gel (SDS-Page, Invitrogen) were equiloaded with 10 µg of protein and transferred onto polyvinylidene difluoride (PVDF) membranes (Millipore). After blocking with 5% nonfat milk in TBST (0.5 M NaCl, Tris-HCl and 0.5% Tween-20) for 1 hr at room temperature and membranes were probed with primary antibodies (all antibodies used for western blotting in this paper at a dilution of 1:1000) in 5% BSA in TBST overnight at 4 °C. The membranes were washed with TBST buffer and then incubated with secondary antibodies (1:5000 dilution) for 1 hr. The results were visualized by enhanced chemiluminescence using an Amersham Imager (Cytiva) according to the manufacturer’s instruction.

### scRNA-sequencing

After tumor digestion, cells were stained with Zombie yellow in a light-protected environment at room temperature for 10 mins to exclude dead/dying cells. Subsequently, the samples were washed and incubated with anti-CD16/32 in FCAS buffer at 4 °C to block binding to Fc receptors. Surface markers were stained for 30 mins at 4 °C in sorting buffer. Live, CD45^+^TCRβ^+^CD4^+^Foxp3^+^ cells were then sort purified. Cells were sorted into PBS with 2% FBS. Single-cell gene expression analysis was performed with the 10x genomics system using Chromium Next GEM single cell 3’ GEM Reagent kit (catalogue number 1000121) and chip G single cell kit (catalogue number 1000120) following the user guide for the Chromium Next GEM Single Cell 3’ Reagent Kits v3.1. In brief, >20,000 cells (viability 96%) were lysed for 3 mins and resuspended in Diluted Nuclei Buffer (10x Genomics, 2000207). After the transportation reaction, nuclei were encapsulated and barcoded. Next-generation sequencing libraries were constructed following the manufacturer’s instruction and were sequenced on an Illumina NovaSeq 6000 system.

### Treg scRNA-seq computational analysis

We processed 10X single-cell RNA-seq (scRNA-seq) reads using Cell Ranger (v7.0.1) on the mm10-2020-A mouse reference. We generated gene-barcode matrices for each sample and imported them into R using Seurat (v4.2.1). For quality control, we retained cells with ≥500 and <6000 detected genes and excluded cells with >10% mitochondrial gene expression. We did not explicitly filter on ribosomal content, nor did we remove specific highly expressed genes (e.g., Malat1, Tmsb4x, Actb). Immunoglobulin and mitochondrial genes were excluded from the set of variable features to avoid bias. We normalized the data using Seurat’s “LogNormalize” method (scale factor = 10,000), identified the top 2,000 variable features with the “vst” method, and scaled the data while regressing out unwanted sources of variation. Samples from IgG- and CTLA4-treated conditions were subsequently integrated using a canonical correlation analysis (CCA)-based workflow (Seurat’s IntegrateLayers and CCAIntegration methods) to correct for batch effects. Dimensionality reduction was performed with principal component analysis (PCA), followed by visualization with uniform manifold approximation and projection (UMAP). We defined four gene sets (Activated_effector, Proliferating, IFN_response, Naive_Resting). Using AddModuleScore, it calculates a module score for each of these gene sets for every single cell. Each cell is then assigned to the group for which it has the highest module score. For differential expression analysis between conditions, we used the Wilcoxon rank-sum test on genes detected in at least 20% of cells in either group and considered only those with a fold change ≥1.2.

### Statistical analysis

Data presented in the figures are mean ± s.e.m. Multiple groups comparison in vivo and ex vivo assay were accomplished by one or two-way ANOVA. For single comparison, statistical significance was determined by the Student’s *t* test. P values < 0.05 were significant. Significance for the GSEA analysis and the differential gene expression based on scRNA-seq was determined using the Wilcoxon rank sum test with Bonferroni multiple-comparison correction. All analysis was completed with Prism V8 software (GraphPad).

## Supplementary Figure Legends

**Figure S1.**
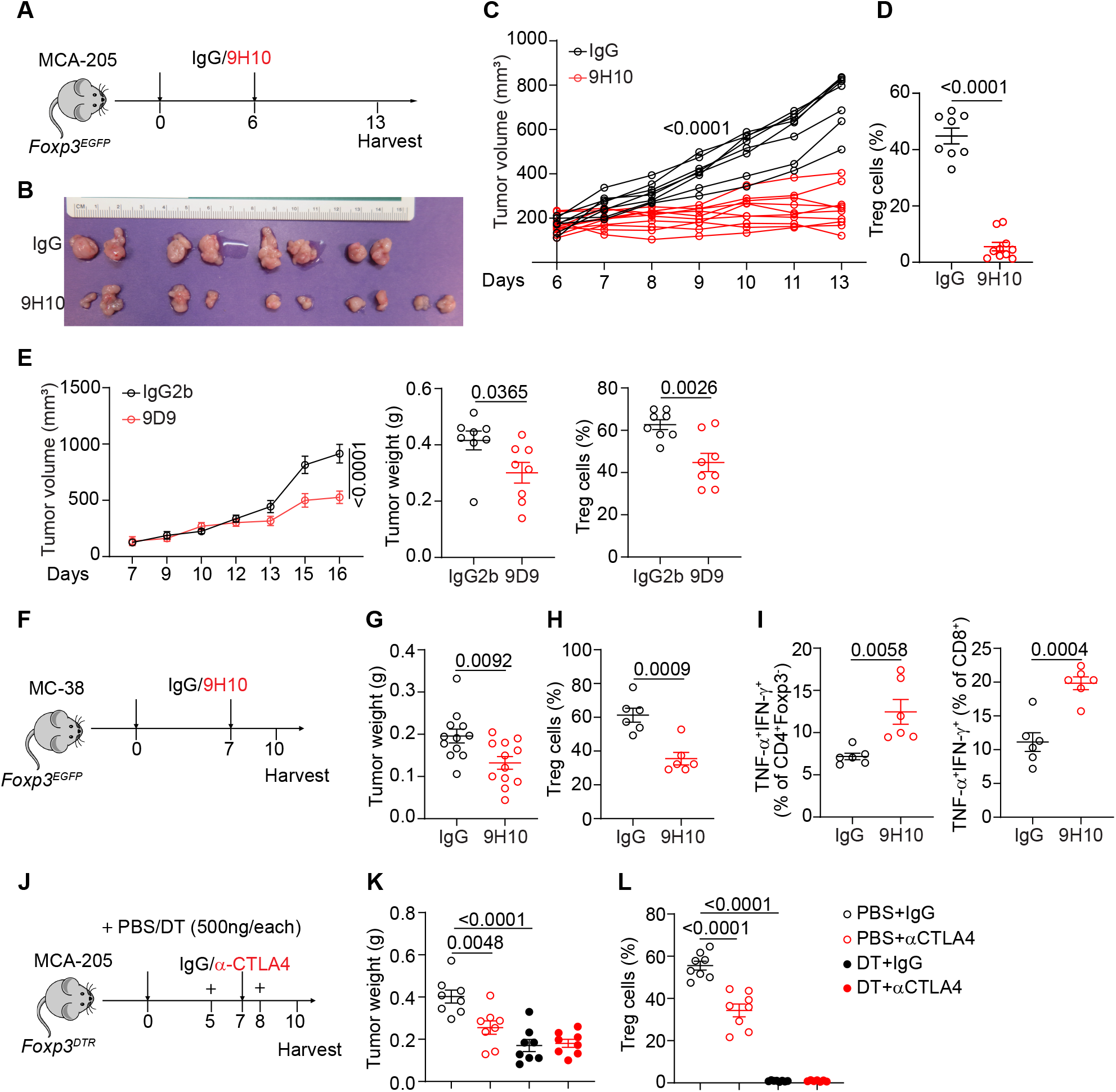
Reduction of intratumor Tregs is required for αCTLA-4 mediated tumor rejection. Related to Figure 1. (A) Experimental design for anti-CTLA-4 treatment on mice bearing MCA-205 tumors. (B-C) Tumor picture (B) and growth curves (C) from mice bearing MCA-205 tumors treated with IgG or 9H10 antibody. (D) Percentage of Tregs (CD4^+^Foxp3^+^) among CD4^+^ T cells from tumors. (E) Growth curves, tumors weight and Tregs frequency from mice bearing MCA-205 tumors treated with Isotype mouse IgG2b or 9D9 antibody. (F-G) Experimental design for anti-CTLA-4 treatment on mice bearing MC-38 tumors (F) and tumor weight between IgG vs. αCTLA-4 treatment (G). (H) Percentage of Tregs (CD4^+^Foxp3^+^) among CD4^+^ T cells from tumors. (I) Percentage of TNF-α^+^IFN-γ^+^ producing Th cells or CD8^+^ T cells in tumors. (J) Schematic for Tregs depletion on *Foxp3^DTR^* mice. (K) Tumors weight from indicated groups. (L) Percentage of Tregs (CD4^+^Foxp3^+^) among CD4^+^ T cells. Data is representative of two independent experiments with similar results. Significance was determined by unpaired two-tailed *t*-test (C-I) or one-way analysis of variance (ANOVA) with Tukey’s multiple comparison (K-L). Data shown as mean ± s.e.m.

**Figure S2.**
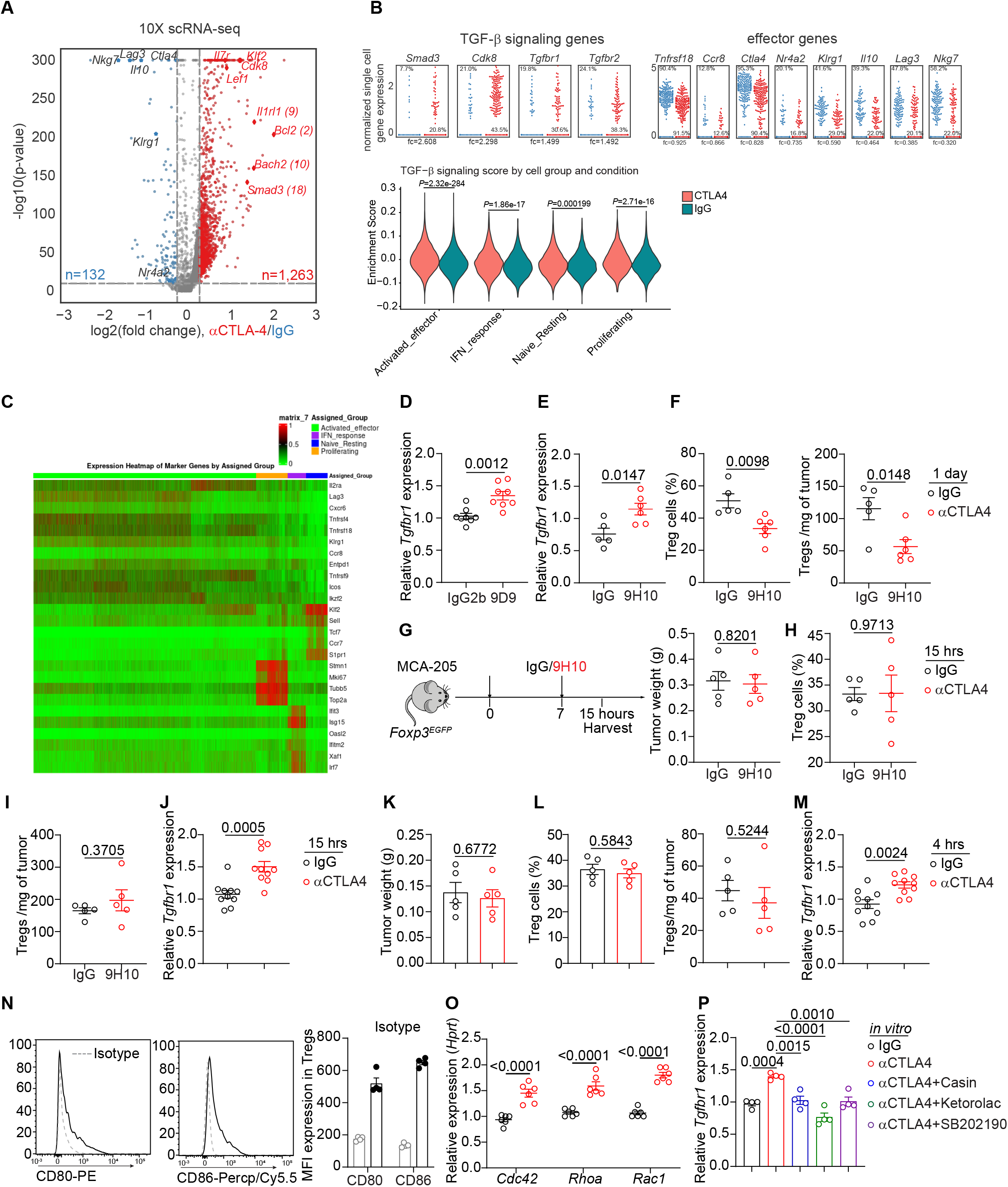
Analysis of Tregs from tumor tissues. Related to Figure 1. (A) Volcano plot of genes presents the magnitude (log2 [fold change], x axis) and significance (−log10 [adjusted p value], y axis) for Tregs from IgG control or αCTLA-4 treatment from scRNA-seq. (B) Dot plots showing differences in expression of genes related to TGF-β signaling and effector status in intratumor Tregs from scRNA-seq dataset, and Violin plots showing the distribution of the TGF-β signature score, grouped by computationally-defined cell states and split by condition (control vs. anti-CTLA-4-treated). (C) Heatmap of differentially expressed genes to identify different subcluster in tumor-infiltrating Tregs from *Foxp3^EGFP^* mice treated with IgG control or αCTLA-4 for 3 days. (D) RT-PCR analysis of *Tgfbr1* mRNA in intratumor Tregs from mice bearing MCA-205 tumors treated with murine IgG2b or 9D9. (E) Expression of *Tgfbr1* mRNA in intratumor Tregs from mice bearing MCA-205 tumors treated with IgG or αCTLA-4 for 1 day. (F) Left, percentage of Tregs (CD4^+^Foxp3^+^) among CD4^+^ T cells from tumors, and right, Tregs number per mg of tumor in mice bearing MCA-205 tumors treated with IgG or αCTLA-4 for 1 day. (G) Left, Experimental design for anti-CTLA-4 treatment for 15 hours on mice bearing MCA-205 tumors and right, tumors weight from IgG or 9H10 antibody treated mice. (H) Percentage of Tregs (CD4^+^Foxp3^+^) in tumors. (I) Tregs number per mg of tumor in mice. (J) Expression of *Tgfbr1* mRNA in tumor-infiltrating Tregs from mice bearing MCA-205 tumors treated with IgG or αCTLA-4 for 15 hours. (K) tumors weight from IgG or 9H10 antibody treated mice for 4 hours. (L) Left, Percentage of Tregs (CD4^+^Foxp3^+^) in tumors and right, Tregs number per mg of tumor in mice. (M) Expression of *Tgfbr1* mRNA in tumor-infiltrating Tregs from mice bearing MCA-205 tumors treated with IgG or αCTLA-4 for 4 hours. (N) Quantification of CD80 and CD86 MFI in Tregs in presence of αCD3 (0.01µg/ml) and αCD28 (1µg/ml) overnight. (O) Expression of Rho-GTPase associated genes in Tregs cultured under CTLA-4 engagement conditions (IgG, isotype control antibody for αCTLA-4). (P) Expression of *Tgfbr1* mRNA in Tregs cultured under CTLA-4 engagement conditions in presence of Rho GTPase inhibitors (Casin: 1 µM; Ketorolac: 1 µM) and p38 inhibitor (SB202190: 0.5 µM) for overnight. Data is representative of two independent experiments with similar results or pooled from three independent experiments with similar results. Significance was determined by unpaired two-tailed *t*-test (D-O) or one-way analysis of variance (ANOVA) with Tukey’s multiple comparison (P). Data are shown as mean ± s.e.m.

**Figure S3.**
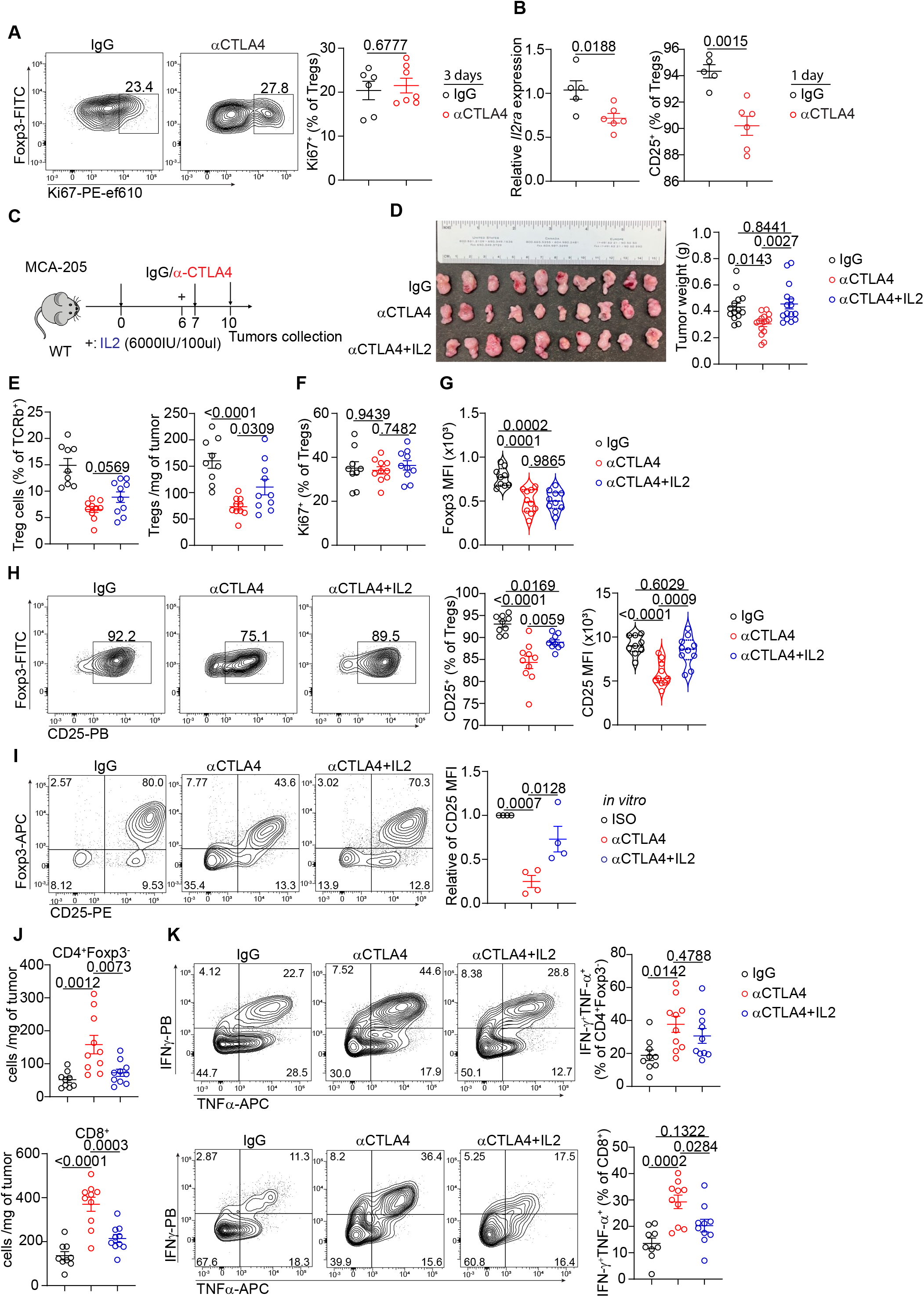
IL-2 administration reverses anti-CTLA4-mediated tumor repression by increasing tumor-infiltrating Tregs. Related to Figure 2. (A) Flow cytometry panels of Ki67 staining in tumor-infiltrating Tregs from mice bearing MCA-205 tumors treated with IgG or αCTLA-4 for 3 days. (B) Left, Expression of *Il2ra* in tumor-infiltrating Tregs by mRNA and right, percentage of CD25 expression in Tregs from tumors treated with IgG or αCTLA-4 for 1 day. (C) Schematic for administration of αCTLA-4 and recombinant IL-2 antibody. (D) Left, Tumor picture and right, tumor weight from indicated groups. (E) Left, Percentage of Tregs (CD4^+^Foxp3^+^) among TCRβ^+^ T cells from tumors and right, Tregs number per mg of tumor in mice. (F) Percentage of Ki67 staining in tumor-infiltrating Tregs from indicated groups. (G) Quantification of Foxp3 MFI in tumor-infiltrating Tregs. (H) Left, Representative flow plot of CD25 expression by tumor-infiltrating Tregs and right, analysis of CD25 expression in tumor-infiltrating Tregs from mice treated with IgG, αCTLA-4 or αCTLA-4 plus IL-2. (I) *Ex vivo* CD25 expression in Tregs cultured under CTLA-4 engagement, with or without IL-2. (J)Th and CD8^+^ T cells number per mg of tumor in mice from indicated groups. (K) Percentage of TNF-α^+^IFN-γ^+^ producing Th cells or CD8^+^ T cells in tumors. Data in (A-B) are representative of two independent experiments with similar results. Data is pooled from two (C-H, J-K) or three (I) independent experiments with similar results. Significance was determined by unpaired two-tailed *t*-test (A-B) or one-way analysis of variance (ANOVA) with Tukey’s multiple comparison (D-K). Data shown as mean ± s.e.m.

**Figure S4.**
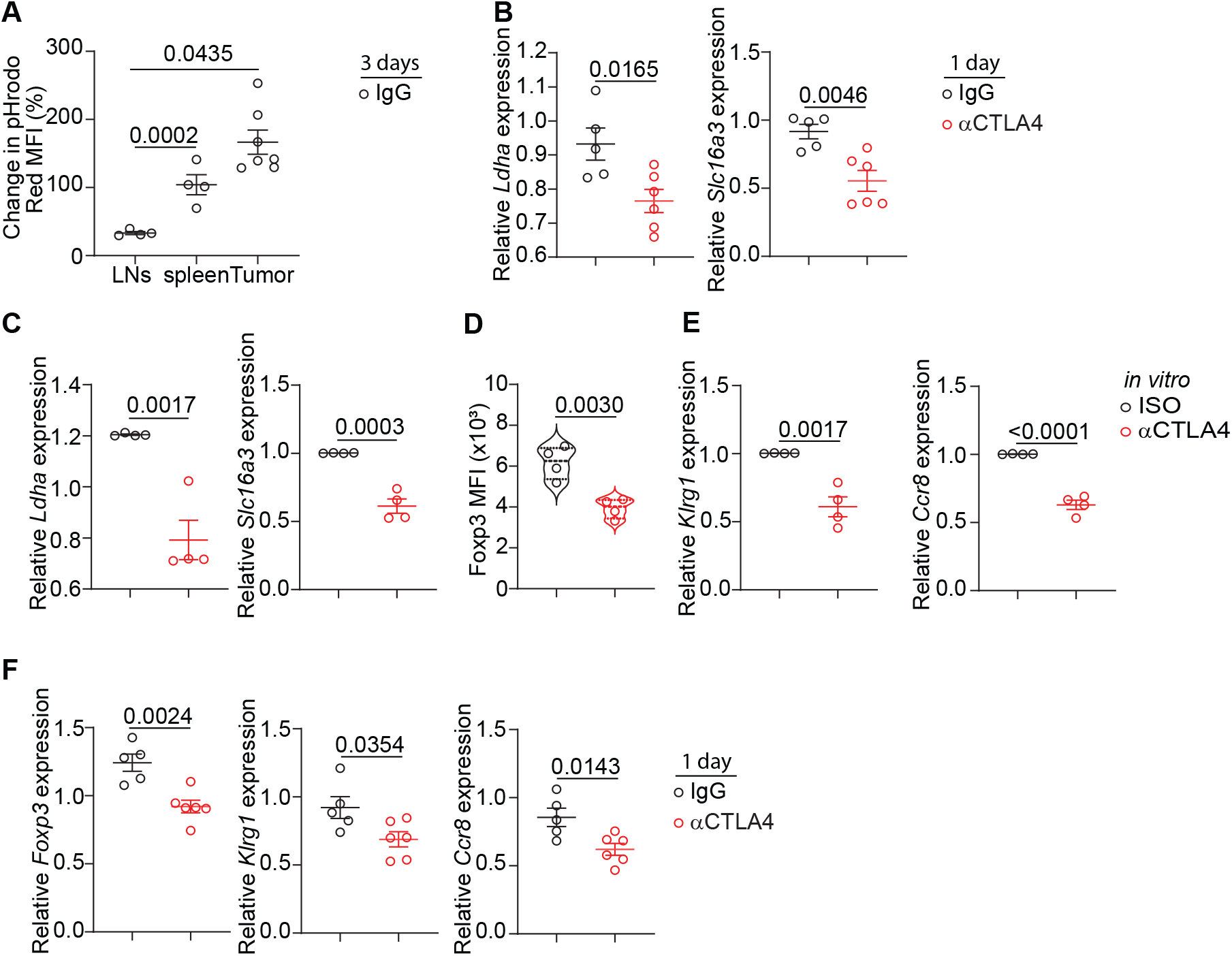
Anti-CTLA-4 treatment drives instability in tumor-infiltrating Tregs. Related to Figure 2. (A) Measurement of lactic acid uptake by Tregs infiltrating tumors and periphery lymphoid tissues. (B) RT-PCR analysis the expression of *Ldha* and *Slc16a3* in tumor-infiltrating Tregs treated with IgG or αCTLA-4 for 1 day. (C) Expression of *Ldha* and *Slc16a3* in Tregs cultured under CTLA-4 engagement condition. (D) Quantification of Foxp3 MFI in cultured Tregs under CTLA-4 engagement for 3 days *ex vivo*. (E) Expression of *Klrg1* and *Ccr8* in Tregs cultured under CTLA-4 engagement condition in presence with tumor-derived medium for 1 day. (F) Expression of *Foxp3*, *Klrg1* and *Ccr8* in tumor-infiltrating Tregs from mice bearing MCA-205 tumors received IgG or αCTLA-4 treatment for 1 day. Data is representative of two independent experiments with similar results. Significance was determined by one-way analysis of variance (ANOVA) with Tukey’s multiple comparison (A) or unpaired two-tailed *t*-test (B-F). Data shown as mean ± s.e.m.

**Figure S5.**
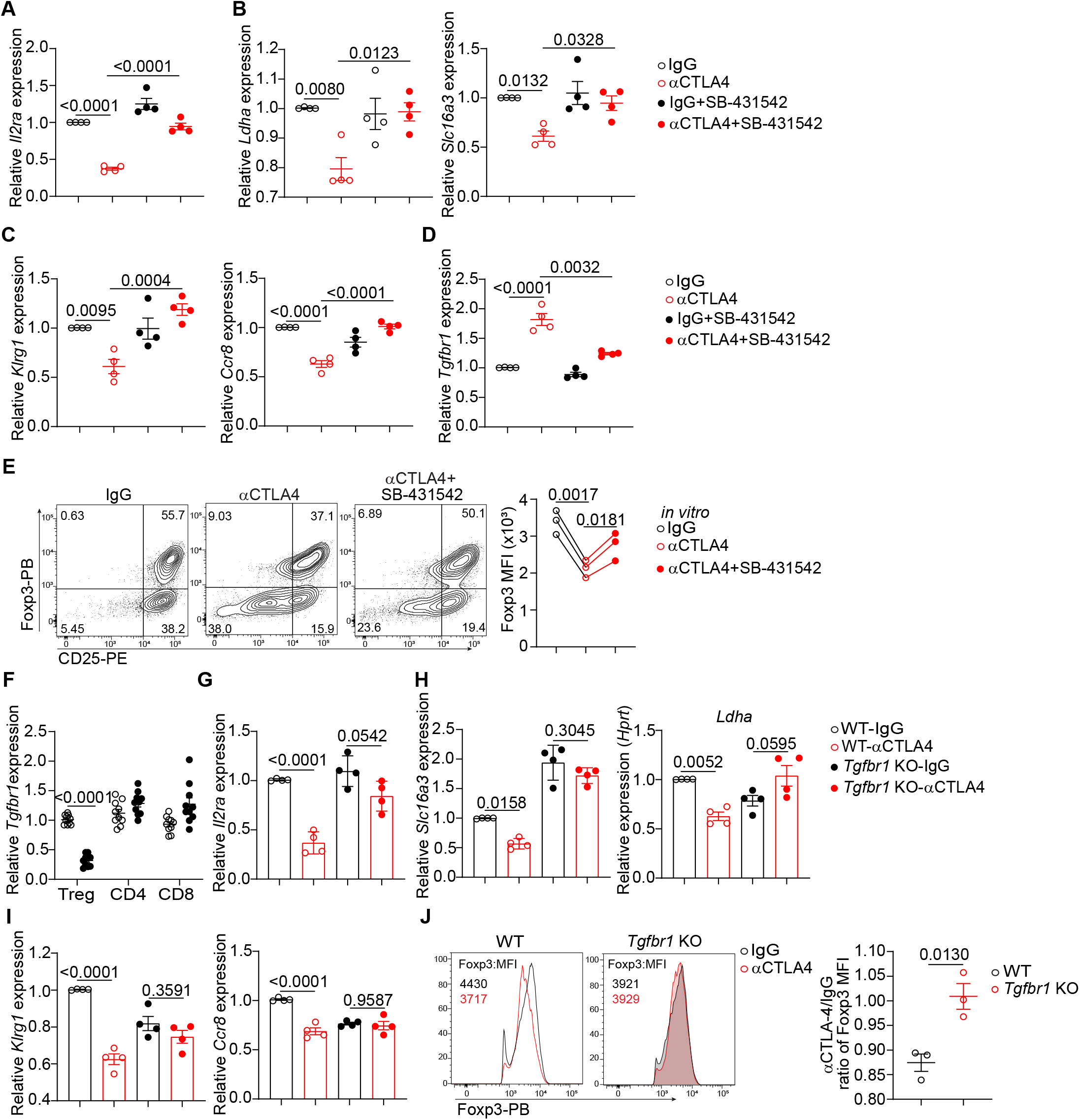
CTLA-4 engagement-mediated Treg dysfunction is dependent on the upregulation of TGF-β signaling *in vitro*. Related to Figure 3. (A-D) Expression of *Il2ra* (A), *Ldha*, *Slc16a3* (B), *Klrg1*, *Ccr8* (C) and *Tgfbr1* (D) in Tregs cultured under CTLA-4 engagement condition with or without the TβRⅠ inhibitor SB-431542 (5 µm). (E) Left, Representative flow plot of Foxp3 and CD25 expression in Tregs cultured under CTLA-4 engagement condition with or without the TβRⅠ inhibitor SB-431542 (5 µm) for 3 days and right, quantification of Foxp3 MFI in cultured Tregs treated as left. (F) Expression of *Tgfbr1* in Tregs, Tcon and CD8^+^ T cells isolated from the spleen and LN of *Tgfbr1^f/f^ Foxp3^cre-ERT^*^2^ mice treated with oil or tamoxifen. (G-I) RT-PCR analysis of the expression of *Il2ra* (G), *Ldha* and *Slc16a3* (H), *Klrg1* and *Ccr8* (I) in Tregs from *Tgfbr1^f/f^ Foxp3^cre-ERT^*^2^ mice treated with oil or tamoxifen and cultured under CTLA-4 engagement condition for overnight. (J) Left, Representative flow histogram of Foxp3 expression in WT and *Tgfbr1* ko Tregs cultured under CTLA-4 engagement for 3 days and right, quantification of Foxp3 MFI in Tregs, showing the relative change with respect to WT-IgG treatment. Data is pooled from three independent experiments with similar results. Significance was determined by unpaired two-tailed *t*-test (J) or one-way analysis of variance (ANOVA) with Tukey’s multiple comparison (A-I). Data shown as mean ± s.e.m.

**Figure S6.**
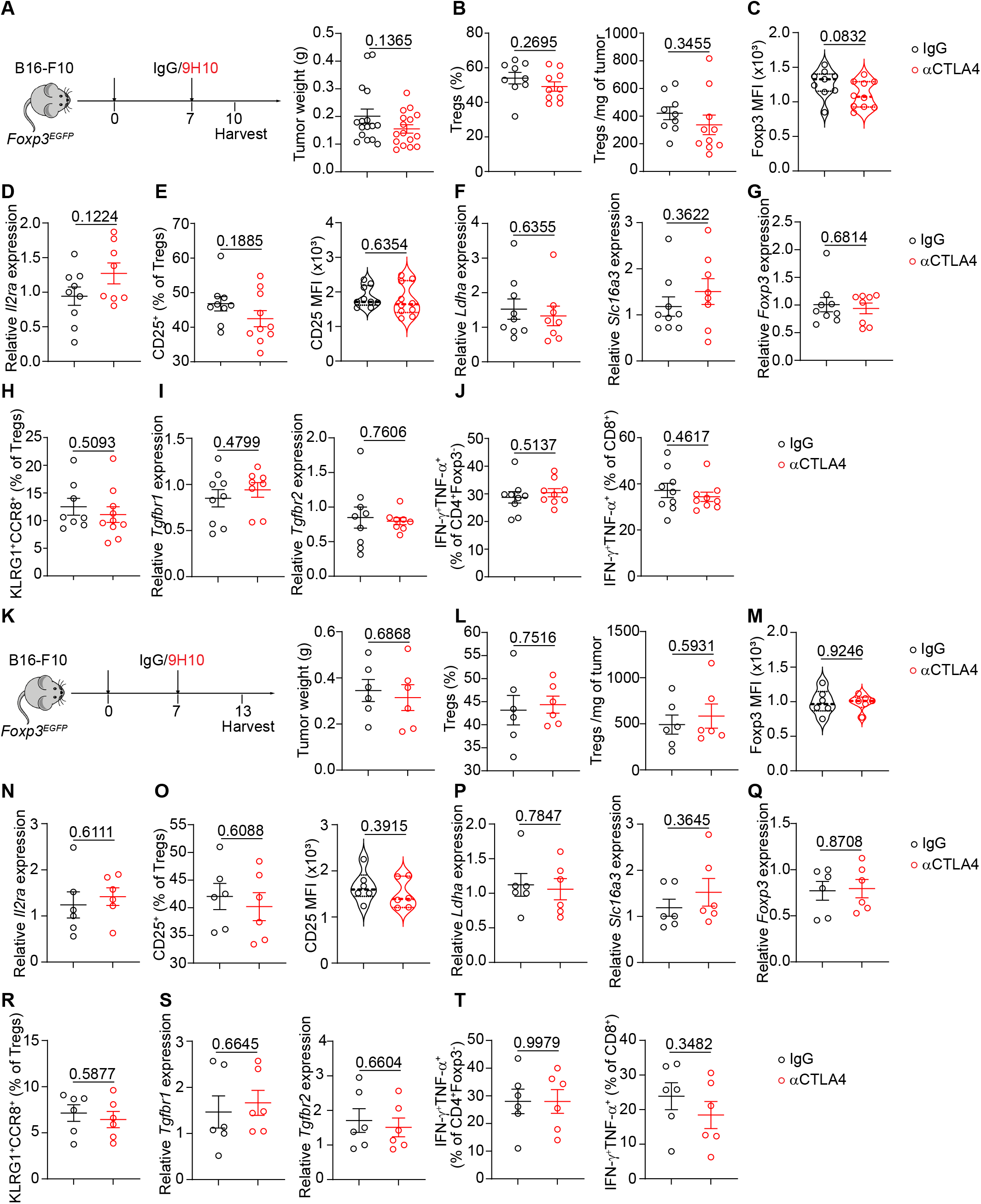
αCTLA-4 therapy failed to up-regulate TGF-β signaling to destabilize Tregs in melanoma tumors, preventing anti-tumor immunity. Related to Figure 3. A-J, Anti-CTLA-4 antibody treatment for 3 days. ( A) Left, Diagram showing the experimental procedure, and right, tumor weight as indicated. (B) Left, Percentage of Tregs (CD4^+^Foxp3^+^) among total CD4^+^ T cells from tumors and right, Tregs number per mg of tumor in mice bearing melanoma B16-F10 cells with αCTLA-4 therapy. ( C) Quantification of Foxp3 MFI in intratumor Tregs from IgG vs. αCTLA-4 treatments. (D) RT-PCR analysis of *Il2ra* in intratumor Tregs from IgG vs. αCTLA4 treatment. (E) Left, percentage of CD25^+^ cells among Tregs from tumors, and right, quantification of CD25 MFI in intratumor Tregs from IgG vs. αCTLA-4 treatment. (F) Expression of genes related to lactate metabolism in isolated intratumor Tregs after IgG vs. αCTLA4 treatment. (G) RT-PCR analysis of *Foxp3* in intratumor Tregs. (H) Analysis of the percentage of KLRG1^+^CCR8^+^ in intratumor Tregs treated with IgG or αCTLA4 antibodies. (I) Expression of *Tgfbr1* and *Tgfbr2* mRNA in intratumor Tregs by RT-PCR from IgG group or αCTLA-4 group. (J) Percentage of TNF-α^+^ IFN-γ^+^ producing CD4^+^Fxop3^-^ cells or CD8^+^ T cells in tumors between IgG vs. αCTLA-4 treatments. K-T, Anti-CTLA-4 treatment for 6 days. (K) Left, Experimental design for anti-CTLA-4 treatment on mice bearing B16-F10 tumors and right, tumor weight as indicated. (L) Left, Percentage of Tregs (CD4^+^Foxp3^+^) among total CD4^+^ T cells from tumors and right, Tregs number per mg of tumor in mice with αCTLA-4 therapy for 6 days. ( M) Quantification of Foxp3 MFI in intratumor Tregs from IgG vs. αCTLA-4 treatments. (N) RT-PCR analysis of *Il2ra* in intratumor Tregs from IgG vs. αCTLA4 treatment for 6 days. (O) Left, percentage of CD25^+^ cells among Tregs from tumors, and right, quantification of CD25 MFI in intratumor Tregs from IgG vs. αCTLA-4 treatments. (P) Expression of genes related to lactate metabolism in isolated intratumor Tregs after IgG vs. αCTLA4 treatment. (Q) RT-PCR analysis of *Foxp3* in intratumor Tregs. (R) Analysis of the percentage of KLRG1^+^CCR8^+^ in intratumor Tregs treated with IgG or αCTLA4 antibodies. (S) Expression of *Tgfbr1* and *Tgfbr2* mRNA in intratumor Tregs by RT-PCR from IgG group or αCTLA-4 group. (T) Percentage of TNF-α^+^ IFN-γ^+^ producing CD4^+^Fxop3^-^ cells or CD8^+^ T cells in tumors between IgG vs. αCTLA-4 treatments. Data is representative of two independent experiments with similar results. Significance was determined by unpaired two-tailed *t*-test. Data shown as mean ± s.e.m.

**Figure S7.**
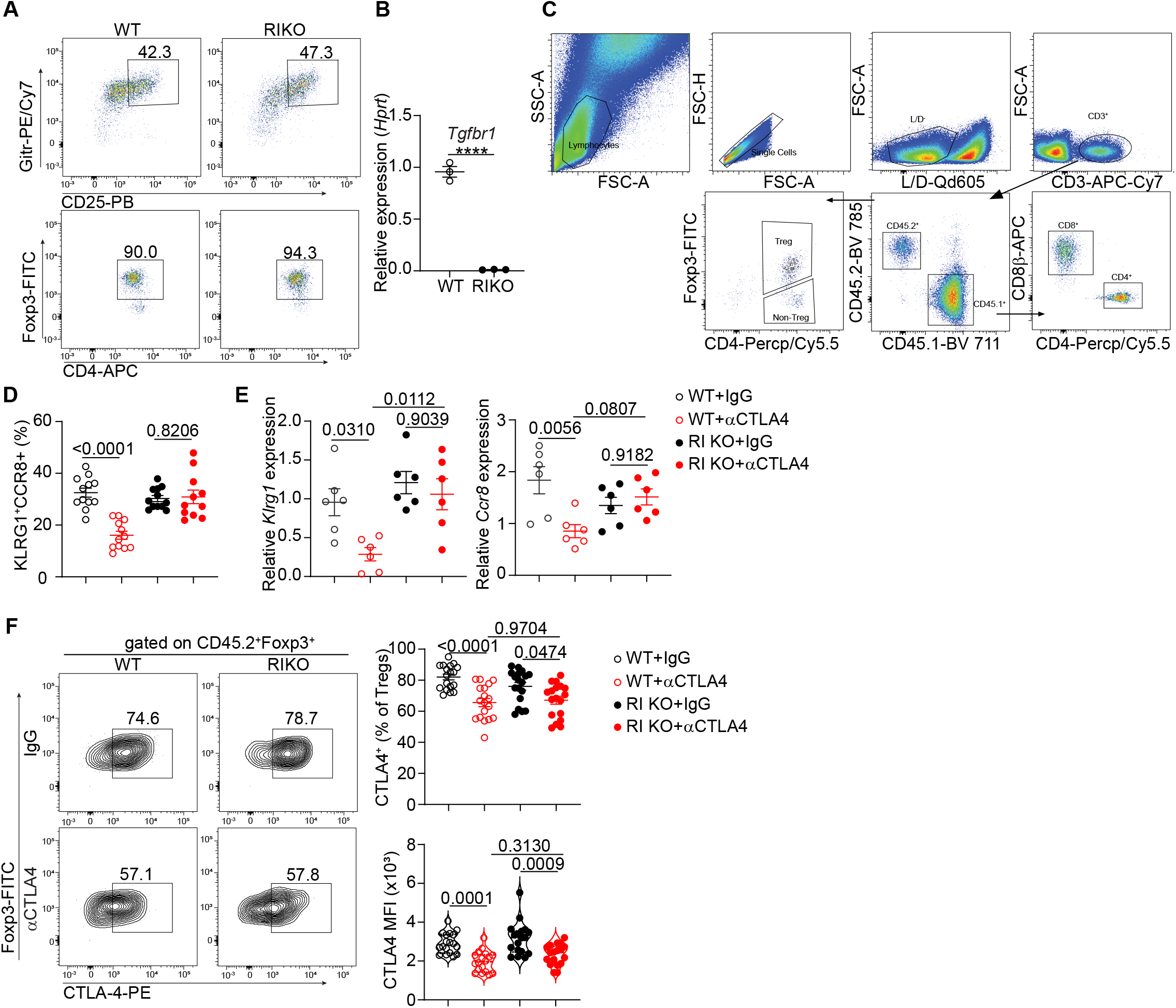
Flow cytometry gating strategy and phenotypic analysis from wide type and *Tgfbr1* ko Tregs in an adoptive transfer model treated with αCTLA-4 therapy. Related to Figure 3. (A) Expression of Foxp3 from flow cytometry-sorted Tregs (CD25^high^GITR^+^) from *Tgfbr1^ER^ ^Cre^* treated with oil or tamoxifen for 5 consecutive days. (B) RT-PCR analysis of expression of *Tgfbr1* in sorted Tregs (CD25^high^GITR^+^). (C) Gating strategy used for analysis of tumor-infiltrating lymphocytes from an adoptive transfer model. (D) Flow cytometry analysis of percentage of KLRG1 and CCR8 expression in tumor-infiltrating Tregs form indicated groups. (E) RT-PCR analysis of *Klrg1* and *Ccr8* in tumor-infiltrating Tregs from indicated groups. (F) Left, Representative flow plot of CTLA-4 staining in tumor-infiltrating Tregs and right, analysis of CTLA-4 expression in tumor-infiltrating Tregs from indicated groups. Data in (A, B, E) are representative of three independent experiments with similar results. Data in (D, F) are pooled from three independent experiments with similar results. Significance was determined by one-way analysis of variance (ANOVA) with Tukey’s multiple comparison. Data shown as mean ± s.e.m.

**Figure S8.**
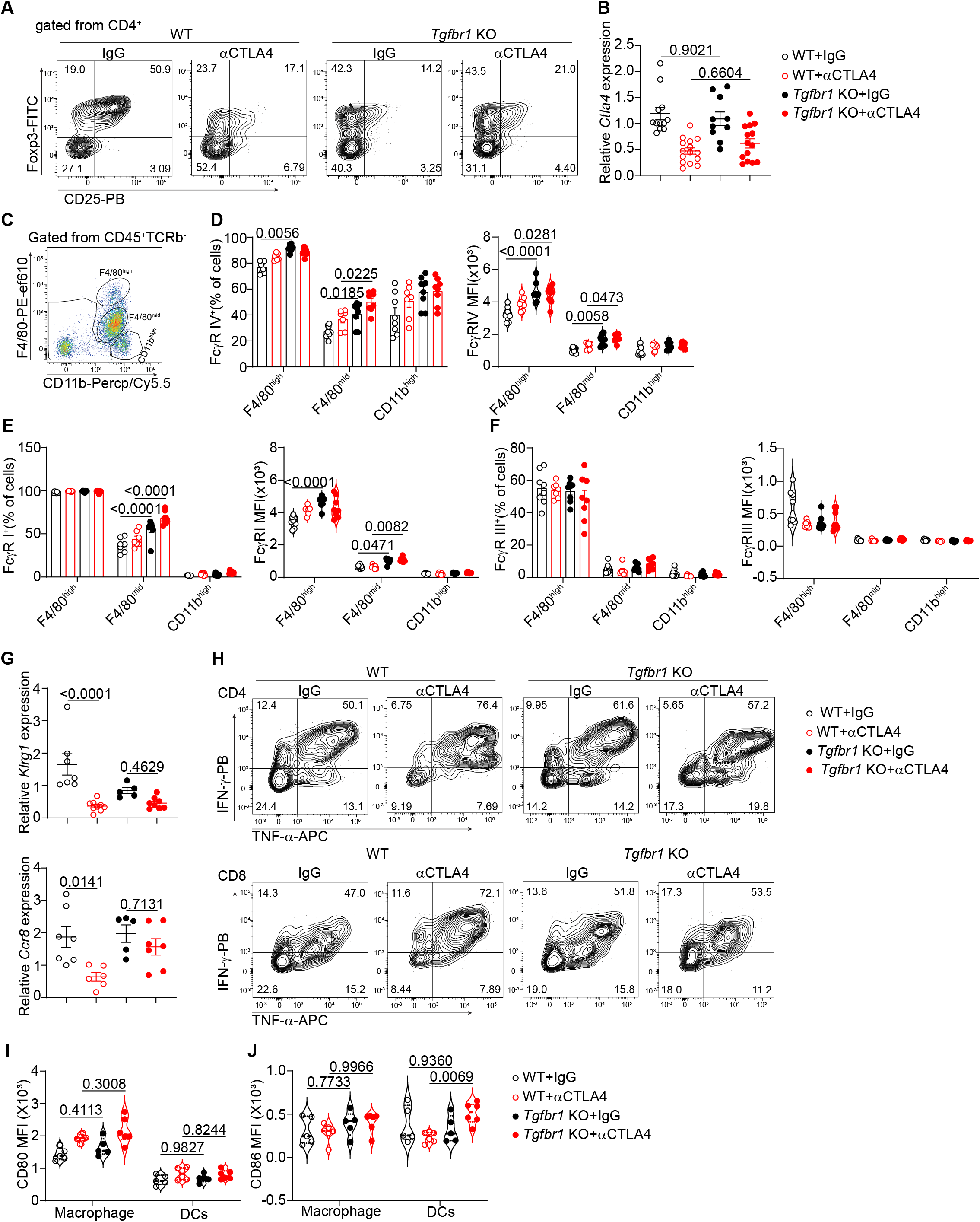
Specific deletion of *Tgfbr1* in Tregs is resistant to αCTLA-4 mediated antitumor therapy. Related to Figure 3. (A) Percentage of Tregs (CD4^+^Foxp3^+^) among CD4^+^ T cells in tumors from WT and *Tgfbr1* ko mice treated with IgG or αCTLA4 antibody. (B) RT-PCR analysis of expression of *Ctla-4* in tumor-infiltrating Tregs from indicated groups. C, Gating strategy used for identification of tumor-infiltrating innate immune cells from tumors. (D-F) Analysis of FcγR IV (D), FcγR I (E) and FcγR Ⅲ (F) expression in intratumor innate immune cells from indicated groups. (G) RT-PCR analysis of expression of *Klrg1* and *Ccr8* in intratumor Tregs from indicated groups. (H) Representative cytometry flow plots of TNF-α^+^IFN-γ^+^ producing CD4^+^Foxp3^−^ or CD8^+^ T cells in tumors from indicated groups. (I-J) Quantification of CD80 (I) and CD86 (J) MFI in macrophages and Dendritic cells from tumors. Data in (A-F, H) are pooled from three independent experiments with similar results. Data in (G, I-J) are representative of two independent experiments with similar results. Significance was determined by one-way analysis of variance (ANOVA) with Tukey’s multiple comparison. Data shown as mean ± s.e.m.

**Figure S9.**
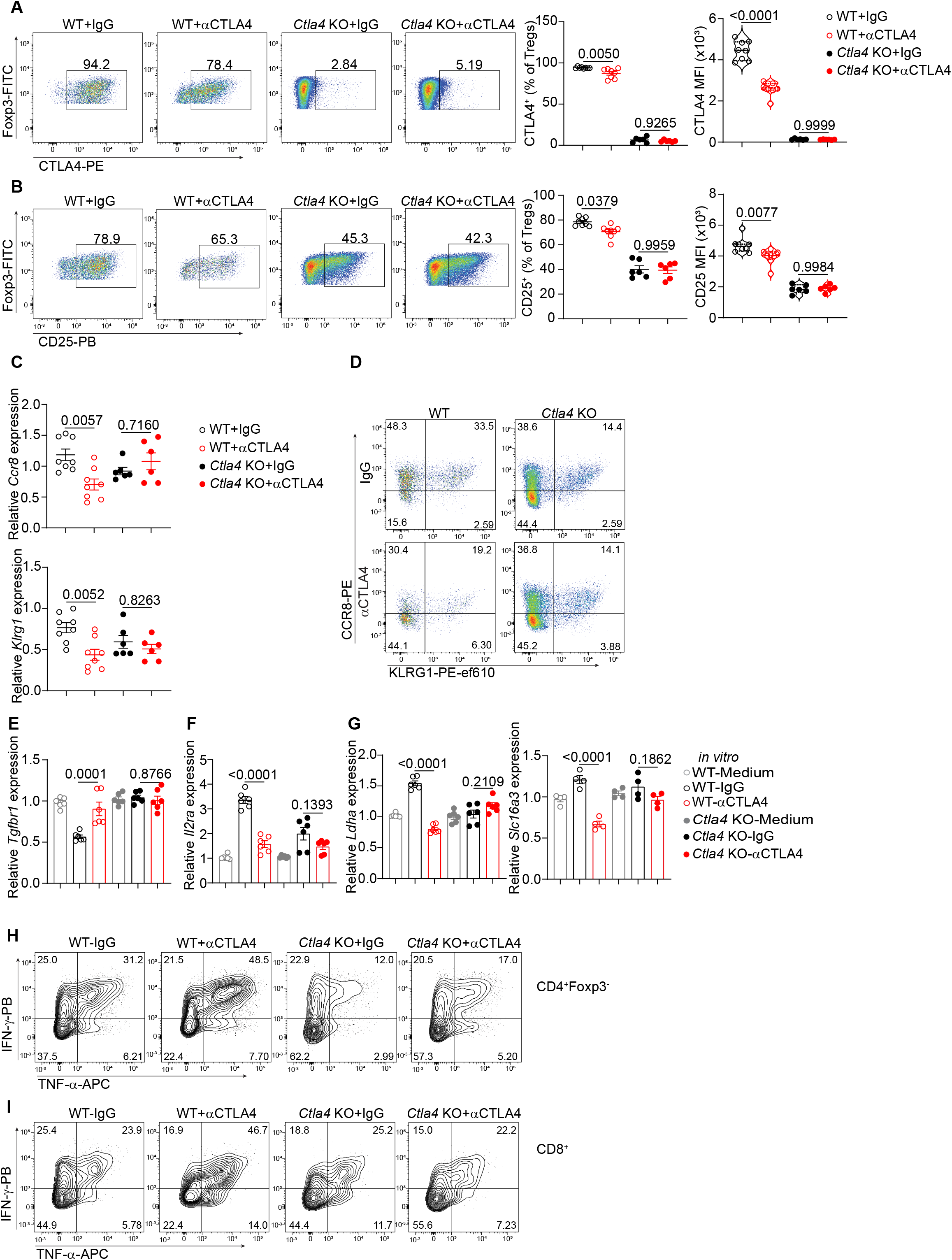
Tregs-specific deletion of CTLA-4 abolishes αCTLA-4 mediated upregulation of TGF-b signaling *in vitro* and anti-tumor immunity *in vivo*. Related to Figure 4. (A) Left, Representative flow plot of CTLA-4 expression in tumor-infiltrating Tregs and right, analysis of CTLA-4 expression in tumor-infiltrating Tregs from indicated groups. (B) Left, Representative flow plot of CD25 expression in tumor-infiltrating Tregs and right, analysis of CD25 expression in tumor-infiltrating Tregs from indicated groups. (C) Expression of *Klrg1* and *Ccr8* in tumor-infiltrating Tregs between IgG vs. αCTLA-4 treatments from WT and *Ctla4* ko mice by RT-PCR. (D) Representative flow plot of KLRG1 and CCR8 expression in tumor-infiltrating Tregs from indicated groups. (E-G) RT-PCR analysis of *Tgfbr1*(E), *Il2ra* (F), *Ldha* and *Slc16a3* (G) in cultured Tregs with IgG or CTLA-4 engagement for overnight from WT and *Ctla4* ko mice *ex vivo*. (H-I) Representative flow plot of TNF-α^+^IFN-γ^+^ producing Th cells or CD8^+^ T cells in tumors from indicated groups. Data is representative of two independent experiments with similar results. Significance was determined by one-way analysis of variance (ANOVA) with Tukey’s multiple comparison. Data shown as mean ± s.e.m.

**Figure S10.**
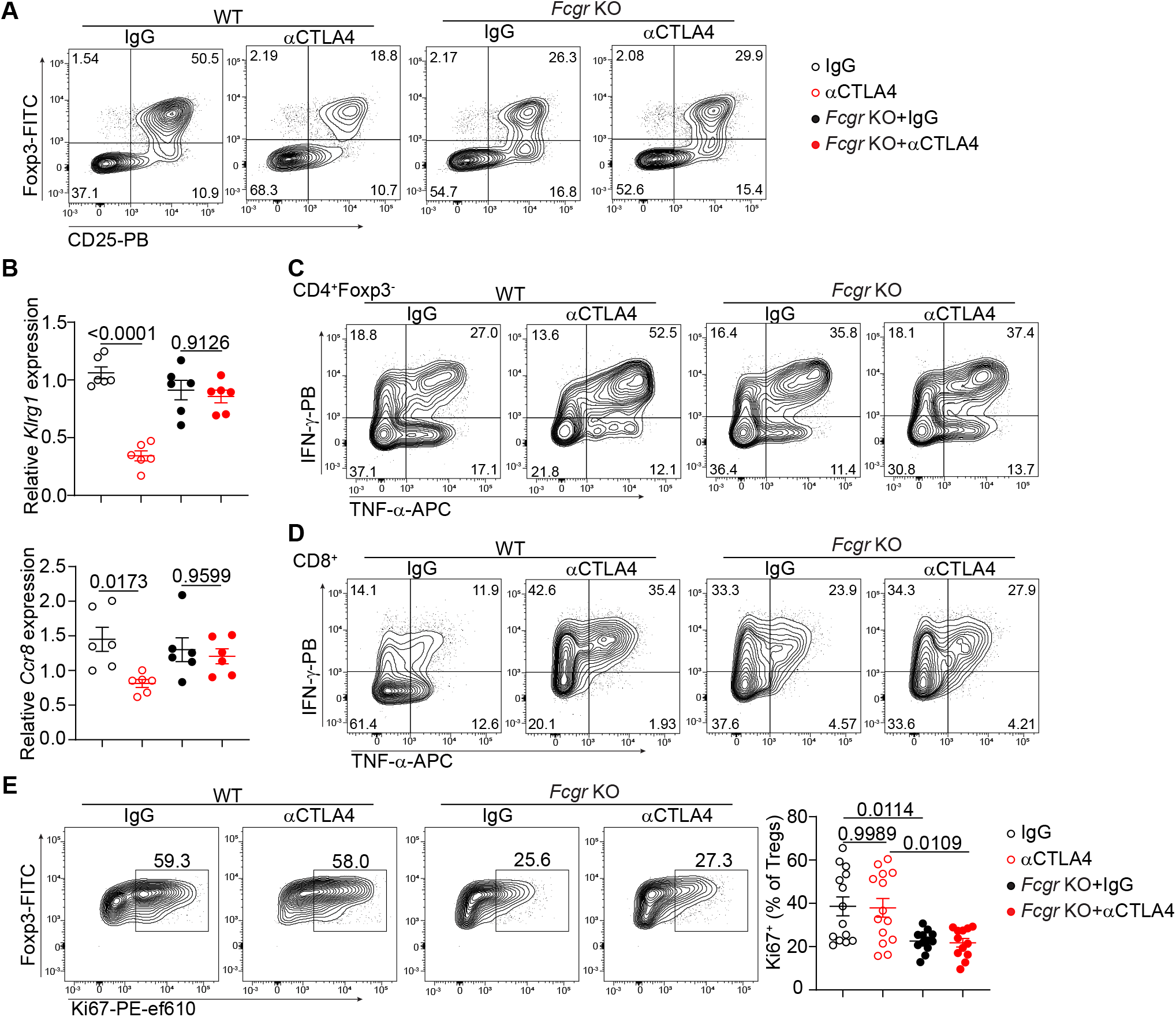
***Fcgr* ko mice exhibits compromised antitumor immunity in response to αCTLA-4 treatment. Related to Figure 4 and 5.** (A) Flow cytometry panels of Foxp3 and CD25 gated from tumor-infiltrating CD4^+^ T cells between IgG vs. αCTLA4 groups. (B) RT-PCR analysis of *Klrg1* and *Ccr8* in tumor-infiltrating Tregs from indicated groups. (C-D) Representative flow plot of TNF-α^+^IFN-γ^+^ producing Th cells (C) or CD8^+^ T cells (D) in tumors from indicated groups. (E) Percentage of Ki67 in tumor-infiltrating Tregs from WT or *Fcgr* ko mice between IgG vs. αCTLA-4 treatment. Data are representative of two independent experiments with similar results (B). Data are pooled from two independent experiments with similar results (A, C-E). Significance was determined by one-way analysis of variance (ANOVA) with Tukey’s multiple comparison. Data shown as mean ± s.e.m.

**Figure S11.**
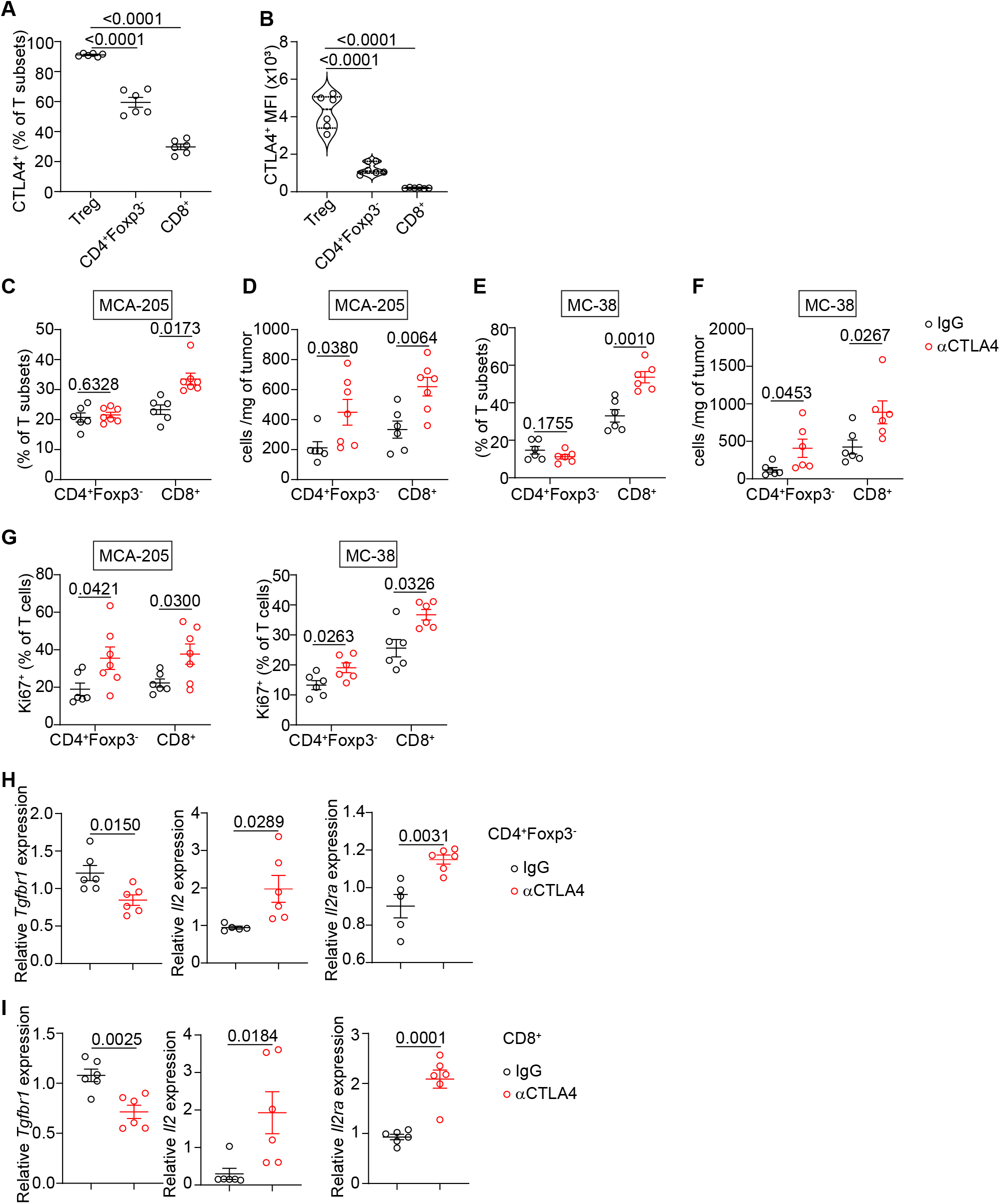
Effects of αCTLA-4 therapy on tumor-infiltrating CD4^+^Foxp3^-^ and CD8^+^ T cells. Related to Figure 5. (A-B) Percentage (A) and quantification (B) of CTLA-4 expression in tumor-infiltrating Tregs, Th and CD8^+^ T cells. (C-D) Percentage of CD4^+^Foxp3^-^ and CD8^+^ T cells among total TCRβ^+^ cells from MCA-205 tumors (C) and Th (CD4^+^Foxp3^-^) and CD8^+^ T cells number per mg of tumor in mice bearing MCA-205 tumors (D) treated with IgG vs. αCTLA-4 antibodies. (E-F) Percentage of Th (CD4^+^Foxp3^-^) and CD8^+^ T cells among total TCRβ^+^ cells from MC-38 tumors (E) and Th (CD4^+^Foxp3^-^) and CD8^+^ T cells number per mg of tumor in mice (F) treated with IgG vs. αCTLA-4 antibodies. (G) Percentage of Ki67 in Th and CD8^+^ T cells from MCA-205 tumors and MC-38 tumors. (H-I) RT-PCR analysis expression of *Tgfbr1*, *Il2* and *Il2ra* in Th and CD8^+^ T cells from MCA-205 tumors treated with IgG vs. αCTLA-4 therapy. Data is representative of two independent experiments with similar results. Significance was determined by unpaired two-tailed *t*-test (C-I) or one-way analysis of variance (ANOVA) with Tukey’s multiple comparison (A, B). Data shown as mean ± s.e.m.

**Figure S12.**
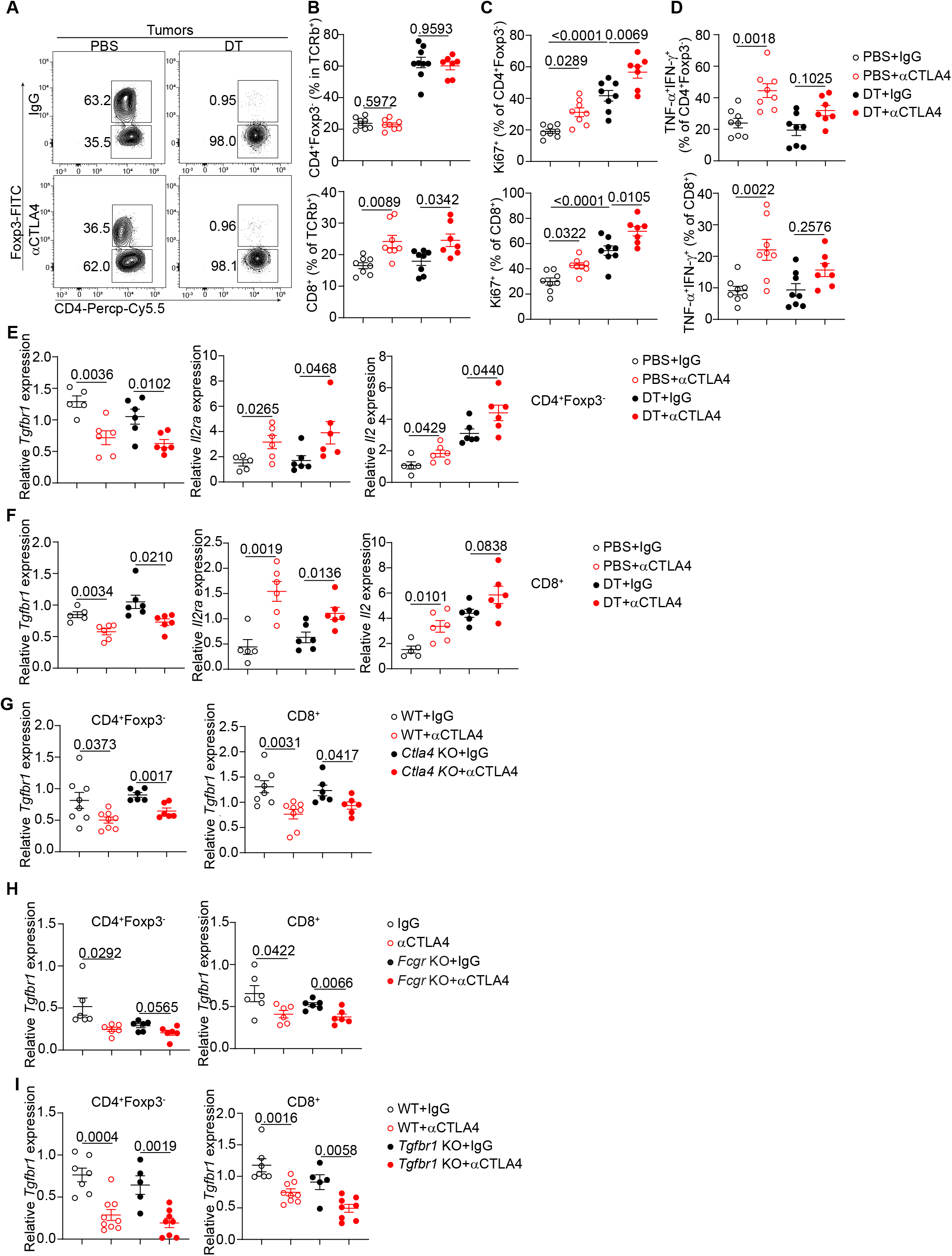
Anti-CTLA-4 therapy blocks CTLA-4 in tumor-infiltrating CD4^+^Foxp3^-^ and CD8^+^ T cells. (A) Representative cytometry flow plots of Foxp3 staining in CD4^+^ T cells from *Foxp3^DT^*^R^ mice bearing MCA-205 tumors. (B) Percentage of CD4^+^Foxp3^-^ or CD8^+^ T cells among total TCRβ^+^ cells in tumors from indicated groups. (C) Percentage of proliferation in CD4^+^Foxp3^-^ and CD8^+^ T cells from tumors. (D) Percentage of TNF-α^+^ IFN-γ^+^ producing CD4^+^Foxp3^-^ or CD8^+^ T cells in tumors from *Foxp3^DTR^* mice treated with IgG vs. αCTLA-4 antibody. (E-F) Expression of *Tgfbr1*, *Il2ra* and *Il2* in CD4^+^Foxp3^-^ (E) or CD8^+^ T cells (F) in tumors from indicated groups detected by RT-PCR. (G) RT-PCR analysis of expression of *Tgfbr1* in CD4^+^Foxp3^-^ or CD8^+^ T cells in tumors from *Ctla4^f/f^Foxp3^cre-ERT^*^2^ mice. (H) Expression of *Tgfbr1* in CD4^+^Foxp3^-^ or CD8^+^ T cells in tumors from *Fcgr* ko mice treated with IgG vs. αCTLA-4 antibodies. I, Expression of *Tgfbr1* in CD4^+^Foxp3^-^ or CD8^+^ T cells in tumors from *Tgfbr1^f/f^Foxp3^cre-ERT^*^2^ mice analyzed by RT-PCR. Data are representative of two independent experiments with similar results. Significance was determined by one-way analysis of variance (ANOVA) with Tukey’s multiple comparison (B-D) or unpaired two-tailed *t*-test (E-I). Data shown as mean ± s.e.m.

**Figure S13.**
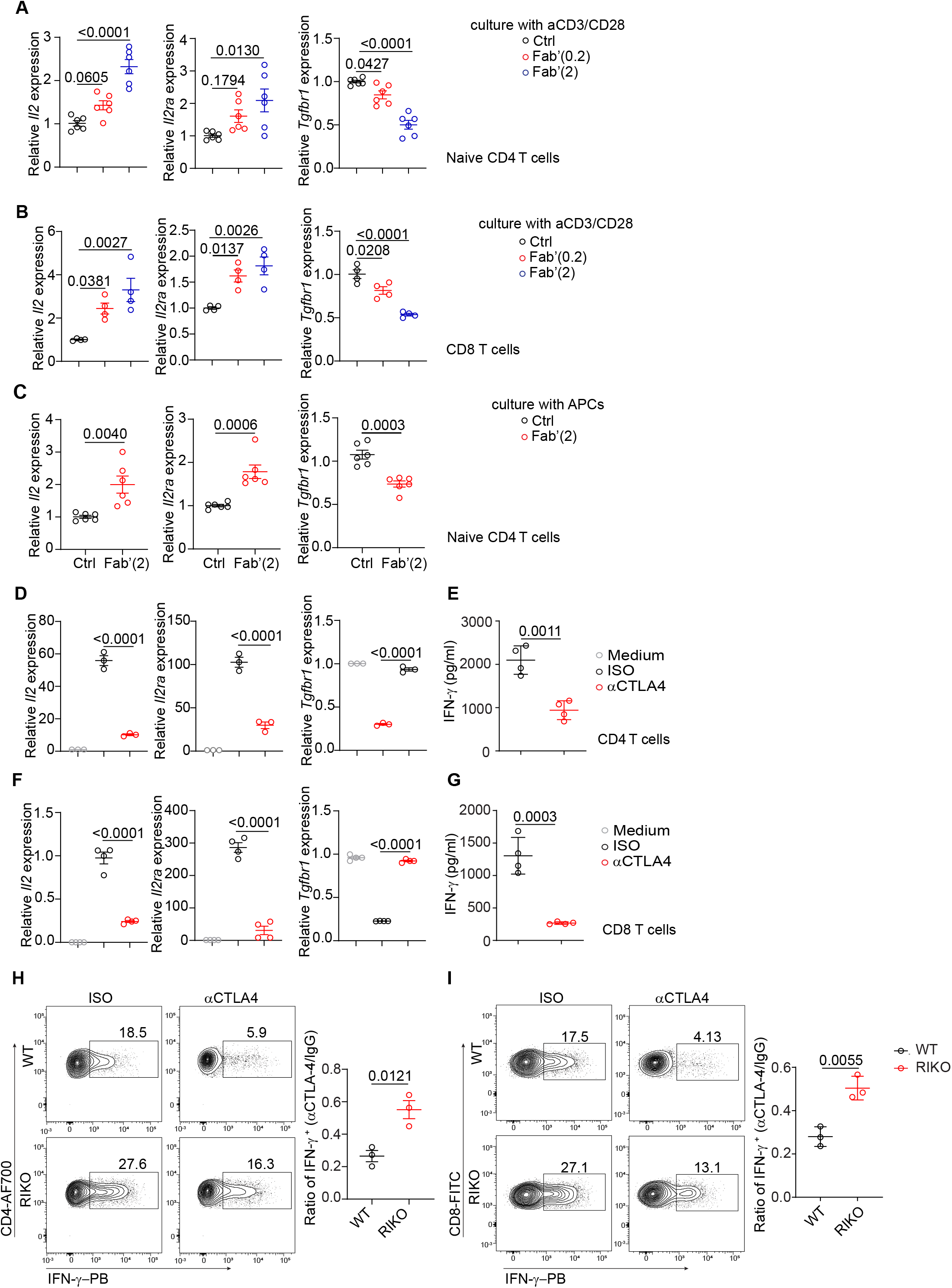
Blockade of CTLA-4 downregulates TGF-β signaling in Naïve CD4^+^ and CD8^+^ T cells, and inhibition of TGF-β signaling compromises IFN-γ suppression induced by CTLA-4 engagement in vitro. Related to Figure 5. (A-B) Expression of *Il2*, *Il2ra* and *Tgfbr1* mRNA in Th cells (A) or CD8^+^ T cells (B) cultured under low dose of αCD3 (0.01 µg/ml) with different does of Fab’ antibody overnight. (C) RT-PCR analysis of *Il2*, *Il2ra* and *Tgfbr1* expression in Th cells cultured with irradiated APC and αCD3 (0.01 ug/ml) in the presence with Fab’ overnight. (D) Expression of *Il2*, *Il2ra* and *Tgfbr1* mRNA in Th cells cultured under CTLA-4 engagement condition. (E) Production of IFN-γ in Th cells cultured under CTLA-4 engagement condition, detected by ELISA. (F) Expression of *Il2*, *Il2ra* and *Tgfbr1* mRNA in CD8^+^ T cells cultured under CTLA-4 engagement condition. (G) Production of IFN-γ in CD8^+^ T cells cultured under CTLA-4 engagement conditions, detected by ELISA. (H-I) Representative flow plot of IFN-γ secretion in Th cells (H) and CD8+ T cells (I) in presence of CTLA-4 engagement from WT or *Tgfbr1* KO mice. Data is pooled from three independent experiments with similar results. Significance was determined by unpaired two-tailed *t*-test (C, E, G, H-I) or one-way analysis of variance (ANOVA) with Tukey’s multiple comparison (A, B, D, F). Data shown as mean ± s.e.m.

**Figure S14:**
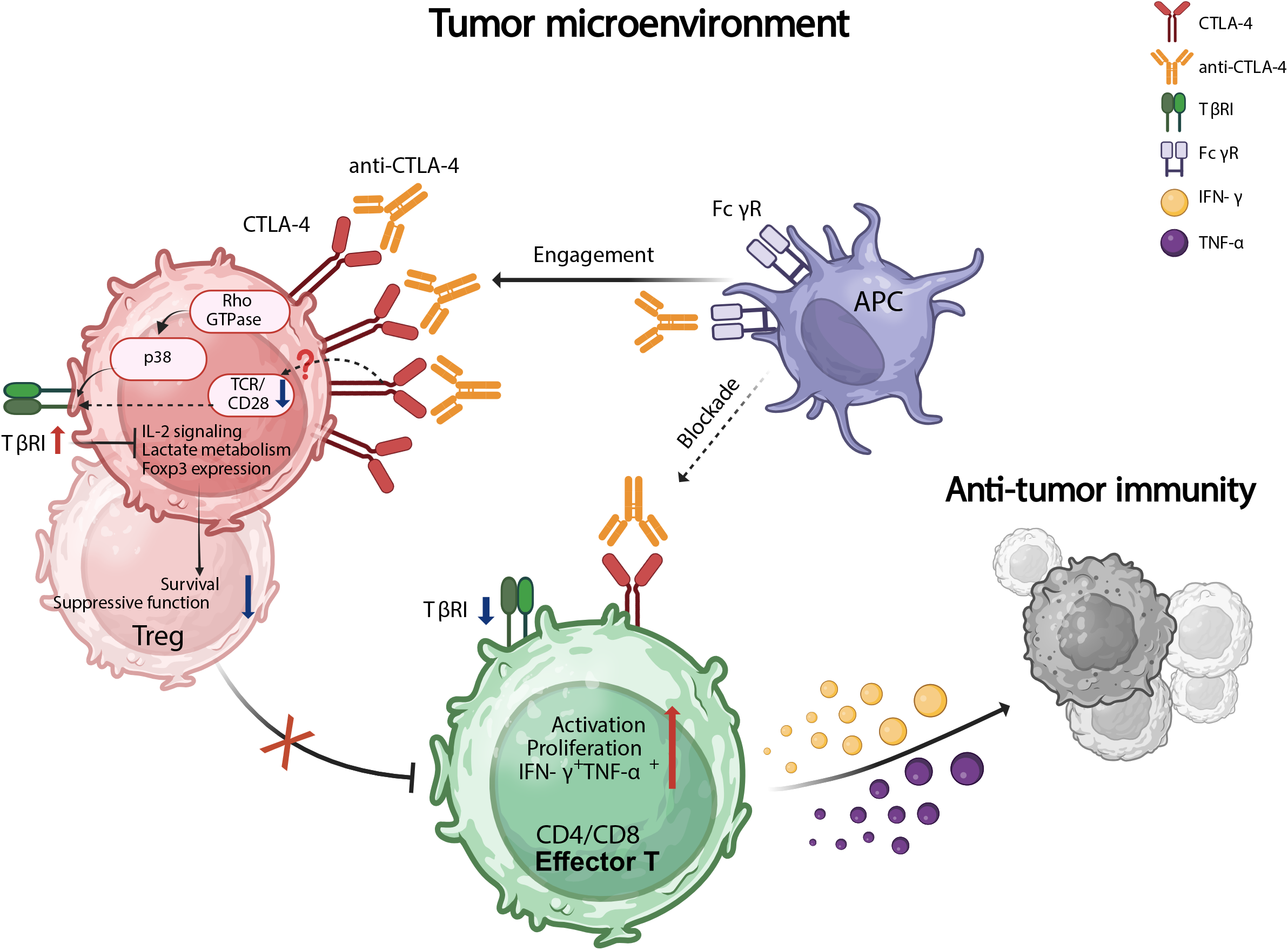
Destabilization of intratumor Tregs by engagement and signaling of CTLA-4 with therapeutic anti-CTLA-4 antibody contributes to the antibody-mediated immunotherapy for cancer. In the tumor microenvironment, therapeutic anti-CTLA-4 antibodies, through the crosslinking and binding of Fc gamma receptors (FcγR) on antigen presenting cells (and/or other FcγR positive cells in the tumor tissues), actively engage the *extremely* high levels of CTLA-4 in the intratumor Tregs to signal through Rho-GTPase-p38 pathway to upregulate TGF-β receptors (e.g. TβRI, TGF-β receptor I), which enhances TGF-β signaling in Tregs in the tumor. This activity suppresses IL-2 signaling (downregulation of CD25) and lactate metabolism to reduce the stability and survival of Tregs, and also inhibits the expression of Foxp3 and other functional associated molecules in Tregs to compromise their suppressive activity. The reduced number and compromised function of Tregs together release the “brake” for CD4^+^Foxp3^-^ and CD8^+^ T effector cells to consequently increase their activation, proliferation and IFNγ^+^TNFα^+^ T effector cells in the tumor. This accounts for an important mechanism for the antitumor immunotherapy by the anti-CTLA-4 antibody. In contrast to the intratumor Tregs, the same anti-CTLA4 antibody may paradoxically block the CTLA-4 in CD4^+^ Foxp3^-^ and CD8^+^ T cells in the tumor tissues. This blockade of CTLA-4 could intrinsically enhance the activation and proliferation of these effector T cells. However, this CTLA-4 blockade-mediated intrinsic activation of T effector cells itself plays minimal, if any, role in anti-CTLA-4-mediated immunotherapy for cancer. Although the exact mechanism by which the same anti-CTLA-4 antibody blocks rather than engages CTLA-4 in these T effector cells remains unknown, it is likely due to the significantly lower levels of CTLA-4 expression in these non-Treg T cells compared to Tregs in the same tumor tissues. The limited amounts of CTLA-4 molecules on the T effector cells restrict their ability to receive the signal from the anti-CTLA-4 antibody, and thus they are unable to reach the minimal threshold to stimulate the signaling cascade through CTLA-4. Instead, the CTLA-4 signaling is blocked. Importantly, this mechanism occurs not only in mouse tumor models, but also in humanized mice with tumor in response to antihuman CTLA-4 antibody treatment. The engagement of CTLA-4 by anti-human CTLA-4 antibody (ipilimumab) also upregulates *TGFBR1* but decreases *Il2RA* expression and lactate metabolism in human Tregs *in vitro* and in humanized mice *in vivo*, leading to suppression of tumor progression.

